# Glassy dynamics in active epithelia emerge from an interplay of mechanochemical feedback and crowding

**DOI:** 10.1101/2025.11.08.687351

**Authors:** Sindhu Muthukrishnan, Phanindra Dewan, Tanishq Tejaswi, Michelle B Sebastian, Tanya Chhabra, Soumyadeep Mondal, Soumitra Kolya, Sumantra Sarkar, Medhavi Vishwakarma

## Abstract

Glassy dynamics in dense epithelial cells remain a subject of debate, as theory predicts its emergence only at unrealistically low cellular activity, yet experimental studies have shown glassy dynamics at physiologically active conditions. In this study, we address this paradox by integrating experimental observations in epithelial monolayers with an active vertex model. We demonstrate that while crowding is essential, it is not sufficient for glassy dynamics to emerge. A mechanochemical feedback loop (MCFL-1), mediated by cell shape changes through the contractile actomyosin network is required to drive glass transition in dense epithelial tissues. Such mechanochemical feedback is captured experimentally via a crosstalk between actin-based cell clustering and dynamic heterogeneity, as well as via force induced actin reorganization in epithelial cells. Incorporating MCFL into the vertex model reveals contrasting results from those previously predicted by theories-we show that the MCFL can counteract cell division-induced fluidisation and enable glassy dynamics to emerge through active cell-to-cell communication. Furthermore, our analysis reveal, for the first time, the existence of novel collective mechanochemical oscillations that arise from the crosstalk of MCFL-1 with oscillatory MCFL-2, capturing ERK mediated cell shape changes. Together, we demonstrate that an interplay between crowding and active mechanochemical feedback enables the emergence of glass-like traits and collective biochemical oscillations in epithelial tissues with active cell-to-cell contacts.

## Introduction

Glass transition is a kinetic phenomenon in which molecular motion slows dramatically as temperature decreases or density increases ^1–3^. A tell-tale feature of glassy dynamics, in addition to jamming, is the coexistence of relatively fast-moving locally fluidised regions and slow-moving arrested regions, or dynamic heterogeneity^3–5^. Similar behaviour is also observed in crowded granular matter and in dense systems of self-propelled, active particles-when density increases, or activity diminishes, a globally arrested state is reached, with the existence of locally fluidised individuals^6–8^. For instance, epithelia-a dense and active tissue-demonstrate glassy dynamics in 2D cell monolayers^9–12^, in organoid cultures mimicking organ architecture^13,14^, and in vivo^15,16^, significant to a plethora of pathophysiological processes from development^15–17^ and cancer^18–21^ to wound healing^22^. In such active systems, dynamic heterogeneity is believed to be dictated by a competition between crowding and internal activity, such that crowding promotes jamming, and activity promotes fluidization^8,23,24^. In fact, theoretical studies modelling epithelial cells, such as the canonical vertex models, predict glassy behaviour only under unrealistic conditions of negligible cell activity and extremely high cell density^25–27^. These predictions, however, contrast with experiments that clearly show glass-like dynamics in crowded, yet metabolically active and proliferating, epithelial monolayers. Such discrepancies suggest that crowding alone cannot explain glassy dynamics in epithelia.

Here, we identified two problems: First, if crowding alone is insufficient, what other factors drive the glass transition in epithelial tissues? And second, how can we incorporate these factors into a model so that biologically plausible features emerge from simulations?

Using actin labelled epithelial cells (LifeAct MDCK), and time-lapse imaging, we mapped actin cytoskeletal rearrangements when cells crowd and dynamic heterogeneity clusters establish, and found that mechanochemical feedback between cytoskeletal dynamics and cell mechanics drives glass transition. Incorporating this feedback into an active vertex model, linking cell geometry to actomyosin signalling, we show that dynamic heterogeneity can emerge even in highly active, dividing tissues, if the mechanochemical feedback is active (Fig. 1). Our experiments and simulations also revealed collective hour-scale actin oscillations, distinct from the minute-scale oscillations observed in isolated cells^28–30^. We show that these slower oscillations arise from mechanochemical feedback that stabilises emerging clusters and disappear when feedback strength or cell density decreases. We show that these oscillations emerge from the interaction between load-dependent actomyosin binding and a second feedback loop coupling the actin cytoskeleton to ERK signalling. Together, our results demonstrate that active mechanochemical feedback enables glass-like dynamics in epithelial tissues through the interplay of cell–cell interactions, signalling, and crowding (Fig.1).

**Figure 1:**
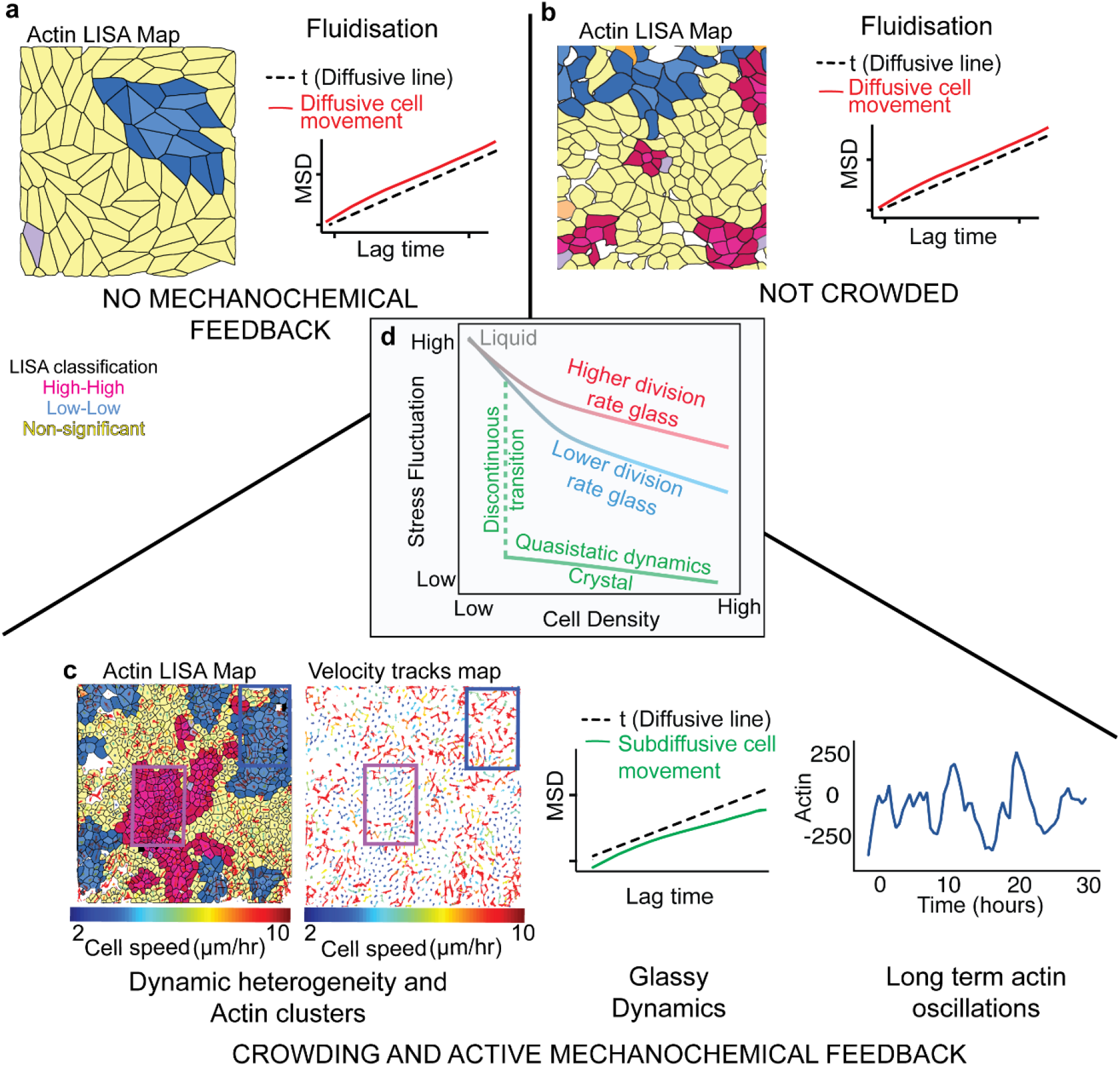
Crowding and Mechanochemical Feedback are necessary for the emergence of glassy dynamics, collective oscillations and spatial clustering of actin in epithelia: a) In the absence of mechanochemical feedback, crowding of the tissue through cell division leads to fluidisation, as revealed by the Left-highly anisotropic shapes from the 1/Area LISA (Local Indicators of Spatial Association) map from our vertex model simulation, where 1/Area is proportional to Actin levels and the Right-superdiffusive mean squared displacement (MSD) of the cells from our vertex model and previous studies17-22. LISA clusters highlight groups of neighbouring cells with similar Actin levels (High-High: Pink and Low-Low: Blue), and cells that are not correlated with their neighbors are classified as Non-Significant (Yellow). b) Mechanochemical feedback without crowding does not lead to glassy dynamics: Right-the MSD remains superdiffusive, and Left-the cells do not form well-defined Actin clusters. c) Crowding and mechanochemical feedback together lead to glassy dynamics. We observe coexisting jammed and unjammed clusters of cells with different motilities, and the MSD becomes subdiffusive revealing dynamic heterogeneity and caging. Furthermore, the cells oscillate collectively with hour-scale time periods. d) The observed glass transition is driven by cell density. Akin to temperature-driven passive glass transition, if the cell density is changed slowly, the tissue undergoes a discontinuous fluid-solid phase transition (green). However, if the density is increased sufficiently rapidly, such as through cell division, the tissue can avoid the discontinuous transition and undergo a glass transition (red and blue) at a density that depends on the division rate: a higher division rate leads to a faster glass transition (red).

## 2. Results

### 2.1 Interplay of crowding and mechanochemical feedback dictates emergence of glassy dynamics

We first investigated any possible coupling between glassy dynamics and actin expression in the epithelial monolayer at homeostasis (Fig. 2a). We compared velocity fields with spatial actin distribution patterns from time-lapse imaging data of MDCK cells tagged for Actin (Supplementary video 1 and Fig. 2b). Spatial organization of F-actin was computed using local Moran’s index, thus allowing us to generate Local Indicators of Spatial Association (LISA)^31^ maps (Fig. 2a, c, Supplementary Fig. 1a-f), which classified cells into clusters representing regions of spatially correlated F-actin levels. Velocity tracks (Fig. 2d) revealed coexisting slow and fast-moving clusters at a densely packed state (Fig. 2c, d), indicating dynamic heterogeneity, a hallmark of glassy systems^4,5,32^. Additional features of glassy dynamics were also observed, including sub-diffusive mean squared displacement and caging (Fig. 2e), and a sub-exponential decay in the overlap function (Q(t), Fig. 2f). To capture the link between the dynamic clusters arising from glassy behavior and the biochemical organization of cells, we quantified the spatial F-actin organization using LISA and found ‘High-High’ & ‘Low-Low’ actin clusters, where neighbors correlated in actin expression. We termed those as hotspots and coldspots, respectively. Interestingly, slow-moving, jammed regions aligned with actin hotspots, whereas fast-moving, unjammed regions corresponded to actin coldspots (Fig. 2c, g), indicating that glassy dynamics and actomyosin organization are linked. From traction force microscopy experiments (Fig. 2h-k), we found that cellular tractions and actin expression are negatively correlated; specifically, unjammed coldspots displayed higher traction forces, whereas jammed hotspots displayed lower traction forces (Fig. 2i-k), as reported previously^10,33,34^.

**Fig 2:**
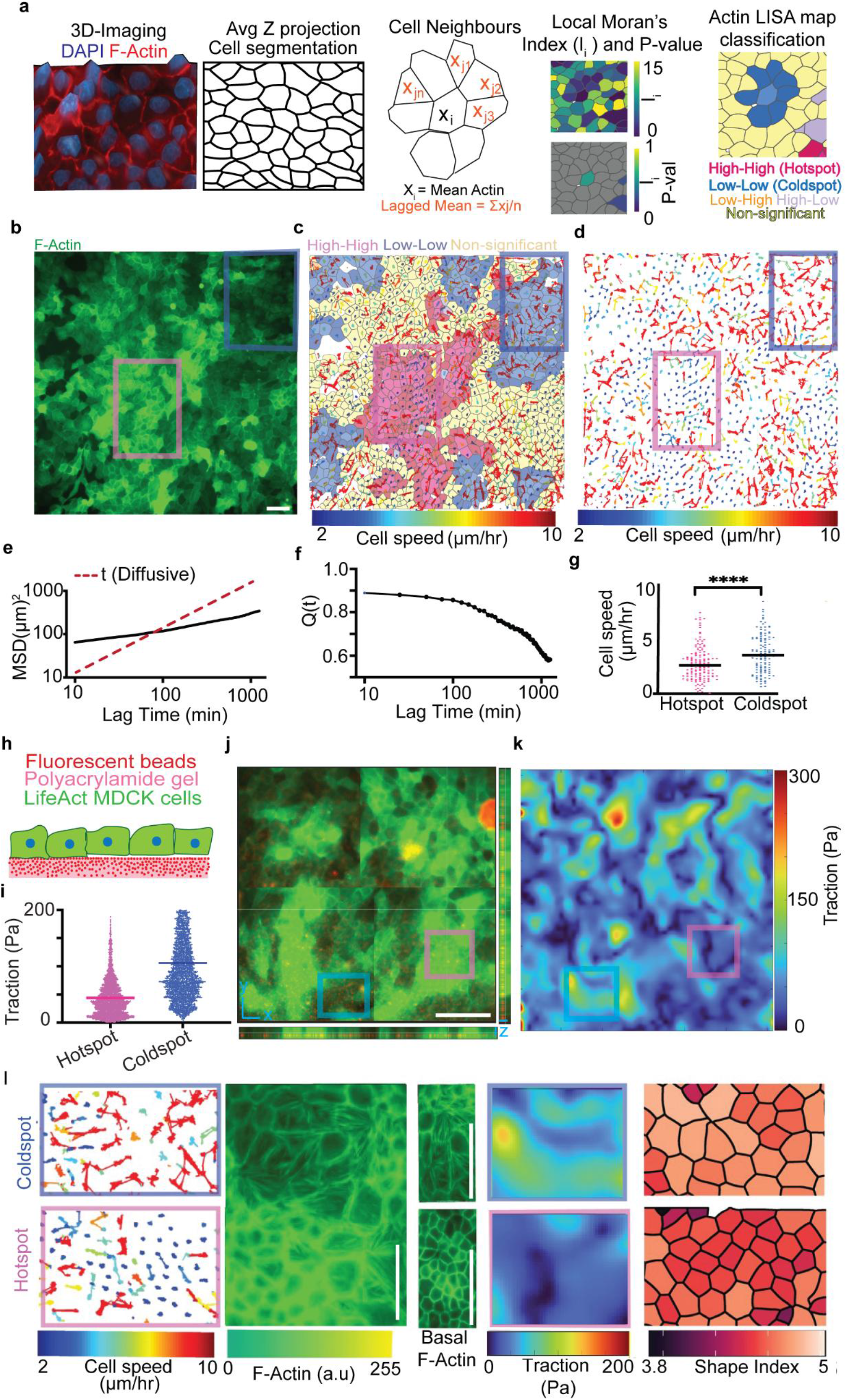
Spatial distribution of Actin and forces follows the dynamic heterogeneity landscape: a)Schematic of Moran’s index analysis with cellpose segmentation and Moran analysis using custom R code, b) Representative image from timelapse microscopy of LifeAct-MDCK cells, c) Cell tracks overlaid on F-Actin LISA map, d) Cell trajectories obtained from trackmate, e) Mean Square Displacement (MSD) plot for one representative timelapse movie of cells, f) Self-overlap function (Q(t)) plot for one representative timelapse movie of cells, g) Actin levels in groups of fast and slow cells and in non-correlated velocity regions, h) Schematic of traction force microscopy, i) Representative image of cells (green for Actin) and fluorescent beads (red), j) Traction magnitude at Actin hotspots and coldspots, k) Left-Traction heatmap (top) at low Actin (bottom) regions and right-Traction heatmap (top) at high Actin regions (bottom). Unpaired t-test: *p < 0.05, **p < 0.01, ***p < 0.001, ****p < 0.0001. Scale bars= 50μm.

While this inverse correlation between actin intensity and both cellular movement and cellular forces may appear non-intuitive, it became clear when actin architecture was closely mapped in these regions. These experiments reveal higher basal stress fibers in unjammed regions, which correspondingly move more, exert higher tractions, and have elongated shapes (Fig. 2l), while jammed regions showed higher cortical actin, explaining higher junctional strength, but lower movement, lower tractions and more compacted shapes (Fig. 2l).

Furthermore, we observed a clear correspondence between actin LISA maps, and other mechano-transducers, such as vinculin, E-cadherin, and phospho-myosin (Supplementary Fig. 2). Together, these results suggest that dynamic heterogeneity manifests itself in biochemical signatures through mechanochemical feedback and directly affects tissue functionality by modulating cells’ internal machinery. To further verify mechanochemical feedback in the system, we examined biochemistry-mediated force changes by measuring traction with and without Blebbistatin (Supplementary Fig. 3) and found that drugs that alter cellular biochemistry also dynamically alter traction (Supplementary Fig. 3e-f). Conversely, applying external force by stretching the cell substrate with a custom cell stretch device changed F-Actin levels, suggesting that external forces can dynamically change cellular biochemistry (Supplementary Fig. 3a-c).

To further ascertain that mechanochemical feedback is an important determinant of glass transition, we inhibited this feedback by contractility inhibitor Y27632, which inhibits ROCK pathway (Supplementary Fig. 4a, d). As expected, contractility inhibition led to reduced stress fibres resulting in uniform actin architectures in the monolayer (Supplementary Fig. 4h). While actin expression still showed clustering as confirmed by the presence of actin hot-spots and cold-spots (Supplementary Fig. 4b, d), the difference in movement in these clusters disappeared (Supplementary Fig. 4c, f). Furthermore, the monolayer was fluidized, with lack of jamming as confirmed by a diffusive MSD (Supplementary Fig. 4g, 4h), suggesting that the disruption of mechanochemical feedback prevents glass transition.

Since crowding is essential for dynamic heterogeneity to emerge in active and passive glass formers, we wondered whether biochemical clustering is also modulated at different levels of crowding. Indeed, LISA maps of F-actin in low and high-density monolayers (Fig. 3a and 3b) showed differences in locally correlated clusters: loosely packed monolayers had smaller clusters, and densely packed monolayers had larger clusters (Fig. 3c). Global Moran’s index, a metric for spatial correlation in actin among immediate neighbours, increased with cell density, while overall spatial actin heterogeneity reduced (Fig. 3d, Supplementary Fig. 5b-e). Furthermore, in loosely packed monolayers, the mean squared displacement was not sub-diffusive (Fig. 3e) and the decay in the overlap function was faster than in a densely packed monolayer (Fig. 3f), suggesting that crowding is essential for the emergence of glassy dynamics. PIV analysis also showed diffusive dynamics in a loosely packed monolayer, but movement consistently became less diffusive with local pockets of unjamming, as density rise (Supplementary Fig. 6f-h).

**Fig 3:**
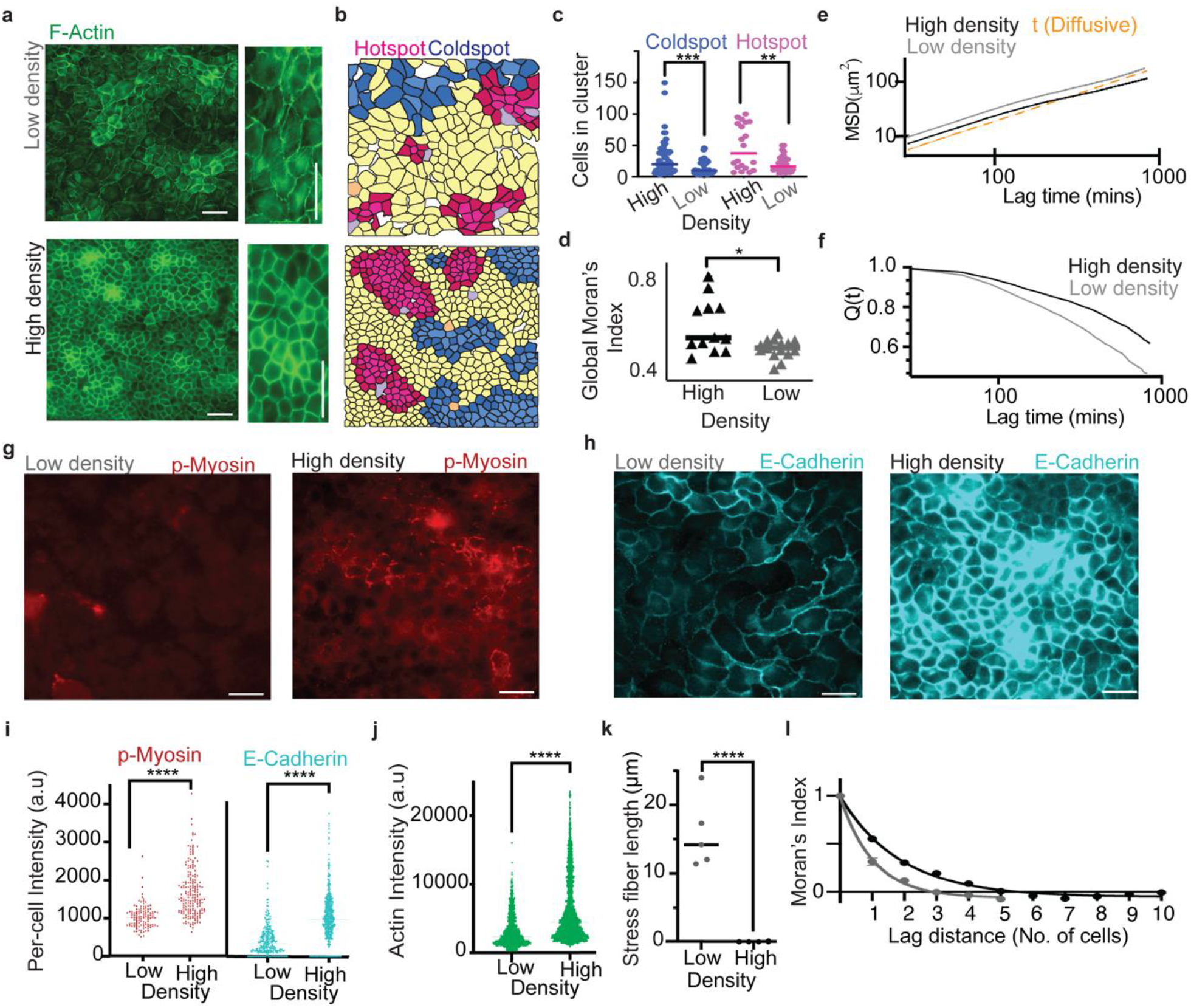
Higher spatial clustering with increasing density: a) Immunocytochemistry images of Actin at low (top) and high (bottom) density, b) LISA cluster maps of Actin at low (top) and high (bottom) density, c) Number of cells in hotspots and coldspots at high and low density, d) Global Moran’s Index at low and high density, e) Mean Square Displacement (MSD) plots for Low density (Gray) and High density (black) monolayers, f) Self Overlap function Q(t) plots for Low density (Gray) and High density (black) monolayers, g) Immunocytochemistry average Z-projected image of Phospho-Myosin (P-Myosin) at low (left) and high (right) density, h) Immunocytochemistry average Z-projected image of E-Cadherin at low (left) and high (right) density, i) Single-cell intensity comparisons of E-Cadherin (blue) at high and low density and P-Myosin (red), j) Comparison of single-cell Actin intensity at high and low density, k) Stress fiber length at high and low density, l) Correlogram showing the decay in Moran’s Index as a function of number of lags. Scale bars=50μm. *p=0.017.

We also observed global differences in cell adhesions and contractility, as indicated by the overall lower intensity of actin, E cadherin as well as myosin in low-density monolayers (Fig. 3a, 3g-j). Longer and more numerous actin stress fibres in low-density monolayers (Fig. 3a, 3k) explains unjamming and higher movement of cells. In densely packed monolayers, cortical actin was higher, stress fibres were shorter and less numerous, and overall actin intensity was higher (Fig. 3a, 3j, 3k). Finally, inhibiting cell division by mitomycin in a homeostatic epithelium inhibits crowding or jamming (Supplementary Fig. 6a-c) and prevents glass transition, as confirmed by cell shapes and MSD plots, such that cells remain unjammed with diffusive MSD profile in the presence of mitomycin (Supplementary Fig.

6d). Together, these results indicate an active interplay between crowding and mechanochemical feedback in dictating cell clustering and glassy behavior at homeostasis.

### 2.2 An active vertex model with mechanochemical feedback and crowding undergoes a glass transition

#### 2.2.1 Actomyosin contractility drives two coupled mechanochemical feedback loops

To delineate the role of crowding and mechanochemical feedback, we developed an active vertex model that integrates cell division and mechanochemical feedback loops (MCFLs). Specifically, we considered the crosstalk of two MCFLs (Fig. 4a). The first, MCFL-I, arises from load-dependent binding of myosin to actin^35–37^, whereas the second, MCFL-II, arises from the feedback between ERK (extracellular signal-regulated kinase) and the actin cytoskeleton^38–40^ (Fig. 4a). MCFL-I is sufficient to understand the glass transition, but the crosstalk of the two is needed to understand the collective oscillation that we describe later.

**Figure 4:**
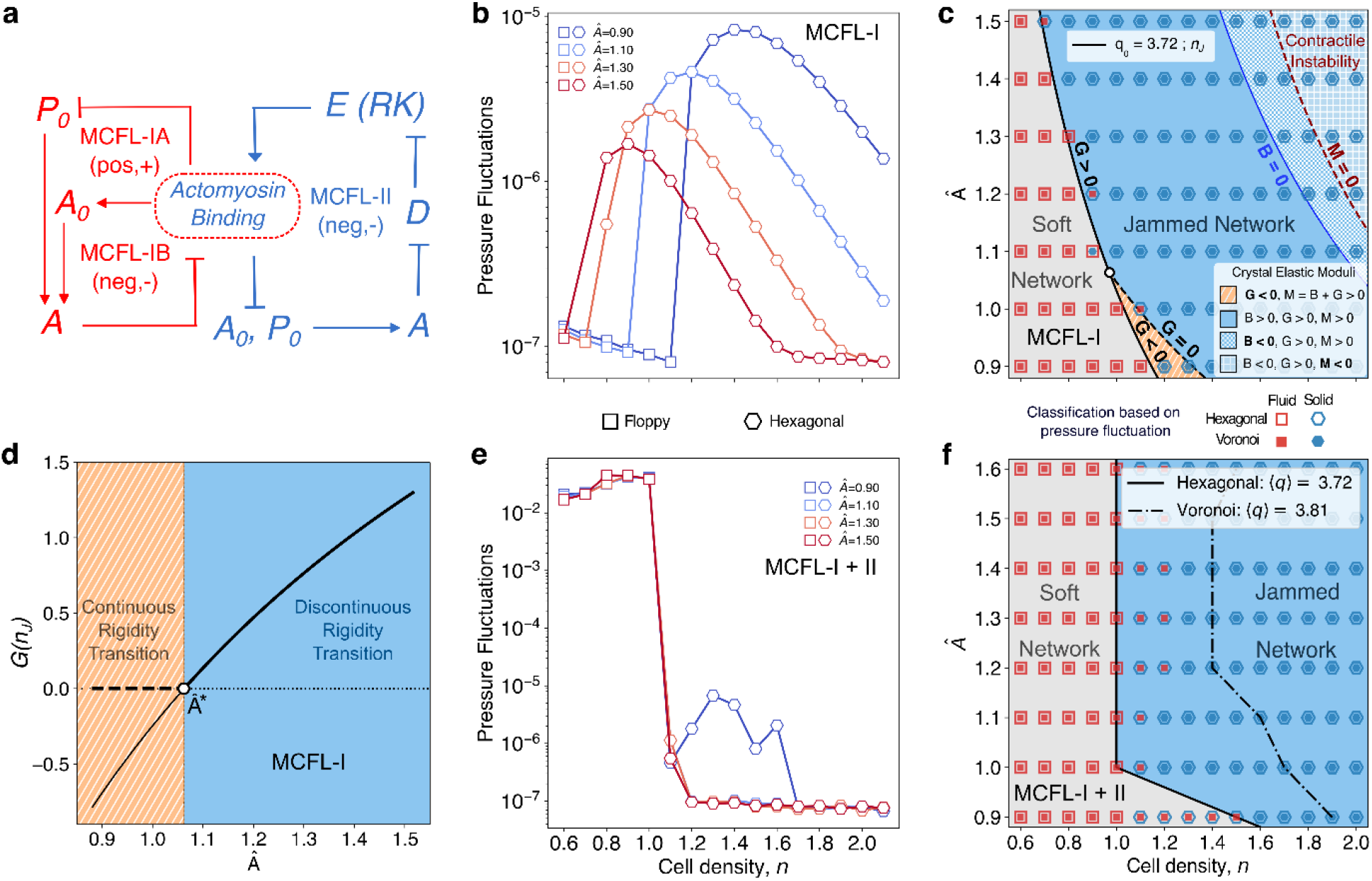
Ground states of active vertex model with mechanochemical feedback: (a) A schematic showing the crosstalk of MCFL-I and MCFL-II. (b) In constant density simulations (no cell division), pressure fluctuations change discontinuously at a density 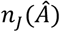. This discontinuous change marks the boundary between soft fluid network to jammed network. Here results from the hexagonal initial condition are shown. (c) Using pressure fluctuation, the states can be classified into fluid (red square) and solid (blue hexagons) phases. Open and filled markers denote hexagonal and Voronoi initial conditions, respectively, which shows that the transition is agnostic to the initial condition. The solid black line corresponds to 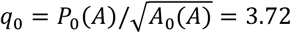 and clearly demarcates the fluid and the solid phases. However, the location of this line is not density independent because of the mechanochemical feedback. Above n_J_, the ground state is a hexagonal crystal for the hexagonal initial condition. The elastic moduli computed in this state can be used to classify the crystalline ground state. Specifically, it shows that at sufficiently high density, the feedback generates spatially heterogeneous patterns through contractile instability. (d) Also, above a threshold feedback strength, 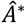 (black open circle), the shear modulus, G(n_J_), is positive, implying a discontinuous transition. Below this threshold, it is negative, implying that the transition is continuous. (e, f) We obtain results similar to (b, c) when both MCFL-I and II are considered.

In the model, the load-dependent binding is captured by cell area (*A*) dependent variation of the vertex model parameters *A*_0_ and *P*_0_ (SI-theory), which we model using a key experimental observation. We observe that as the tissue matures, F-actin and myosin localise near the cell-cell junctions at the expense of the stress fibres (Fig. 3a, k), suggesting an increase in line tension, but reduced contractility as *A* decreases upon crowding.

In the vertex model, with Γ and *K* as the perimeter and area elasticity coefficients, respectively, contractility is equal to 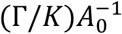 and tension is equal to 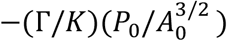^41,42^ (SI-theory). Hence, crowding, i.e. decreasing *A*, must increase *A*_0_ and decrease *P*_0_ (SI-theory, Fig. 4a). Crucially, this choice renders *A*_0_ and *P*_0_ as dynamical variables, which changes with respect to time, captured here by the following two equations:

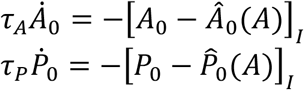

Here, 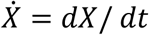 and the subscript *I* indicates that the terms originate from MCFL-I. At a constant area *A, A*_0_ and *P*_0_ relax to *A*_0_ = 2*a*_0_*p*_*bound*_ and 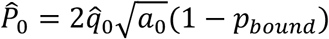, with timescales *τ*_*A*_ and *τ*_*P*_, where 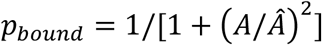 is the fraction of cortical actomyosin and 1 − *p*_*bound*_ is the fraction of stress fibres (SI-theory). The parameter 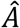 measures the strength of MCFL-I and is inversely proportional to the binding affinity of myosin to actin. The other parameters, *a*_0_ = 1 and 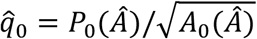, defines the *A*_0_ and the shape index when 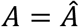, i.e., *p*_*bound*_ = 0.5 (SI-theory). Because the cell motility is driven by the stress fibres, we expect the motility to be directly proportional to 1 − *p*_*bound*_:

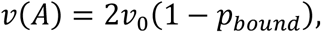

where *v*_0_ is the motility when 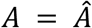. Surprisingly, this simple modification (Supplementary Fig. 9) is sufficient to drive tissue solidification through cell division. In contrast, in the canonical vertex model, where *A*_0_, *P*_0_, and *v* are fixed parameters of the model, cell division leads to tissue fluidisation (Supplementary video S9).

The equations governing MCFL-II are more complicated because they incorporate the ERK as a Hopf oscillator. We model ERK (*E*) as a Brusselator, but any choice of Hopf oscillators would work^40^. The equations describing *E* are:

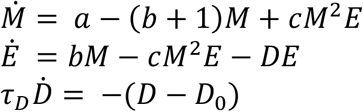

where *M* is a regulator of *E* and *D* is a degrader of *E. a, b, c*, and *D*_0_ are the parameters of the Brusselator (SI-theory). MCFL-II couples these equations with the cell mechanics. Specifically, deviation of *E* from its homeostatic value *E*_0_ changes *A*_0_ and *P*_0_ with strength *α*, whereas compressing the cell below a threshold area of 1 upregulates the production of the degrader *D* with a strength *β*, such that *μ* = *αβ* determines the strength of MCFL-II. Taken together, the equations governing MCFL-I and II are:

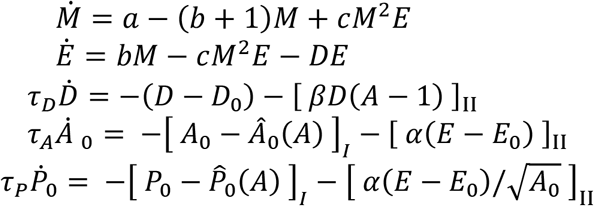

We use this final set of equations to investigate the origin of tissue solidification.

#### 2.2.2 In the quasistatic limit, tissue solidification is density-dependent and occurs through a discontinuous rigidity transition

Is the observed tissue solidification a glass transition? In passive, temperature-driven glass transitions, the glass-forming liquid must be cooled through its freezing point faster than its nucleation rate. By doing so, it can transition to the metastable supercooled liquid phase, which can undergo a glass transition. Otherwise, it discontinuously transitions to the crystalline ground state^43^. It has been argued that cell density, *n*, is the driver of glass transition in tissues, such that the cell division rate is the analogue of the cooling rate^12,15^. If this analogy is exact, then we expect that if *n* is changed quasistatically in the absence of cell division, then tissue solidification must happen through a discontinuous phase transition.

The pioneering works of Staple et.al^44^. and Bi et.al^46^. have shown that in the context of the canonical vertex model, tissue solidification is *density-independent* and is driven only by cell shapes. As the cell shape, characterized by the shape parameter 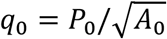, crosses the jamming point 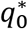, the tissue undergoes a continuous transition from a floppy to a rigid network of cells, marked by the continuous change of the shear moduli *G* from zero to a nonzero value. Considering these results, our hypothesis of density-driven discontinuous rigidity transition is rather surprising.

To test this hypothesis, we disabled cell division in the model, set *v* = 0, and fixed the values of *n* and 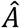 and let the system relax to the ground state. These choices enabled us to turn off any time-dependent variation of *A*_0_ and *P*_0_, while preserving the density-dependent variation, such that our model can be directly compared to the earlier works. First, we turned off MCFL-II and studied the effect of just MCFL-I on the rigidity transition, using discontinuous changes in the pressure fluctuation (SI-theory), as the marker of the transition (Fig. 4b). To understand the dependence of this transition on the feedback strength 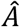 and the cell density *n*, we marked the fluid-like floppy network from the rigid network on a phase diagram (Fig. 4c). Interestingly, the rigidity transition was very similar when either disordered hexagonal lattice (hex-IC) or Voronoi tiling (vor-IC) was used as the initial condition. Consistent with the earlier works, the transition occurred when *q*_0_ (for hex-IC) or, equivalently, 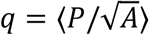 (for vor-IC) was approximately 3.722, the shape index of the hexagonal lattice (Fig. 4c). However, the transition was not independent of *n*. Because *A*_0_ and *P*_0_ are coupled to *n* through *p*_*bound*_ in MCFL-I, *q*_0_ ≈ 3.722 occurs at a density *n*_*J*_ that depends on the feedback strength 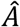. Therefore, our first important conclusion is that although tissue solidification can be understood only through cell shapes, cell shapes and cell density are intimately connected through mechanochemical feedback loops. Hence, tissue rigidity transitions are *not density independent*.

Next, we sought to understand how *G* changes across the transition. To do so, we computed the shear modulus of a hexagonal crystal for 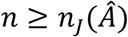, where it is the ground state. Because the rigidity transition was nearly identical between hex- and vor-IC, this approximation was sufficient to delineate the difference in the rigidity transition between the vertex models with and without feedback. Indeed, in our model, the transition at *n*_*J*_ was not accompanied by the vanishing of *G*. Instead, above a threshold 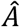, referred to here as 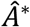, *G*(*n*_*J*_) changed abruptly to a positive value, while it was negative below this threshold (Fig. 4d). The abrupt positive change in *G* above 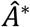 reinforces our observation that the density-driven tissue rigidity transition is a discontinuous transition. *G* < 0 below 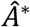 indicates the fragile nature of the crystalline state for densities just above *n*_*J*_. Specifically, these states are unstable to infinitesimal shear^47–49^. Indeed, for 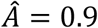, which is below 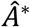, we observe anomalous shape and pressure fluctuations in this range when the tissue was simulated with MCFL-II (Fig. 4e) and motility (SI-theory). Hence, below 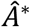, we expect rigid states to emerge only when *G* = 0 through a continuous transition, as in the jamming transitions^44,46,50^.

We also observe a discontinuous change in pressure fluctuations when the crosstalk of MCFL-I and II is considered (Fig. 4e). The transition corresponds to ⟨*q*⟩ ≈ 3.722 for the hex-IC, as in the case with only MCFL-I. However, the density at which this transition happens is different from MCFL-I (Fig. 4f), as MCFL-II adds another coupling of *A*_0_ to the density via the *βD*(*A* − 1) term. The discontinuous transition in the case of vor-IC also happens at densities that depend on 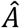, where 3.81 < ⟨*q*⟩ < 3.9. Therefore, our second important conclusion is that when the mechanochemical feedback is sufficiently strong, tissue solidification occurs discontinuously at 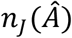. Hence, increasing cell density above *n*_*J*_ with a sufficiently fast division rate should make tissues glassy. This is indeed what we observe.

#### 2.2.3 Increasing cell density at a finite division rate leads to a glass transition

Increasing density through constant division rate simulations at different division rates *r*, we find that at sufficiently low densities, irrespective of the division rates, the pressure fluctuations decay at the same rate as the fluid state in the quasistatic simulations (Fig. 5a, Supplementary Fig. 10), indicating that a unique fluid state exists at these densities. As the density increases, the system exits the fluid phase at a density *n*_*G*_(*r*), that decreases with increasing division rate (triangular markers in Fig. 5a, b). Above *n*_*G*_, the decay in pressure fluctuations or mean asphericity do not depend on 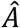, but below this density they vary with 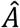. This observation implies that cell-cell interactions are the key drivers of relaxation below *n*_*G*_, whereas above it, crowding is the key driver^51^. This observation parallels the phenomenology of the passive glass transition, where a similar cooling-rate dependence is observed ^1,3^. Furthermore, the switching of the relaxation mechanism is reminiscent of the transition between the mode-coupled and activated relaxation postulated in passive glass transition. Hence, we claim that at constant division rates, the tissue undergoes a glass transition.

**Figure 5:**
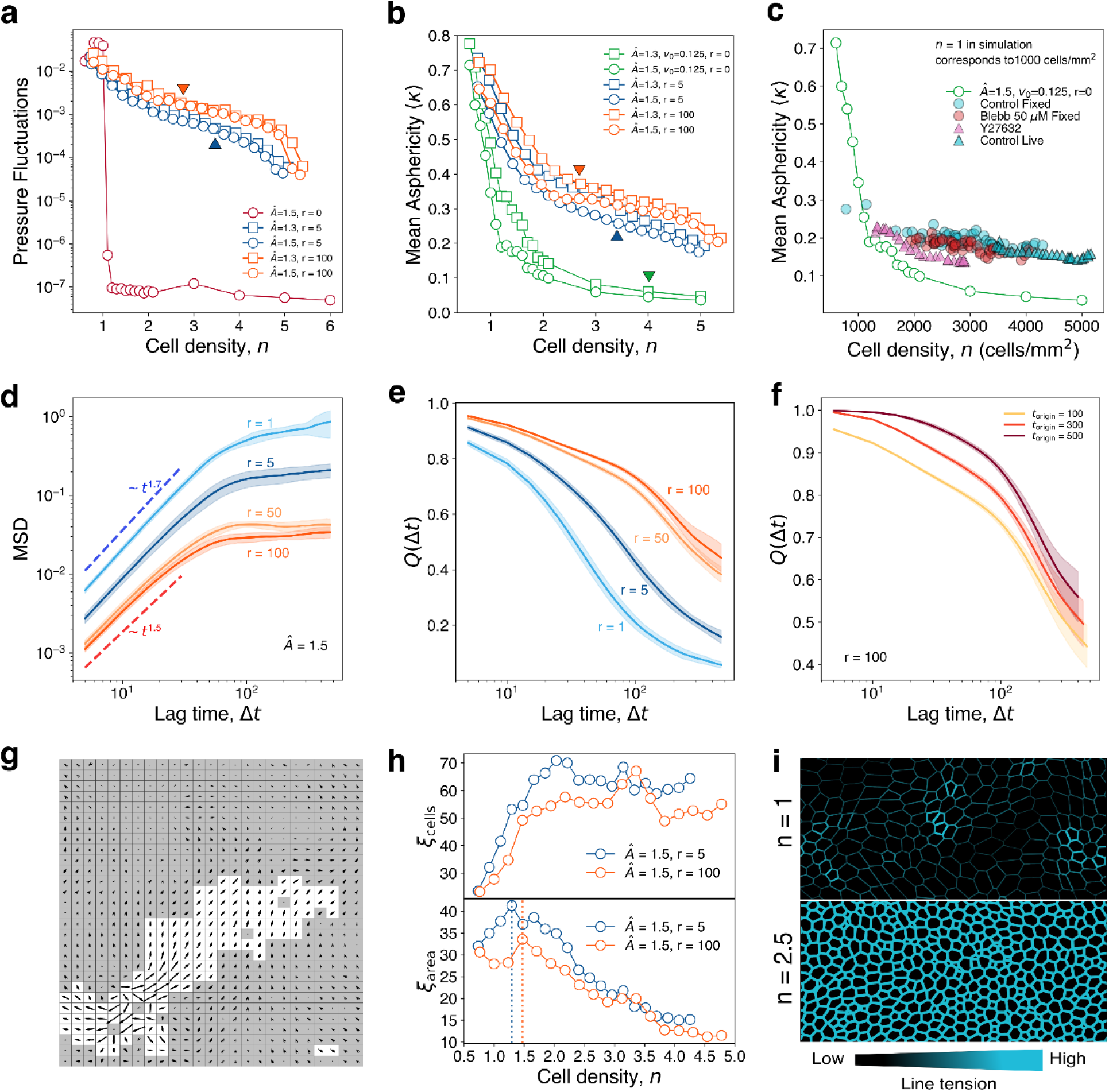
Glassy dynamics in the vertex model with mechanochemical feedback: (a) Pressure fluctuation decay with density depends on the rate of cell division, akin to the cooling rate dependence of passive glass transition. The arrows indicate n_G_(r). The discontinuous constant-density simulation data is also shown for reference. (b) Mean asphericity of the cells, ⟨κ⟩, shows a similar rate dependence. ⟨κ⟩ from constant density simulations (r = 0) decays with n. The arrows indicate n_G_(r). (c) Mean asphericity of untreated, Blebbistatin-treated, and Y27632-treated cells shows trends similar to simulations performed at different division rates, which is consistent with the known effect of Blebbistatin and Y27632 on epithelial cells (see text). (d) Mean squared displacement (MSD) as a function of lag time (Δt) for different division rates shows faster transition to subdiffusive plateaus at higher division rates. (e) The overlap function Q(Δt) exhibits similar behaviour, suggesting that the tissue transitions into the glassy state more rapidly at a higher division rate (see legends in d). The shaded regions indicate the standard deviation. (f) Q(Δt) calculated with respect to different time origins (for trajectories of the cells in simulation) shows ageing. (g) The simulations show persistent dynamic heterogeneities. (h) Consistent with prior experiments, ξ_cells_, the size of dynamic heterogeneity increases with cell density at low densities, peaks at an intermediate density and saturate to a slightly lower value at high densities. The normalized area of DH has been computed as ξ_area_ = ξ_cells_/n. The markers show mean ξ_h_ and ξ_area_. (i) A colormap of the line tension, 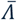 at low and high densities is consistent with E-cadherin expression in the experiments. The colormaps also show contractile instabilities. For all the panels (except h and i), we simulated a system with 10×10 box size, corresponding to 1000 cells at n = 1. Panel h is generated from a simulation of a system with 20×20 box size. For all panels, error bars or shaded regions indicate standard deviation.

Because tissue rigidity transition is driven by changes in cellular shapes, we tested whether the glass transition is also manifested in the parameters describing the cell shapes. To do so, we computed the anisotropy of the cell shape through the mean asphericity (SI-theory), ⟨*κ*⟩, as a function of *n*. In the quasistatic limit (*r* = 0), ⟨*κ*⟩ decreases monotonically (Fig. 5b), with marked difference above and below *n* ≈ 4. For simulations with nonzero division rates and motility, the ⟨*κ*⟩ trajectory deviates significantly from the quasistatic limit (Fig. 5b, Supplementary Fig. 10). The rate-dependence of the glass transition is observed here as well; systems with lower division rates show a lower ⟨*κ*⟩ in the glassy phase and a higher *n*_*G*_.

To experimentally verify these results, we perturbed the tissue with drugs which inhibit cellular contractility: Y27632 and Blebbistatin, both of which lowered 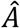 by preventing myosin binding and reduced growth rate of cells and higher shape indices (Supplementary Fig. 12). The effect of the reduced growth rate can be captured by a lower effective growth rate *r*_*eff*_ in our model. By segmenting the cells, we computed ⟨*κ*⟩ at different densities *n* under various treatment conditions (Fig. 5c). Consistent with our theoretical observations, we find that the lower *r*_*eff*_ at higher drug concentrations led to a delayed glass transition, as indicated by lower ⟨*κ*⟩ values (Fig. 5c).

To prove beyond doubt that the observed dynamics arises from an underlying glass transition, we next computed the MSD (Fig. 5d) and the overlap function (*Q*(Δ*t*), Fig. 5e), both of which showed flattening at intermediate timescales at all *r*. In particular, the relaxation of *Q*(Δ*t*) shows the classic signatures of ageing (Fig. 5f). Additionally, the flattening was more pronounced at higher *r* (Fig. 5e), which is consistent with our claim that the glass transition occurs sooner for higher division rates. Although this observation contradicts previous theories describing glassy dynamics in biological cells, where a high division rate enhances fluidisation^26,27,53,54^, it agrees with experiments describing glassy dynamics in epithelial cells^9,11,12,15,16,52,55–60^. Next, we calculated the cell velocities from the simulations, which exhibited features of dynamic heterogeneity (Fig. 5g). The size of the dynamic heterogeneity, *ξ*_*cells*_ or *ξ*_*area*_, changed nonmonotonically with density (Fig. 5h). These observations are consistent with prior experiments on glassy tissue dynamics^7,8,10^, thereby establishing our claim.

#### 2.2.4 Mechanochemical feedback engenders dynamic heterogeneity and glassy dynamics

Because cell motility is inversely related to cell density in our model, the dynamic heterogeneity reflects the underlying spatiotemporal heterogeneity in the tissue. Indeed, experiments^12^ including ours (Fig. 2l) suggest that mechanochemical feedback links heterogeneity in local cell density to heterogeneity in cell motility. Hence, the origin of dynamic heterogeneity can be traced to static density heterogeneity arising from elastic instabilities in the underlying tissues. Indeed, through the analysis of the crystalline ground state, we find that, beyond a threshold density, the P-wave modulus, *M* = *B* + *G*, becomes negative, while *G* stays positive (Fig. 4c, SI-theory). This transition marks the onset of contractile instabilities, during which the homogeneous tissue spontaneously separates into high- and low-density phases. Because *G* > 0 and the tissue is confluent, the phase-separated tissue is stable, and the high- and low-density phases mature into high- and low-density regions (Supplementary Fig. 11) that we identify with the actin hot- and coldspots. Indeed, as in the experiments, the mechanics of the high- and low-density regions in our simulation arise from the line tension and contractility, respectively (Fig. 5i, Supplementary Fig. 10). As the high-density regions slowly grow and percolate through the system, cell motility remains high only in the spatially heterogeneous low-density regions, giving rise to the dynamic heterogeneity and slow glassy dynamics. Also, because of this reason, stresses generated by cell division mobilize only the cells in the low-density regions, ensuring tissue solidification through cell division.

2.3 Mechanochemical feedback stabilises activity clusters to generate long-term collective oscillations

The glass transition, driven by crowding and mechanochemical feedback, enables cells to assemble into jammed and unjammed clusters, characterized by high and low actin levels, respectively. Next, we wondered about the stability of these clusters over time and whether collective temporal patterns in biochemical signaling can emerge from dynamic heterogeneity. When we measured spatial trends in actin expression by imaging LifeAct MDCK cells over time (Fig. 6a and b), we observed collective hour-scale actin oscillations, with locally distinct time periods in hotspots and cold spots-hotspots exhibited dominant 11-hour oscillations in actin expression, while coldspots showed 4-hour oscillations (Fig. 6c, d, f), suggesting cold spots are more dynamic than hotspots. The difference in actin expression in hotspots and coldspots is large enough to not transition a hotspot into a cold spot, and vice versa, even during the oscillatory cycle (Supplementary Fig. 8a, 8b, 8e), suggesting that these regions are not interchangeable. Similar oscillatory behavior was also observed in traction force analysis, which showed global oscillations over a timescale of several hours (Supplementary Fig. 15). These results indicate that dynamic heterogeneity clusters are also synchronized in their biochemistry, with correlated and stabilized actin expression that oscillates over a timescale of several hours. To explain the biochemical origin of the observed oscillations, we consider ERK, a common biochemical oscillator.

**Figure 6:**
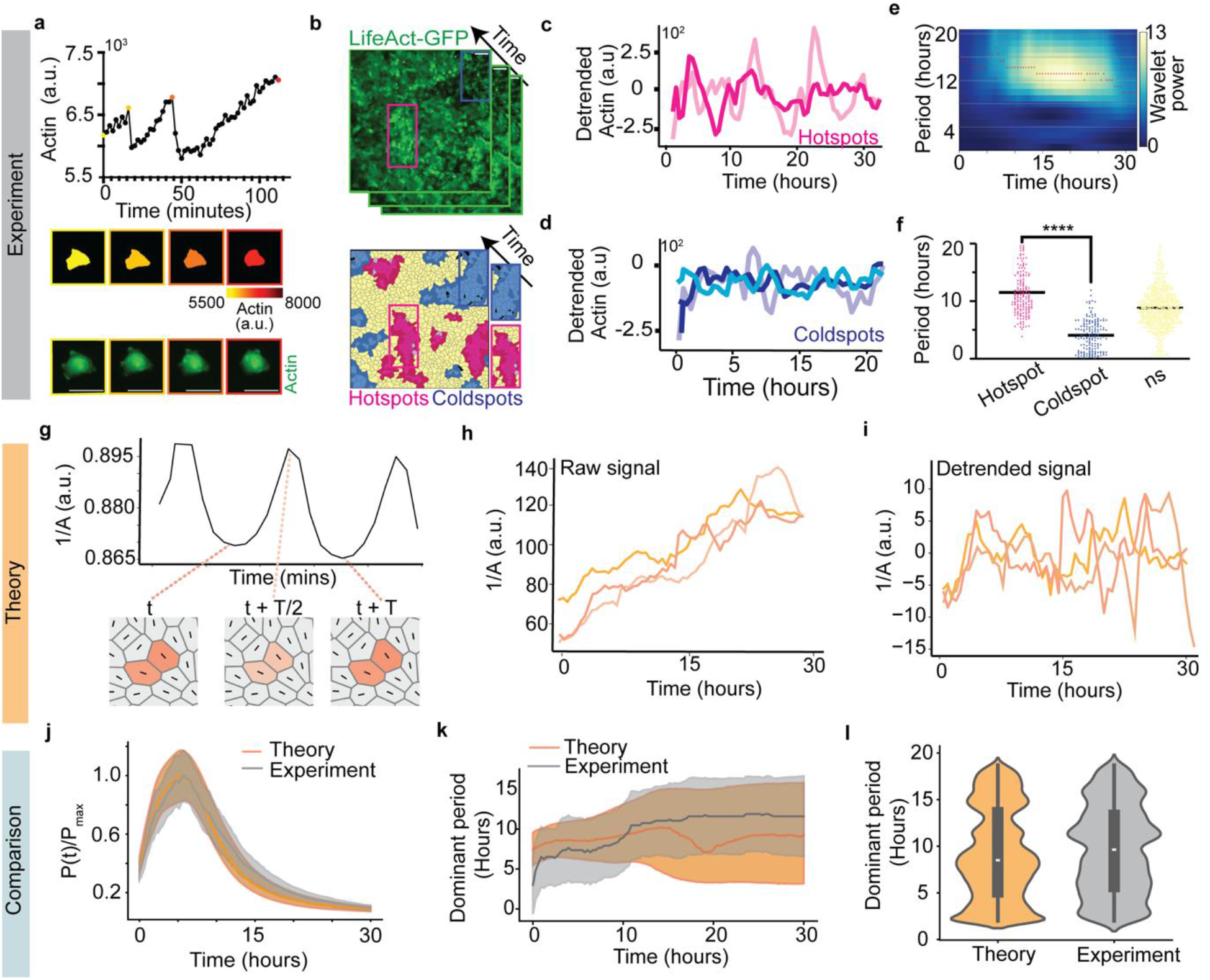
Spatially collective temporal dynamics from the crosstalk of MCFL-I and MCFL-II: a) Single MDCK cells show periodic actin oscillations of ~ 20 min period. Top-Plot showing Actin intensity over time in single MDCK cells without neighbours. Raw images of single cells (bottom), and cell images pseudocolored for Actin (middle). The color of the box around the images (middle and bottom) correspond to the timepoint of the image corresponding to the coloured dots in the top image. b) In contrast, confluent epithelia (top) exhibit collective temporal actin oscillations that depend on spatial actin distribution (bottom), c) Actin hotspots show oscillations with ~10-hour period, d) and Actin coldspots show faster oscillations with ~4-hour period. e) Wavelet analysis of the detrended experimental signals shows decreasing wavelet power and increasing period over time, f) Comparison of dominant period at the hotspots and coldspots, g) Our model predicts single cell area oscillation of ~ 18 min period, originating from the MCFL-II. h) Because the actin level in the cell is proportional to 1/A(1⁄A), the latter is plotted here as a function of time, i) Detrended 1/A is plotted as a function of time, j) Distributions of oscillation power over time, and k) Distributions of oscillation period over time show quantitative match between theory and experiments, l) Comparison of the dominant oscillation period from theory and experiment. Scale bars=50μm. In figures j-l, lines represent the median and the shaded regions represent 25^th^ to 75^th^ quartile.

Interestingly, oscillations in ERK and actin signal are reported in mammalian cells before, including in immune cells^28^, and in embryonic stem cells^29,30^. However, in those studies, oscillations are over timescales of several minutes. Interestingly, *lifeAct* MDCK cells, when cultured as single cells, i.e. in the absence of cell-cell interactions, show oscillatory behaviour of actin with a period of approximately 20 mins (Fig. 6a). The transition from minutes timescale of actin oscillations in single cells, to hours timescale of oscillations in crowded monolayer reflects the coupling of glassy dynamics to cellular biochemistry, allowing for stabilized biochemical signalling in the local dynamic heterogeneity clusters. Furthermore, wavelet power initially increased, and then decreased with cell density, suggesting oscillations strengthen when cells crowd, but dampens with overcrowding (Fig 6e and Supplementary Fig. 7j). Finally, ERK inhibition using PD98059 led to loss in actin oscillations, as observed in temporal actin trend (Supplementary Fig. 7d, e), as well as in wavelet and Fourier power analysis (Supplementary Fig. 7f-k) suggesting a mechanochemical feedback loop, coupling ERK and actin with cell density, determines the characteristics of observed biochemical oscillations.

To probe this hypothesis, we explored the role of MCFL-II in regulating tissue dynamics. In the model, the negative feedback loop in MCFL-II couples the nonlinear chemistry of ERK with the mechanics of the actin cytoskeleton (Fig. 4a). It has been shown earlier that MCFL-II, by itself, generates collective oscillations and travelling waves in the absence of cell division^38–40^. In contrast, here, it is coupled to the glassy dynamics induced by cell division and MCFL-I, which should change the nature of the oscillations. Owing to the inverse relationship between actin and cell area in the experiments (SI-theory), we plotted 1/*A* over time to predict cellular actin dynamics. Our model predicts high-frequency cellular actin oscillations, with a period of approximately 18 minutes, for loosely packed monolayers (Fig. 6g), similar to the single-cell oscillations observed in our experiments (Fig. 6a). However, it exhibits a non-oscillating state for densely packed monolayers, punctuated by collective oscillations with a period of several hours at intermediate densities, owing to mechanochemical feedback originating from cell-cell contacts (Fig. 6h-i). Using our model, we can explain the mechanism underlying the collective oscillation as follows. Cell division generates daughter cells with smaller areas that are stabilised by MCFL-I in the crowded regions. As cell area decreases over time, ERK and actin undergo compression-induced oscillation death^40^, causing MCFL-II oscillations to dampen. This manifests as a progressive loss of oscillation amplitude over time (Fig. 6j), increased median oscillation periods (Fig. 6k), and broadly distributed time periods (Fig. 6l), consistent across experiments and the model. Together, we show that in epithelia, biochemical and mechanical signalling modulate each other via mechanochemical feedback loops, resulting in glassy dynamics and unique spatiotemporal oscillations.

## Discussion

Epithelial homeostasis, essential for organ function, relies on regulating cell density and cell shapes. Experiments and theory reveal spatial variability in cell shapes, driven by cell activity, communication, and deformability^52,^. Experimental analysis of crowded epithelial cells also reveal glassy dynamics with typical features including caging, local cooperativity and subdiffusive movement^9,11–15,15,55^. In contrast, existing theoretical studies suggest that cellular activity should fluidise the system and prevent glass transition^7,25–27,53,62^. Due to this mismatch between theoretical predictions and experimental observations, the emergence of glassy dynamics in epithelial cells remains a matter of debate. We identified a missing link in the traditional, crowding-based explanation for glassy dynamics in epithelial tissues, and suggest that cell-to-cell communication may dictate emergent cellular dynamics. Here, we experimentally demonstrate that dynamic heterogeneity is associated with biochemical clustering and suggest that the collective physical properties of the tissue are dictated by mechanochemical feedback, which enables the effects of crowding to be propagated to cellular activity at long length scales and stabilise the locally fluidised, and locally arrested clusters. Thus, we elucidate that activity cooperates with crowding via mechanochemical feedback to slow down the system, contrasting the previous notion of competition between crowding and activity^8,23,24^.

Existing models of tissue solidification focused on the emergence of rigidity only through the lens of mechanics. Yet, as our experiments show, the interplay of cellular signalling and mechanics is essential to explain glass transition. To remedy this, we developed an active vertex model that couples cell mechanics to cell signalling via mechanochemical feedback loops. Specifically, in our model, cell density controls the relative abundance of cortical actin, which increases with increasing cell density, at the expense of basal stress fibres. Because stress fibres are responsible for cell motility, mean motility decreases with increasing density, which ultimately leads to the observed glassy dynamics. A key implication of this observation is that both crowding and feedback are necessary for the glass transition. Indeed, disabling any one of them leads to tissue fluidisation in both experiments and the model.

We also theoretically established that the epithelial glass transition is density-dependent, a key difference from tissue jamming, which is density-independent and is driven entirely by shape^44,46^. Because of the mechanochemical feedback, in our model, cell shapes cannot be changed independently of cell density. As a result, although the tissue solidifies when the shape parameter crosses a threshold, the density *n*_*J*_ at which it occurs varies with the feedback strength. Furthermore, unlike the jamming transition, the tissue solidification occurs discontinuously in the absence of motility when the feedback strength is sufficiently high. This is the fundamental difference between the tissue solidification observed here and the jamming transition, where the rigidity changes continuously across the critical shape parameter. Because a discontinuous rigidity transition is an essential requirement for the glass transition, our observation provides strong support for the existence of a glass transition in epithelial tissues. The active glass transition that we report here is driven by cell division and mechanochemical feedback and contrasts with the motility-driven active glass transition in tissues^45^. Because of the feedback, motility cannot be controlled independently of the cell shape, and a tissue is constrained to follow a particular trajectory in the shape-motility space, which changes with the feedback strength. Hence, we expect to observe distinct tissue dynamics as the feedback strength is varied in the model. Investigation in this direction may explain how cancerous tissues behave differently across tissues^63^, and may provide a fundamental understanding of the emergent properties of active viscoelastic materials.

We show that mechanochemical feedback loops result not only in glassy spatiotemporal relaxation but also in unique collective biophysical oscillations as observed in cellular actin levels, area, and traction forces. We observed two interesting trends: first, time period of emergent actin oscillations is several hours, as opposed to typical minutes time scale oscillations observed previously in mammalian cell, and second, time period of oscillations are distinct for locally jammed and unjammed clusters, with the less dynamic jammed clusters oscillating over 10 hours period, and the more dynamic unjammed clusters oscillating over 4 hours period, similar to the period of cell area oscillations in MDCK monolayers reported earlier. Thus, we elucidate the mechanism of Actin oscillations, which have implications in morphogenesis and other functions^65–73^. Consistent with our findings linking actin oscillations to cell shape dynamics, coordinated actomyosin and cell shape oscillations have also been reported during several morphogenetic processes^65,66,71^. Clustering of cells and temporal oscillations are both perturbed if mechanochemical feedback is inhibited in the jammed state.

Together, we establish that an interplay between active mechanochemical feedback and cell crowding enables the emergence and stabilization of glassy dynamics in active epithelial tissues. The glass transition in biological cells has implications ranging from the development of organs to diseases^15–22,60^, yet we are only now beginning to understand the biophysical crosstalk that leads to its emergence. We believe our study represnts a first step towards enhancing understanding of the spatiotemporal dynamics of epithelial cells, specifically revealing the emergence and maintenance of dynamic heterogeneity clusters through active cell-to-cell communication.

## Methods

### Cell culture

Madin-Darby canine kidney cells MDCK cells, LifeAct-GFP MDCK cells were cultured in Dulbecco’s Modified Eagle Medium (DMEM, Gibco) supplemented with 10 μg ml^−1^ streptomycin (Pen Strep, Invitrogen), and 5% fetal bovine serum (FBS, Invitrogen) in a humidifier incubator maintained at 37°C and 5% CO_2._.

### Widefield Microscopy

Fluorescence imaging was carried out on a Zeiss Axio Observer 7 inverted microscope equipped with a scientific sCMOS camera (Iberoptics). Images were captured using both a 20X objective (air) or 40X objective (air) for live imaging and a 63X oil-immersion objective for immunofluorescence images that were used for LISA cluster analysis. For live experiments, the samples were placed in a stage-top humidified incubator and maintained at 37 °C with 5% CO_2_ during the imaging.

### Traction force microscopy

First, glass-bottom dishes (Cellvis) were activated by treating first with 0.1 % NaOH, followed by 4% APTMS treatment ((3-Aminopropyl)trimethoxysilane, Sigma #281778) for 15 min and 2% glutaraldehyde (Sigma #354400) treatments. Polyacrylamide gels of different stiffness were prepared by mixing different ratios of Acrylamide and Bisacrylamide with TEMED and APS to initiate polymerisation, mixed with fluorescent beads as reported previously (r). A sandwich of the gel mix between the glass bottom dish and a Sigmacote (Sigma)-treated coverslip for hydrophobicity was allowed to polymerise, and the coverslip was then removed. Then, the gel surface was functionalised by coating with Sulfo-SANPAH (Invitrogen) and UV treatment, followed by coating with Fibronectin (Sigma) to allow cell adhesion to the gel. Cells were then seeded onto the gels and grown to confluence. Then images of the cells and beads were acquired, followed by drug treatment on the microscope stage followed by image acquisition of the beads and cells post addition of the drug. Then, the monolayer was detached from the gel by the addition of trypsin to obtain reference images. Then, images were aligned to correct stage drift using reference positions that were taken in cell free regions pre and post trypsin addition, followed by obtaining bead displacements. From the bead displacements, the traction field was computed the FTTC ImageJ plugin^74^. The traction field was then plotted using a custom written code on MATLAB.

For traction force microscopy experiments with Blebbistatin, pre-blebbistatin stressed bead image and Actin image was taken first. Then 20μM Blebbistatin was added on the microscope stage and 30 minutes later, post-blebbistain stressed bead and Actin images were captured. This was followed by trypsinisation and acquisition of the reference bead images.

### Immunostaining

Cell fixation was done with 4% formaldehyde diluted in 1× phosphate-buffered saline (PBS; pH 7.4) at room temperature (RT) for 10 min, followed by 1× PBS washes (three times). Cell permeabilization was carried out with 0.25% (v/v) Triton X-100 (Sigma) in PBS for 10 min at RT, followed by washing three times with PBS to remove the reagent. To block non-specific antibody binding, samples were incubated in 2% BSA and 5% FBS in PBS at RT for 45 min. The blocking buffer was removed after 45 min, and the primary antibody dilution (1:300) prepared in the blocking buffer was added to the samples. The samples were incubated with the primary antibody, at 4°C overnight. Then, samples were washed three times with 1× PBS. Each wash was done for 5 minutes on a gel rocker. Next, secondary antibodies tagged with a fluorophore were (1:300) prepared in 25% blocking buffer diluted in PBS and added to the sample for 60 min at RT. To counterstain cell nuclei, the samples were added with a DNA-binding dye, 4′,6-diamidino-2-phenylindole (DAPI; 1 μg/mL in PBS, Invitrogen), along with the secondary antibody solution. Then, thorough washing of the samples was done with PBS before imaging.

The following primary antibodies were used: Zo1- (1:200) (CST #5406), E-Cadherin (24E10) -(1:500) (CST #3195), P-myosin (Ser19) (1:50) (CST #3671), Beta catenin (1:500) (CST #9562). The following secondary antibodies were used: Alexa Fluor 488-conjugated anti-rabbit IgG (1:500) (CST #4412), Alexa Fluor 488-conjugated anti-mouse IgG (1:500) (Invitrogen #A32723), Alexa Fluor 594-conjugated anti-rabbit IgG (1:500) (CST #8889). F-Actin was statined with Alexa Fluor 594-conjugated phalloidin (1:1000) (CST #12877S) and 4′,6-diamidino-2-phenylindole (DAPI) (1:1000 and 1:2000) (CST #4083S) was used to stain the nucleus.

### Drug treatments

#### Blebbistatin

For immunocytochemistry experiments, Blebbistatin (Sigma) was added at a final concentration of 50μM and 100μM after the monolayer reached high density, and dishes were fixed 6 hours after the addition of the drug. For live imaging experiments, the final concentration was 10μM for the duration of imaging.

#### Thymidine double block

Cells were treated with 2mM Thymidine (Sigma) and incubated for 16 hours. After this, they were washed and cultured in regular cell culture media for 9 hours. Then, 2mM Thymidine was added again and incubated for 14 hours. Cells were then washed and used immediately for experiments.

#### PD98059

Cells were treated with 10 μM PD98059 (CST#9900) for the duration of the live imaging. Stock solution of 20mM was prepared in DMSO, as per the instructions of the supplier. Finally, the stock solution was diluted in cell culture medium to obtain the final concentration.

#### Y27632

Cells were treated with 10 μM Y27632 (Sigma #Y0503) for the duration of the live imaging. Stock solution of 10mM was prepared in DMSO, as per the instructions of the supplier. Finally, the stock solution was diluted in cell culture medium to obtain the final concentration.

#### Cell stretch

To perform cell stretching, a custom-made device was used, which applied mechanical strain to a PDMS substrate. PDMS was prepared by mixing the elastomer and crosslinker (SYLGARD 184, corning) at a ratio of 10:1 by weight, followed by thorough mixing and degassing the solution, which was then cast in a plexiglass mould and cured at 80 °C for 6 hours. Then, the chamber was removed from the cast and thoroughly washed with 70% Ethanol. Plasma treatment in an oxygen environment was then performed for 10 min to make the PDMS surface hydrophilic, followed by a fibronectin coating to allow cell adhesion. LifeAct-MDCK Cells were seeded on the PDMS chamber at different densities to obtain confluent and sub-confluent monolayers. Cells were then imaged in the relaxed configuration of the PDMS using 10X objective, Zeiss Axio Observer. The PDMS chamber with cells was then kept in the stretched configuration for 5 minutes using the custom cell stretcher device and immediately kept back on the microscope to obtain post-stretch images of the cells.

### Image analysis

#### Cell segmentation

Images were segmented with cellpose^75^ to get the cell masks with Actin as the primary and DAPI as the second channel in immuostained images. For segmentation in timelapse microscopy images, phase-contrast images were used and were segmented with a custom trained model on cellpose. Area, Perimeter, ellipse fit, and mean fluorescence intensity within masks were computed using LabelsToRois plugin^76^ in FIJI for fixed images and using Trackmate plugin for live microscopy data. To remove wrongly segmented spots misidentified as cells by the software, cells were filtered out based on area cutoff in subsequent analysis.

#### Cell tracking

Cells were tracked using Trackmate Plugin^78-79^ from the segmented images of Cellpose using the Intersection of Union (IoU) method. Prior to tracking,wrongly segmented labels were removed using the area filter on trackmate.

#### Moran’s Index^**77**^

The Global Moran’s Index was calculated using the following definition:

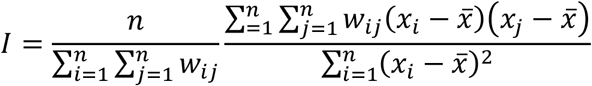

where *w*_*ij*_ represents the weights assigned to the neighbours of each cell using a custom written R code. We have assumed equal weights to all the immediate neighbours defined as the cells which are in direct contact with the cell of interest. Moran’s plot was plotted with the mean intensity of cell of interest on x axis, and the average intensities of its neighbours on y axis. Correlograms were plotted by computing the global Moran’s Index at different lag neighbours. The Local Moran’s Index was calculated to determine the spatial autocorrelation of the proteins of interest for every cell and its neighbours which are in direct contact with it using the following definition:

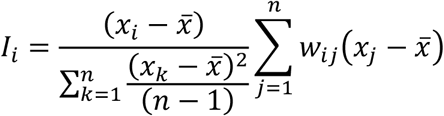

where *w*_*ij*_ represents the weights assigned to the neighbours of each cell. We assigned equal weights to all the first nearest neighours of a cell. The “localmoran” function in the ‘spdep’ package in R was used for this purpose. For each cell and its neighbours, based on deviation from the global mean of the protein amount and the value of its local Moran’s index, the cells were classified into four categories “High-High”, “High-Low”, “Low-High” and “Low-Low”, while insignificant deviations were marked as such and were plotted in the form of a LISA^31^ cluster map. To assess statistical significance, we used 999 Monte carlo simulations and used a p-value cutoff of 0.05. To capture the spatial clusters to its complete extent of the LISA Actin clusters, cells that are immediate neighbours to the High-High and Low-Low clusters have been represented with darker shades, since LISA analysis only captures the core of spatial clusters^80^.

#### Dynamic heterogeneity domains

To identify the fast and slow regions of dynamic heterogeneity, we used LISA clustering. ‘Low-low’ clusters were determined to be the slow regions of dynamic heterogeneity, and ‘High-High’ clusters were determined to be the fast regions of dynamic heterogeneity.

#### Mean square displacement

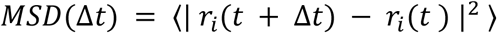

#### Overlap function

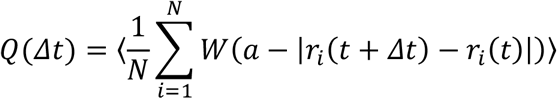

As there are cell divisions in both experiment and simulation, we only consider those cells that are present at both *t* and *t* + Δ*t* when we calculate the MSD and Q(t). Here, angular brackets denote average over *t* and ensembles (in simulation). For Fig. 2f, *a* = 3.25 *μm*. For Fig. 3f, *a* = 10 *μm*. For simulation, *a* = 0.1.

Both in MSD and overlap function, the COM drift has been subtracted during calculation.

#### Actin signals

Actin timeseries signals were obtained from the fluorescence intensity of LifeAct-GFP images by coarse-graining the image with grid sizes comparable to cell size, i.e 30*μ*m, using custom-written python code, from confluent monolayers. For Actin signals from single LifeAct-MDCK cells without any neighbours, per cell intensity was obtained by cell segmentation followed by measuring mean intensity within the cell outline determined by Trackmate.

#### Particle Image Velocimetry

To compute PIV, iterative PIV using FTTC registration method was computed using the MATLAB toolbox PIVlab^81^. Grid sizes were chosen based on cell size.

#### Relative cell pressure analysis

Relative cell pressures were computed using Bayesian Force inference^82^ adapted using custom code for 2D monolayer data, based on the principle of balancing forces at cell vertices.

#### Stress fiber analysis

Length of stress fibers were manually measured in FIJI using the line and measure tools.

### Timeseries analysis

#### Pyboat

PyBoat^83^ is python-based fully automatic stand-alone software that integrates multiple steps of non-stationary oscillatory time series analysis which is being used for the quantification of biochemical and physical heterogeneity spatiotemporally. Pyboat implements continuous wavelet analysis using a sliding Morlet wavelet of different frequencies and determines the power for different frequencies at each time. Then, the main oscillatory component is determined from the heatmap of frequency power at different times and frequency. It also provides optimized detrending, amplitude removal, spectral analysis, ridge detection, oscillatory parameters readout and visualization plots along with an integrated batch-processing option. Using the batch processing option, we also get the most dominant time period of all the spatial positions in the time-series. Using pyboat, we sinc-detrended the Actin time-series signal to obtain the detrended signal,with detrending period taken to be the imaging duration to remove the trend of increasing Actin with increasing time due to increasing density.

#### Statistical analysis

Statistical analyses were performed by Unpaired t-test with Welch’s correction using GraphPad Prism 10, unless otherwise mentioned. All experiments were repeated at least three times. Lines represent the median in scatter bar plots unless otherwise mentioned. In violin plot, top line represents 75^th^ percentile, middle line represents the median and the bottom line represents the 25^th^ percentile.

## Data availability

Data supporting the findings of this work are available in this Article and its extended data figures and Supplementary Information. Source data are provided with this paper. All other data supporting the findings of the study are available from the corresponding author upon reasonable request. The time lapse imaging data (other than that provided as video files with the manuscript) can only be made available upon request due to large file sizes and associated storage limitations.

## Code availability

LISA analysis codes are available online at https://github.com/sindhum98/LISA-analysis-for-epithelial-tissue. The codes for modelling can be obtained from the authors upon reasonable request.

## Author contributions

M.V conceived the project. S.M. and M.V. designed experiments. S.M. performed and analyzed all experiments except the traction force microscopy with LifeAct MDCK cells, which was performed by M.B.S. Theoretical model was contributed by S.S. and P.D. T.T. performed the LISA analysis, and T.C. contributed to oscillations analysis using pyBOAT. S.K. contributed to glassy analysis. Analysis and interpretation of data was done by S.M., P.D., Sd.M., S.S., and M.V. M.V. and S.S. developed and wrote the manuscript with help from S.M. and P.D. All authors read, discussed and commented on the manuscript.

## Acknowledgements

We thank Sriram R. Ramaswamy, Saroj K. Nandi, Tamal Das and Srikanth Sastry for critical discussions and suggestions. M.V. is a partner group leader of the Max Planck Society (MPG), Germany, which has supported part of this work. This work is also supported by the Infosys foundation, Anusandhan National Research Foundation-previously called the Science and Engineering Research Board (project number: SERB SRG/2022/000534), and Indo German Science and Technology Centre (IGSTC WISER scheme). SS acknowledges funding from IISc, Axis Bank Center for Mathematics and Computing, and a startup grant from SERB-DST (SRG/2022/000163). We also acknowledge intramural funds at IISc Bangalore for providing support towards equipment and facilities and for salaries/fellowships of the authors. S.M. and Sd.M and acknowledge funding from Prime Minister’s Research Fellowship (PMRF).

## References

1. Debenedetti, P. G. & Stillinger, F. H. Supercooled liquids and the glass transition. Nature 410, 259–267 (2001).

2. Berthier, L. Dynamical Heterogeneities in Glasses, Colloids, and Granular Media. (OUP Oxford, 2011).

3. Berthier, L. & Biroli, G. Theoretical perspective on the glass transition and amorphous materials. Rev. Mod. Phys. 83, 587–645 (2011).

4. Kob, W., Donati, C., Plimpton, S. J., Poole, P. H. & Glotzer, S. C. Dynamical Heterogeneities in a Supercooled Lennard-Jones Liquid. Phys. Rev. Lett. 79, 2827–2830 (1997).

5. Ediger, M. D. Spatially heterogeneous dynamics in supercooled liquids. Annu. Rev. Phys. Chem. 51, 99–128 (2000).

6. Janssen, L. M. C. Active glasses. J. Phys. Condens. Matter 31, 503002 (2019).

7. Berthier, L., Flenner, E. & Szamel, G. How active forces influence nonequilibrium glass transitions. New J. Phys. 19, 125006 (2017).

8. Berthier, L., Flenner, E. & Szamel, G. Glassy dynamics in dense systems of active particles. J. Chem. Phys. 150, 200901 (2019).

9. Angelini, T. E. et al. Glass-like dynamics of collective cell migration. Proc. Natl. Acad. Sci. 108, 4714–4719 (2011).

10. Park, J.-A. et al. Unjamming and cell shape in the asthmatic airway epithelium. Nat. Mater. 14, 1040–1048 (2015).

11. Malinverno, C. et al. Endocytic reawakening of motility in jammed epithelia. Nat. Mater. 16, 587–596 (2017).

12. Garcia, S. et al. Physics of active jamming during collective cellular motion in a monolayer. Proc. Natl. Acad. Sci. 112, 15314–15319 (2015).

13. Smeets, B. et al. Compaction Dynamics during Progenitor Cell Self-Assembly Reveal Granular Mechanics. Matter 2, 1283–1295 (2020).

14. Ilina, O. et al. Cell–cell adhesion and 3D matrix confinement determine jamming transitions in breast cancer invasion. Nat. Cell Biol. 22, 1103–1115 (2020).

15. Mongera, A. et al. A fluid-to-solid jamming transition underlies vertebrate body axis elongation. Nature 561, 401–405 (2018).

16. Schötz, E.-M., Lanio, M., Talbot, J. A. & Manning, M. L. Glassy dynamics in three-dimensional embryonic tissues. J. R. Soc. Interface 10, 20130726 (2013).

17. Atia, L., Fredberg, J. J., Gov, N. S. & Pegoraro, A. F. Are cell jamming and unjamming essential in tissue development? Cells Dev. 168, 203727 (2021).

18. Blauth, E., Kubitschke, H., Gottheil, P., Grosser, S. & Käs, J. A. Jamming in Embryogenesis and Cancer Progression. Front. Phys. 9, (2021).

19. Oswald, L., Grosser, S., Smith, D. M. & Käs, J. A. Jamming transitions in cancer. J. Phys. Appl. Phys. 50, 483001 (2017).

20. Gottheil, P. et al. State of Cell Unjamming Correlates with Distant Metastasis in Cancer Patients. Phys. Rev. X 13, 031003 (2023).

21. Palamidessi, A. et al. Unjamming overcomes kinetic and proliferation arrest in terminally differentiated cells and promotes collective motility of carcinoma. Nat. Mater. 18, 1252–1263 (2019).

22. Vishwakarma, M. et al. Mechanical interactions among followers determine the emergence of leaders in migrating epithelial cell collectives. Nat. Commun. 9, 3469 (2018).

23. Berthier, L. Nonequilibrium Glassy Dynamics of Self-Propelled Hard Disks. Phys. Rev. Lett. 112, 220602 (2014).

24. Klongvessa, N., Ginot, F., Ybert, C., Cottin-Bizonne, C. & Leocmach, M. Active Glass: Ergodicity Breaking Dramatically Affects Response to Self-Propulsion. Phys. Rev. Lett. 123, 248004 (2019).

25. Malmi-Kakkada, A. N., Li, X., Samanta, H. S., Sinha, S. & Thirumalai, D. Cell Growth Rate Dictates the Onset of Glass to Fluidlike Transition and Long Time Superdiffusion in an Evolving Cell Colony. Phys. Rev. X 8, 021025 (2018).

26. Czajkowski, M., M. Sussman, D., Cristina Marchetti, M. & Lisa Manning, M. Glassy dynamics in models of confluent tissue with mitosis and apoptosis. Soft Matter 15, 9133–9149 (2019).

27. Ranft, J. et al. Fluidization of tissues by cell division and apoptosis. Proc. Natl. Acad. Sci. 107, 20863–20868 (2010).

28. Das, J. et al. Digital Signaling and Hysteresis Characterize Ras Activation in Lymphoid Cells. Cell 136, 337–351 (2009).

29. Raina, D., Fabris, F., Morelli, L. G. & Schröter, C. Intermittent ERK oscillations downstream of FGF in mouse embryonic stem cells. Development 149, dev199710 (2022).

30. Nakayama, K., Satoh, T., Igari, A., Kageyama, R. & Nishida, E. FGF induces oscillations of Hes1 expression and Ras/ERK activation. Curr. Biol. 18, R332–R334 (2008).

31. Anselin, L. Local Indicators of Spatial Association—LISA. Geogr. Anal. 27, 93–115 (1995).

32. Berthier, L. Dynamic Heterogeneity in Amorphous Materials. Physics 4, 42 (2011).

33. Serra-Picamal, X. et al. Mechanical waves during tissue expansion. Nat. Phys. 8, 628–634 (2012).

34. Saraswathibhatla, A. & Notbohm, J. Tractions and Stress Fibers Control Cell Shape and Rearrangements in Collective Cell Migration. Phys. Rev. X 10, 011016 (2020).

35. Sknepnek, R., Djafer-Cherif, I., Chuai, M., Weijer, C. & Henkes, S. Generating active T1 transitions through mechanochemical feedback. eLife 12, e79862 (2023).

36. Ioratim-Uba, A. Mechanochemical Active Feedback Generates Convergence Extension in Epithelial Tissue. Phys. Rev. Lett. 131, (2023).

37. Abhishek, M., Dhanuka, A., Banerjee, D. S. & Rao, M. Excitability and travelling waves in renewable active matter. Preprint at 10.48550/arXiv.2503.19687 (2025).

38. Boocock, D. Interplay between Mechanochemical Patterning and Glassy Dynamics in Cellular Monolayers. PRX Life 1, (2023).

39. Boocock, D., Hino, N., Ruzickova, N., Hirashima, T. & Hannezo, E. Theory of mechanochemical patterning and optimal migration in cell monolayers. Nat. Phys. 17, 267– 274 (2021).

40. Dewan, P., Mondal, S. & Sarkar, S. Oscillation death by mechanochemical feedback. Preprint at 10.48550/arXiv.2504.19655 (2025).

41. Farhadifar, R., Röper, J.-C., Aigouy, B., Eaton, S. & Jülicher, F. The Influence of Cell Mechanics, Cell-Cell Interactions, and Proliferation on Epithelial Packing. Curr. Biol. 17, 2095–2104 (2007).

42. Barton, D. L., Henkes, S., Weijer, C. J. & Sknepnek, R. Active Vertex Model for cell-resolution description of epithelial tissue mechanics. PLOS Comput. Biol. 13, e1005569 (2017).

43. Cavagna, A. Supercooled liquids for pedestrians. Phys. Rep. 476, 51–124 (2009).

44. Staple, D. B. et al. Mechanics and remodelling of cell packings in epithelia. Eur. Phys. J. E Soft Matter 33, 117–127 (2010).

45. Bi, D., Yang, X., Marchetti, M. C. & Manning, M. L. Motility-Driven Glass and Jamming Transitions in Biological Tissues. Phys. Rev. X 6, 021011 (2016).

46. Bi, D., Lopez, J. H., Schwarz, J. M. & Manning, M. L. A density-independent rigidity transition in biological tissues. Nat. Phys. 11, 1074–1079 (2015).

47. Bi, D., Zhang, J., Chakraborty, B. & Behringer, R. P. Jamming by shear. Nature 480, 355–358 (2011).

48. Sarkar, S., Bi, D., Zhang, J., Behringer, R. P. & Chakraborty, B. Origin of Rigidity in Dry Granular Solids. Phys. Rev. Lett. 111, 068301 (2013).

49. Sarkar, S. et al. Shear-induced rigidity of frictional particles: Analysis of emergent order in stress space. Phys. Rev. E 93, 042901 (2016).

50. Behringer, R. P. & Chakraborty, B. The physics of jamming for granular materials: a review. Rep. Prog. Phys. 82, 012601 (2018).

51. Chisolm, S. J., Guo, E., Subramaniam, V., Schulze, K. D. & Angelini, T. E. Transitions between cooperative and crowding-dominated collective motion in non-jammed MDCK monolayers. Cells Dev. 181, 203989 (2025).

52. Atia, L. et al. Geometric constraints during epithelial jamming. Nat. Phys. 14, 613–620 (2018).

53. Matoz-Fernandez, D. A., Martens, K., Sknepnek, R., Barrat, J. L. & Henkes, S. Cell division and death inhibit glassy behaviour of confluent tissues. Soft Matter 13, 3205–3212 (2017).

54. Matoz-Fernandez, D. A. Nonlinear Rheology in a Model Biological Tissue. Phys. Rev. Lett. 118, (2017).

55. Park, J.-A. et al. Unjamming and cell shape in the asthmatic airway epithelium. Nat. Mater. 14, 1040–1048 (2015).

56. Nnetu, K. D., Knorr, M., Strehle, D., Zink, M. & Käs, J. A. Directed persistent motion maintains sheet integrity during multi-cellular spreading and migration. Soft Matter 8, 6913– 6921 (2012).

57. Giavazzi, F., Malinverno, C., Scita, G. & Cerbino, R. Tracking-Free Determination of Single-Cell Displacements and Division Rates in Confluent Monolayers. Front. Phys. 6, (2018).

58. Lin, S.-Z. et al. Universal Statistical Laws for the Velocities of Collective Migrating Cells. Adv. Biosyst. 4, 2000065 (2020).

59. Kim, J. H. et al. Unjamming and collective migration in MCF10A breast cancer cell lines. Biochem. Biophys. Res. Commun. 521, 706–715 (2020).

60. Vishwakarma, M., Thurakkal, B., Spatz, J. P. & Das, T. Dynamic heterogeneity influences the leader–follower dynamics during epithelial wound closure. Philos. Trans. R. Soc. B Biol. Sci. 375, 20190391 (2020).

61. Pandey, S. et al. The structure-dynamics feedback mechanism governs the glassy dynamics in epithelial monolayers. Soft Matter 21, 269–276 (2025).

62. Tjhung, E. & Berthier, L. Discontinuous fluidization transition in time-correlated assemblies of actively deforming particles. Phys. Rev. E 96, 050601 (2017).

63. Datta, A. et al. Differential interfacial tension between oncogenic and wild-type populations forms the mechanical basis of tissue-specific oncogenesis in epithelia. eLife 14, (2025).

64. Zehnder, S. M., Suaris, M., Bellaire, M. M. & Angelini, T. E. Cell Volume Fluctuations in MDCK Monolayers. Biophys. J. 108, 247–250 (2015).

65. Blanchard, G. B., Scarpa, E., Muresan, L. & Sanson, B. Mechanical stress combines with planar polarised patterning during metaphase to orient embryonic epithelial cell divisions. Development 151, dev202862 (2024).

66. He, L., Wang, X., Tang, H. L. & Montell, D. J. Tissue elongation requires oscillating contractions of a basal actomyosin network. Nat. Cell Biol. 12, 1133–1142 (2010).

67. David, D. J. V., Tishkina, A. & Harris, T. J. C. The PAR complex regulates pulsed actomyosin contractions during amnioserosa apical constriction in Drosophila. Development 137, 1645–1655 (2010).

68. Kim, H. Y. & Davidson, L. A. Punctuated actin contractions during convergent extension and their permissive regulation by the non-canonical Wnt-signaling pathway. J. Cell Sci. 124, 635–646 (2011).

69. Martin, A. C., Kaschube, M. & Wieschaus, E. F. Pulsed contractions of an actin–myosin network drive apical constriction. Nature 457, 495–499 (2009).

70. Rauzi, M.Lenne, P.-F. & Lecuit, T. Planar polarized actomyosin contractile flows control epithelial junction remodelling. Nature 468, 1110–1114 (2010).

71. Solon, J., Kaya-Çopur, A., Colombelli, J. & Brunner, D. Pulsed Forces Timed by a Ratchet-like Mechanism Drive Directed Tissue Movement during Dorsal Closure. Cell 137, 1331– 1342 (2009).

72. Wollman, R. & Meyer, T. Coordinated oscillations in cortical actin and Ca2+ correlate with cycles of vesicle secretion. Nat. Cell Biol. 14, 1261–1269 (2012).

73. Vicker, M. G. F-actin assembly in Dictyostelium cell locomotion and shape oscillations propagates as a self-organized reaction–diffusion wave. FEBS Lett. 510, 5–9 (2002).

74. Tseng, Q. et al. Spatial organization of the extracellular matrix regulates cell–cell junction positioning. Proc. Natl. Acad. Sci. 109, 1506–1511 (2012).

75. Stringer, C., Wang, T., Michaelos, M. & Pachitariu, M. Cellpose: a generalist algorithm for cellular segmentation. Nat. Methods 18, 100–106 (2021).

76. Waisman, A., Norris, A. M., Elías Costa, M. & Kopinke, D. Automatic and unbiased segmentation and quantification of myofibers in skeletal muscle. Sci. Rep. 11, 11793 (2021).

77. Moran, P. A. P. The Interpretation of Statistical Maps. J. R. Stat. Soc. Ser. B Methodol. 10, 243– 251 (1948).

78. Ershov, D. et al. TrackMate 7: integrating state-of-the-art segmentation algorithms into tracking pipelines. Nat. Methods 19, 829–832 (2022).

79. Tinevez, J.-Y. et al. TrackMate: An open and extensible platform for single-particle tracking. Methods 115, 80–90 (2017).

80. GeoDa : An Introduction to Spatial Data Analysis - Anselin - 2006 - Geographical Analysis - Wiley Online Library. https://onlinelibrary.wiley.com/doi/full/10.1111/j.0016-7363.2005.00671.x.

81. Thielicke, W. & Sonntag, R. Particle Image Velocimetry for MATLAB: Accuracy and enhanced algorithms in PIVlab. J. Open Res. Softw. 9, (2021).

82. Ishihara, S. & Sugimura, K. Bayesian inference of force dynamics during morphogenesis. J. Theor. Biol. 313, 201–211 (2012).

83. Mönke, G., Sorgenfrei, F. A., Schmal, C. & Granada, A. E. Optimal Time Frequency Analysis for Biological Data - pyBOAT. http://biorxiv.org/lookup/doi/10.1101/2020.04.29.067744 (2020) xdoi:10.1101/2020.04.29.067744.

