## Supplementary Information file for "Glassy dynamics in active epithelia emerge from an interplay of mechanochemical feedback and crowding"

**Table of contents:**


Supplementary Figures 1 to 14 3

Supplementary Movies S1 to S10 18

Supplementary Theory 19

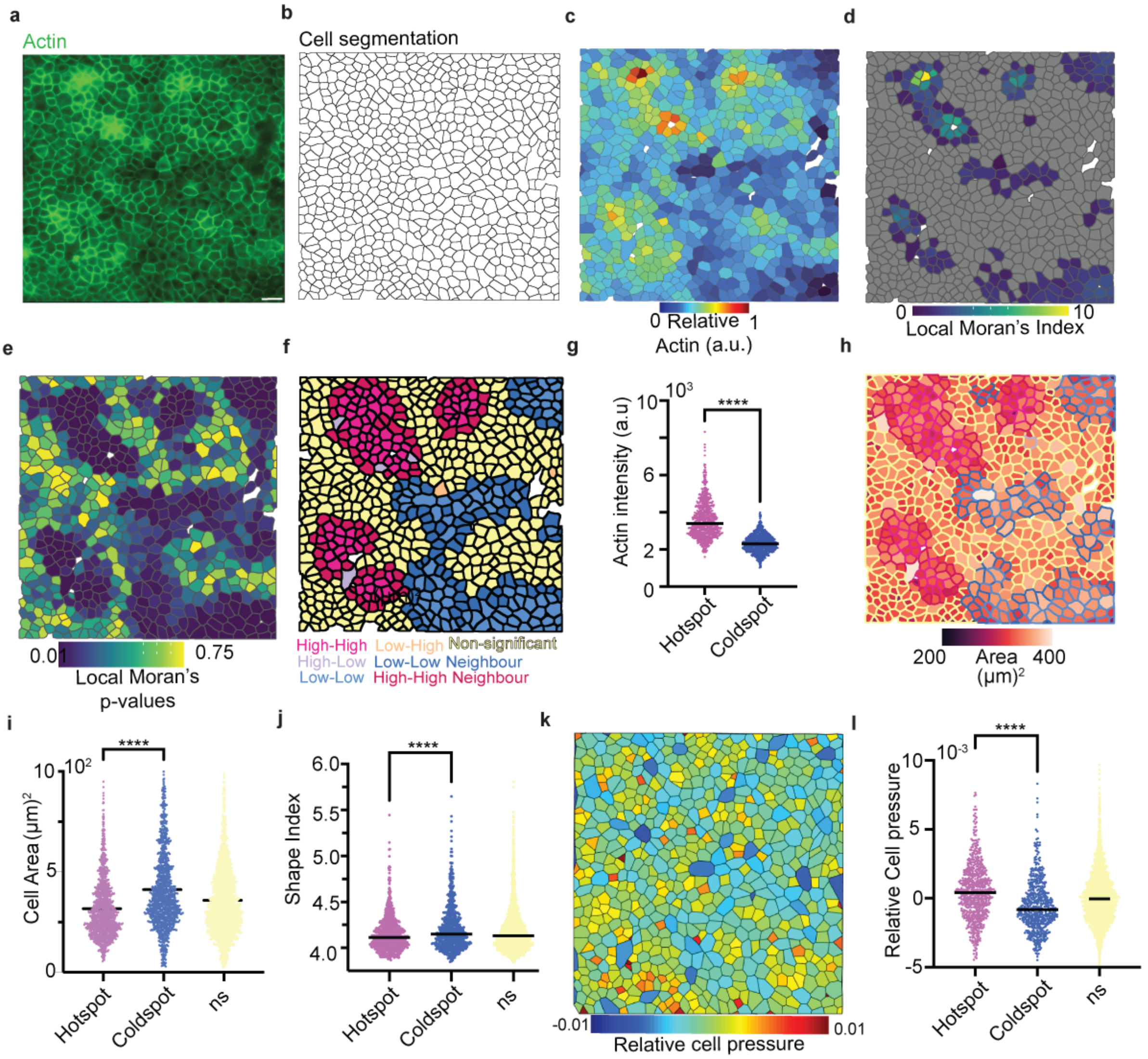


***Supplementary Fig 1- Spatial characterization of Actin distribution and Mechanochemical feedback:*** *a) Raw microscopy image of cells stained for Actin with Phalloidin, b) Cell boundaries obtained from cell segmentaion, c) Cells pseudocolored for normalized F-Actin levels, d) Heatmap of Local Moran's Index of cells, e) Heatmap of p-values of cells of Local Moran's Index obtained from , f) LISA cluster map of F-Actin, g) Actin intensity of cells classified as High-High (Hotspotspink) and Low-Low (Coldspots-blue), h) Cells color-coded for cell area and outlines colored for LISA cluster category, i) Scatter dot plot of single cell area in High-High, Low-Low and Nonsignificant clusters, j) Scatter dot plot of Shape Index in High-High, Low-Low and Non-significant clusters, k) Heatmap of relative cell pressures obtained by Bayesian Force Inference, l) Scatter dot plot of relative cell pressures in High-High, Low-Low and Non-significant LISA clusters. Scale bars= 50μm. Lines represent the median.*

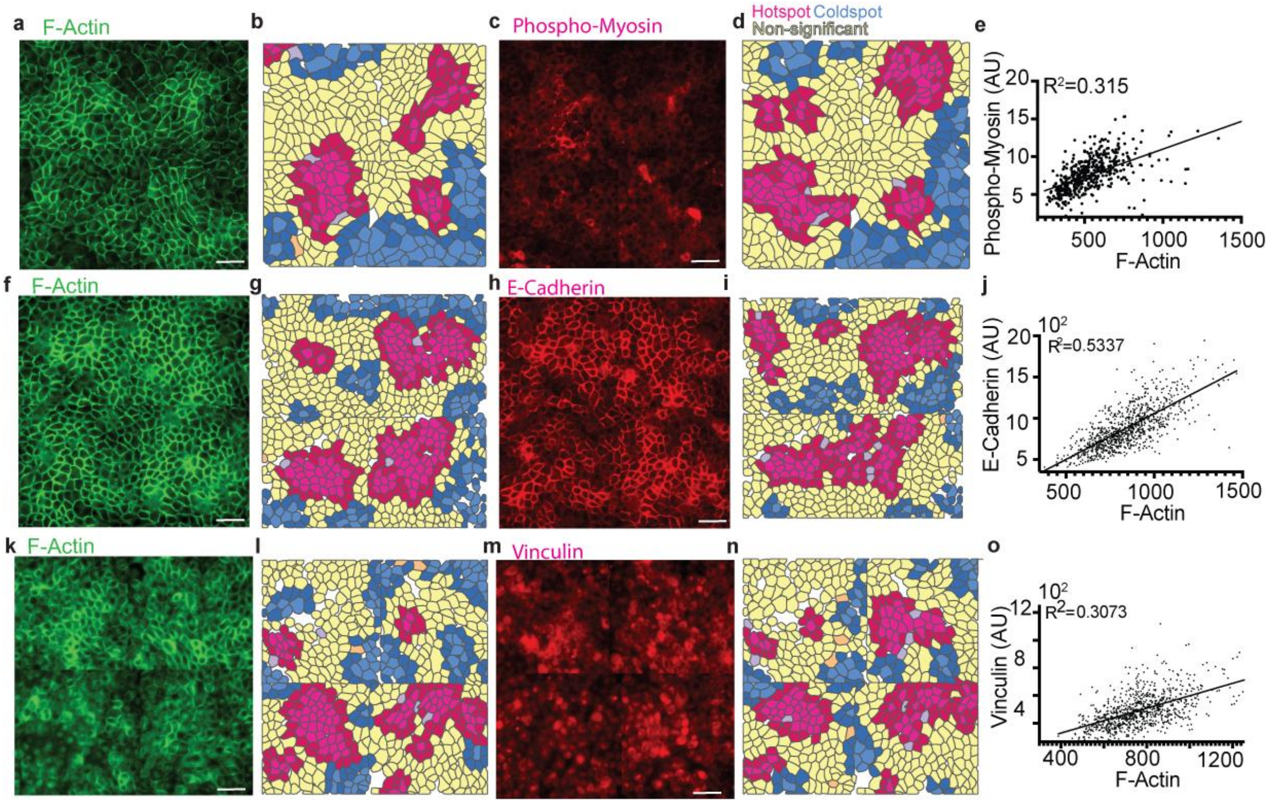


***Supplementary Figure 2: Spatial correlation in cytoskeletal proteins correspond to the spatial clustering in Actin:*** *a) Immunostaining image of F-Actin, b) LISA cluster map of F-Actin, c) Immunostaining image of P-Myosin corresponding to a), d) LISA* cluster map of P-Myosin, e) Scatterplot of F-Actin vs P-Myosin, *f) Immunostaining image of F-Actin corresponding to h, g) LISA cluster map of F-Actin corresponding to f, h) Immunostaining image of E-Cadherin corresponding to f), i) LISA* cluster map of E-Cadherin, j) Scatterplot of F-Actin vs E-Cadherin, k*) Immunostaining image of F-Actin corresponding to m, l) LISA cluster map of F-Actin corresponding to k, m) Immunostaining image of E-Cadherin corresponding to k), n) LISA* cluster map of Vinculin, o) Scatterplot of F-Actin vs Vinculin.

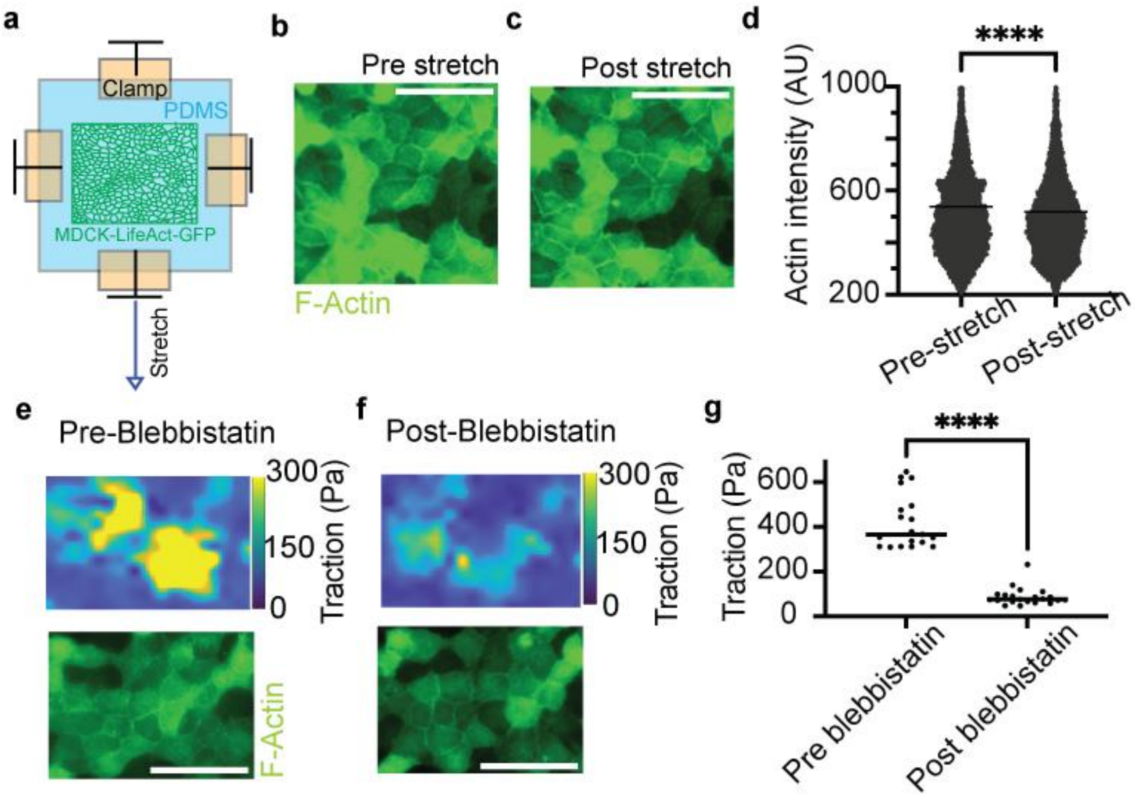


***Supplementary Figure 3: Dynamic Mechanochemical feedback**: a) Schematic sketch of the cell stretching experiment, b) Representative LifeAct MDCK cells before(b) and after (c) stretching the substrate, d) Comparison of Actin intensity in cells pre and post stretch, e) Traction force (top) and Actin from a representative region before (e) and after (f) Blebbistatin addition, g) Comparison plot of traction pre and post Blebbistatin addition.*

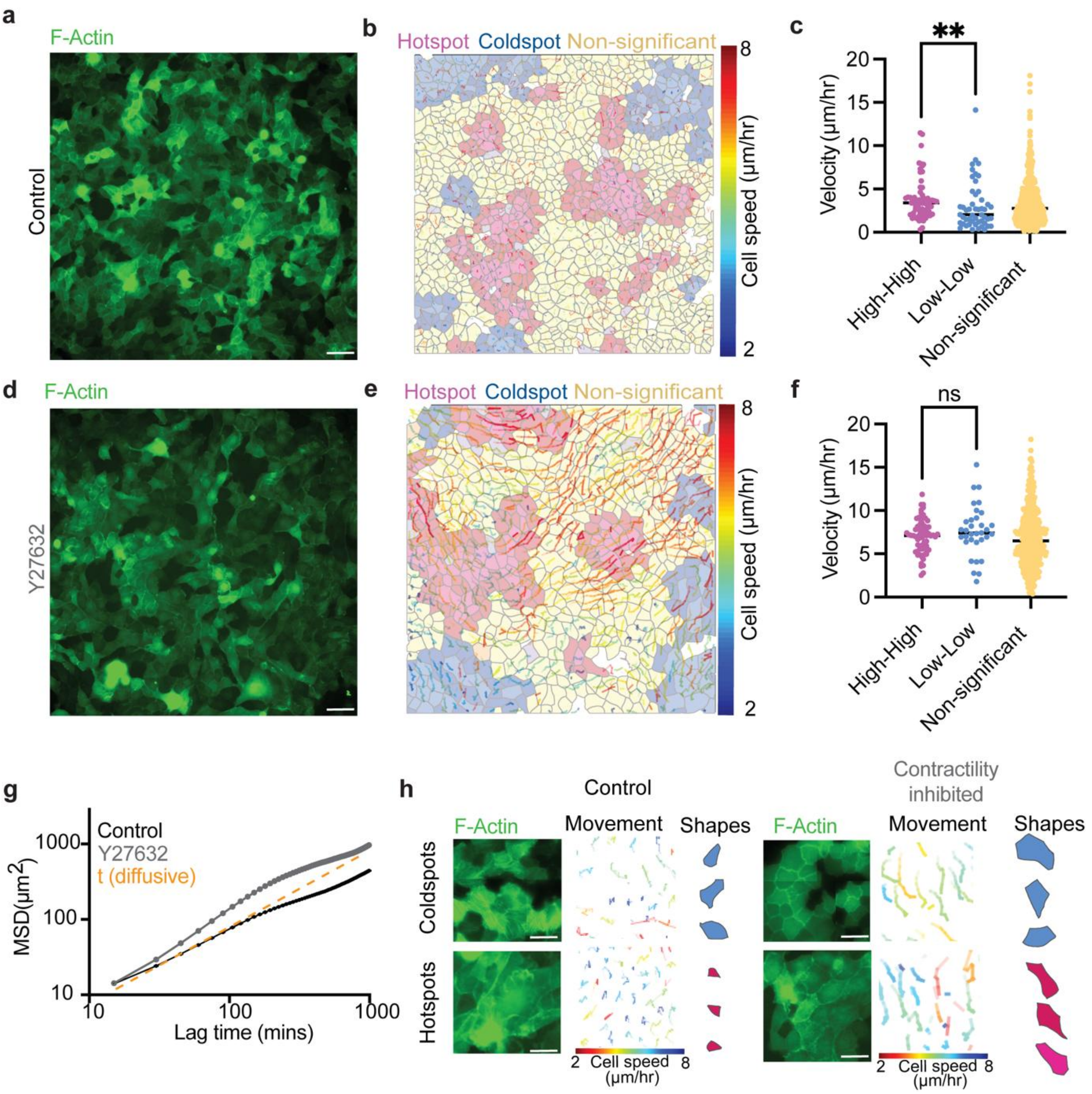


***Supplementary Figure 4: Disrupting contractility leads to loss of mechanochemical feedback and fluidises the tissue:*** *a) LifeAct MDCK cells from a representative control sample at the first timepoint, b) LISA cluster map corresponding to a and cell tracks overlaid on top, c) Comparison of cellular velocity at the different Actin LISA clusters, d) LifeAct MDCK cells from a representative Y27632 treated sample at the first timepoint, e) LISA cluster map corresponding to f and cell tracks overlaid on top, f) Comparison of cellular velocity at the different Actin LISA clusters, g) Mean Square Displacement plots for control and contractility inhibited monolayers, h)* Left/top- *High-resolution images of F-Actin at a representative coldspot from control sample, cell tracks and cell shapes;* Left/ bottom- *High-resolution images of F-Actin at a representative hotspot, cell tracks and cell shapes ;* Right/ top- *High-resolution images of F-Actin at a representative coldspot from Y27632 sample, cell tracks and cell shapes;* Right/ bottom- *High-resolution images of F-Actin at a representative coldspot from control sample, cell tracks and cell shapes.*

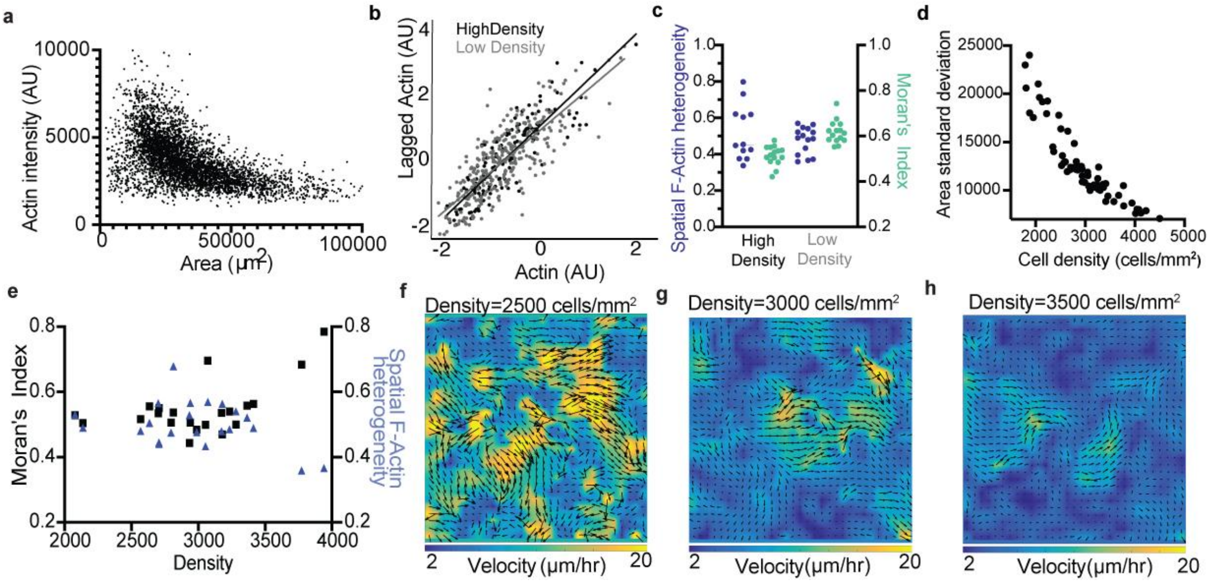


***Supplementary Fig 5: Cytoskeletal changes and Moran analysis of loosely and densely packed monolayers:*** *f) Plot of Actin vs Area at low and high density, g) Moran's plot of Actin vs lagged Actin at low and high density, h) Spatial F-Actin heterogeneity (percentage of Non-significant cells) and Global Moran's Index at high and low density, i) Standard deviation of cell area vs density, j) Spatial F-Actin heterogeneity and Moran's Index vs density, k-m) Velocity heatmap from Particle Image Velocimetry map at low (k), medium (l) and high density (m). Scale bars= 50µm. Lines represent the median.*

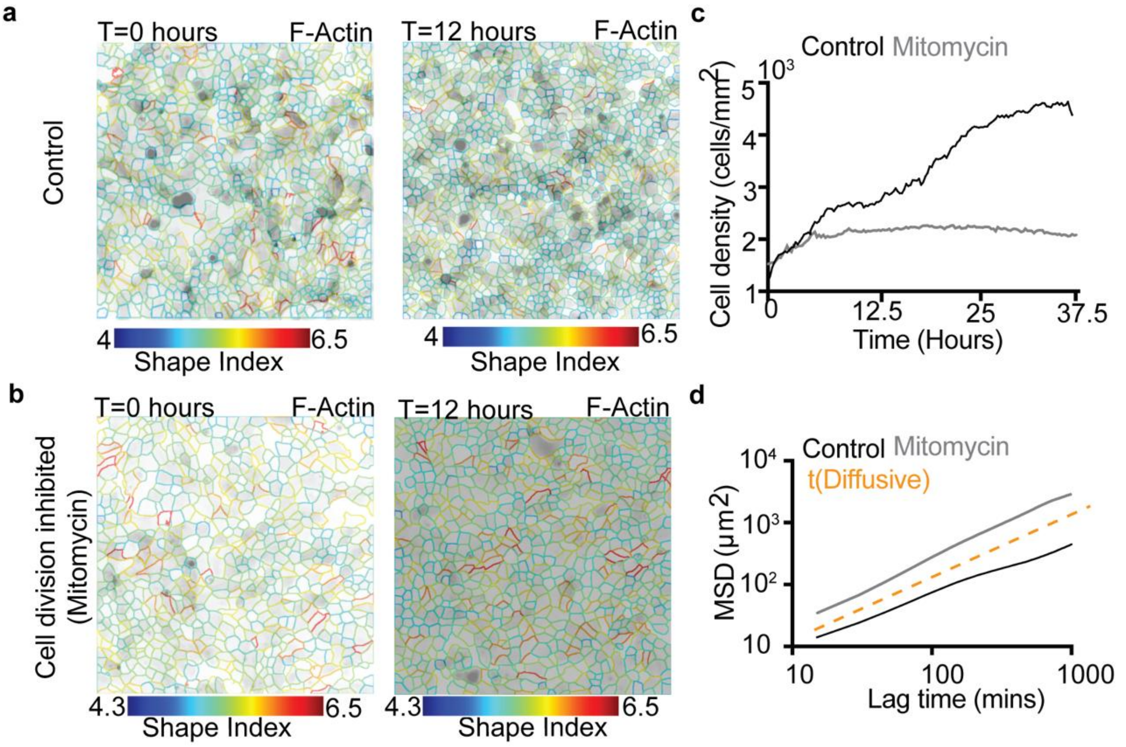


***Supplementary Figure 6: Fluidisation in the absence of crowding:*** *a) Timelapse images at T=0 hours (left) and T=12 hours (right) from a control sample, with cell outlines pseudocolored for Shape index, b) Timelapse images at T=0 hours (left) and T=12 hours (right) from a Mitomycin-C treated sample with cell outlines pseudocolored for Shape index, c) Plot of cell density over time for control and Mitomycin-C treated samples, d) Mean Square Displacement plots for control and Mitomycin-C treated samples*

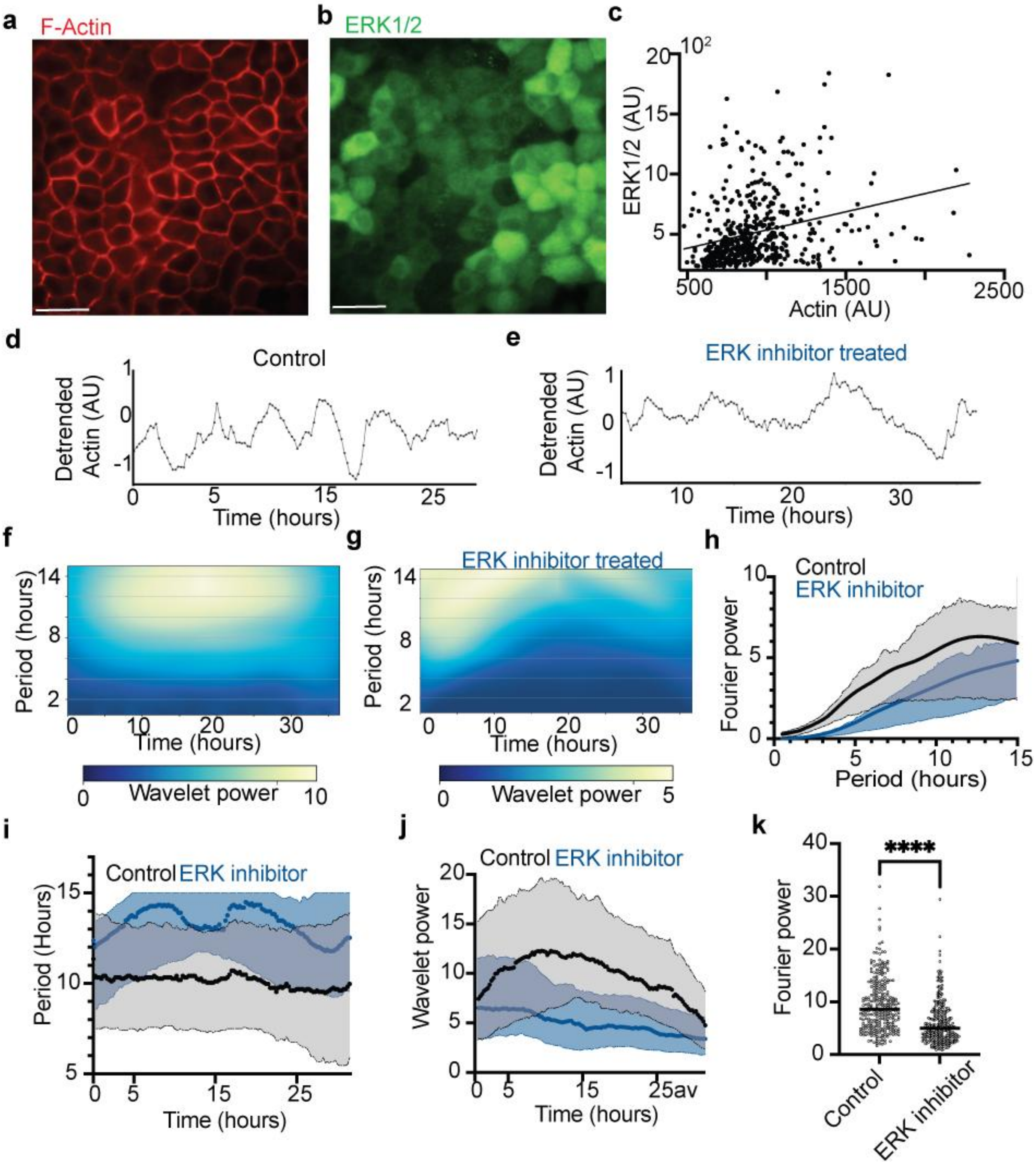


***Supplementary Fig 7: ERK inhibition leads to loss of Actin oscillations:*** *a) Representative image of monolayer stained for F-Actin (a) and ERK1/2 (b), c) Per-cell F-Actin plotted against per cell P-ERK1/2, d) Representative local Actin signal from a control monolayer, e) Representative local Actin signal from ERK inhibitor treated monolayer, f) Wavelet heatmap of control Actin signal, g) ) Wavelet heatmap of ERK-inhibitor treated Actin signal, h) Trend of power over time, i) Period determined by wavelet analysis as a function of time in the control (black) and ERK inhibitor treated (blue) Actin signals, j) Wavelet power as a function of time in the control (black) and ERK inhibitor treated (blue) Actin signals, k) Comparison of Fourier powers in the control and ERK inhibitor treated Actin signals. In h-j, lines represent the median and the shaded regions correspond to the quarters.*

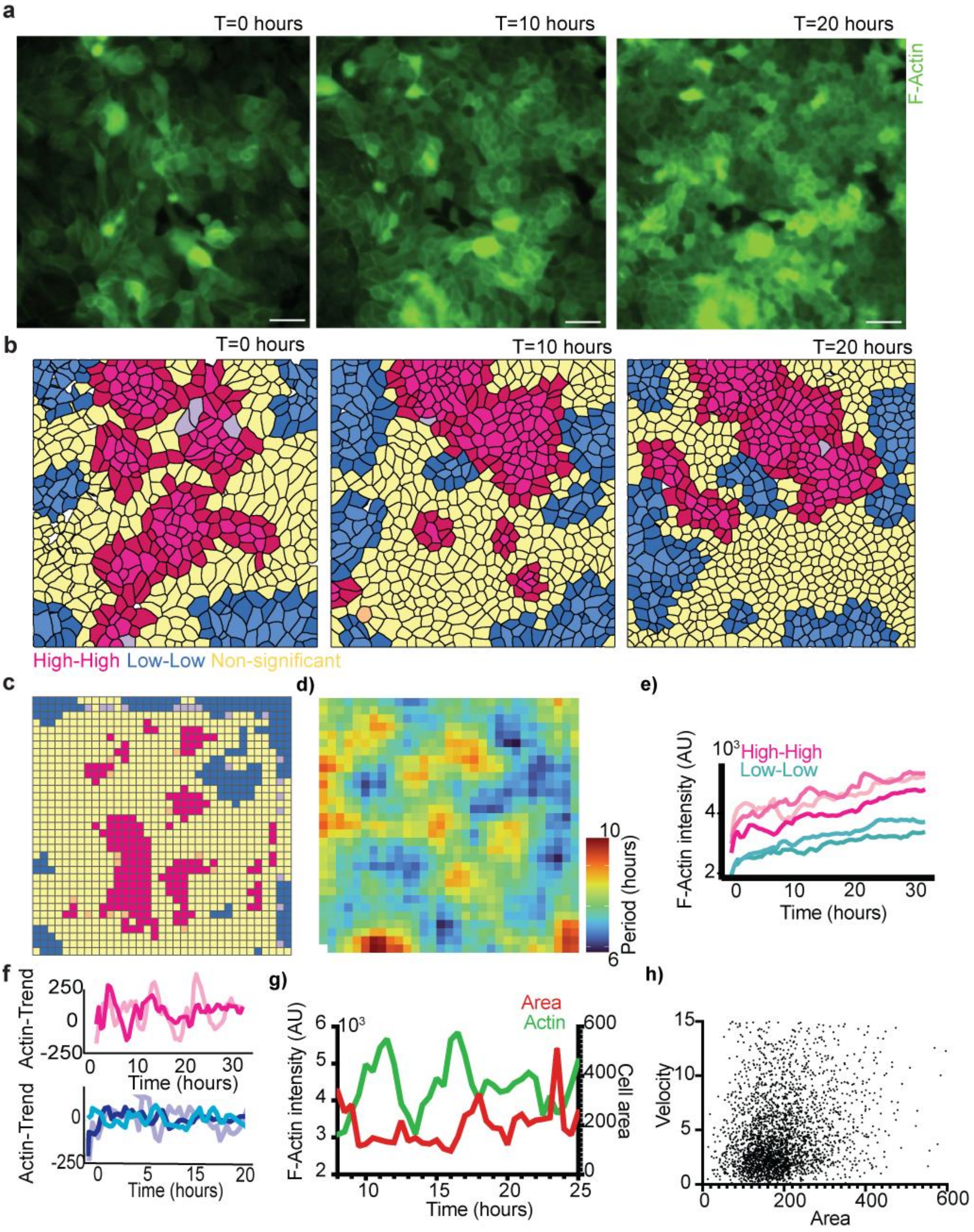


*Supplementary figure 8*: ***Hotspots and coldspots are stable over time and show local Actin oscillations with distinct periods that are spatially continuous with non-significant regions, related to area, velocity and local density fluctuations*****:**
*a) Representative timelapse images from homeostatic MDCK-LifeAct-GFP monolayers at 0 hours (left), 10 hours (middle) and 20 hours (right), b) LISA cluster maps of Actin at 0 hours (left), 10 hours (middle) and 20 hours (right), c) Representative LISA cluster map from the gridding analysis (methods section) used for Actin signal analysis, d) Heatmap of timeperiod at different locations in the monolayer demonstrating continuity in timeperiods, e) Actin signals without detrending from hotspots (pink) and coldspots (blue) as a function of time, f) Detrended Actin signals from hotspots (top) and coldspots (bottom) as a function of time, g)Single-cell Area (red- right y axis)and F-Actin (Green, left y-axis) obtained from cell segmentation and tracking plotted as a function of time, h) Single cell velocity plotted against single cell area, obtained from cell segmentation and tracking.*

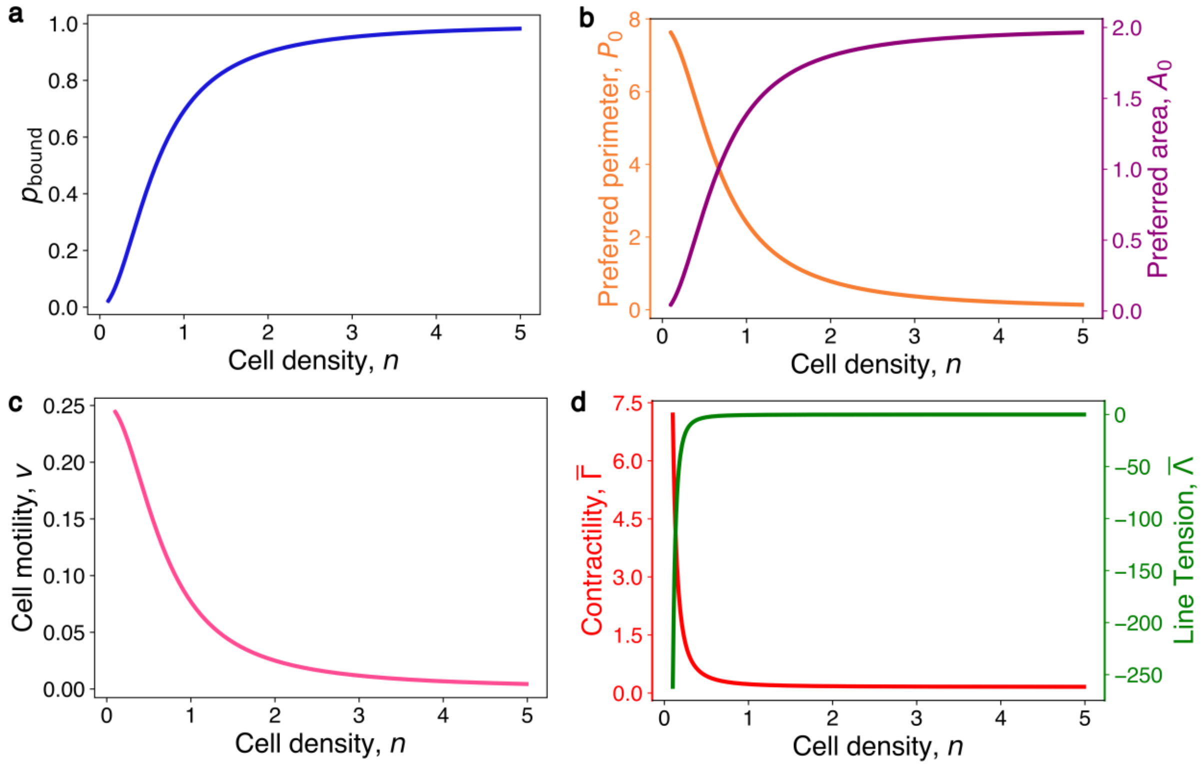


***Supplementary Fig 9: Mechanochemical feedback couples vertex model parameters to density:*** *a) Probability of bound cortical actin ($p_{bound}$) vs. density. b) The vertex model parameters $P_0$ and $A_0$ vs. density. c) Motility vs. density. d) Normalized contractility and normalized line tension vs, density in the model.*

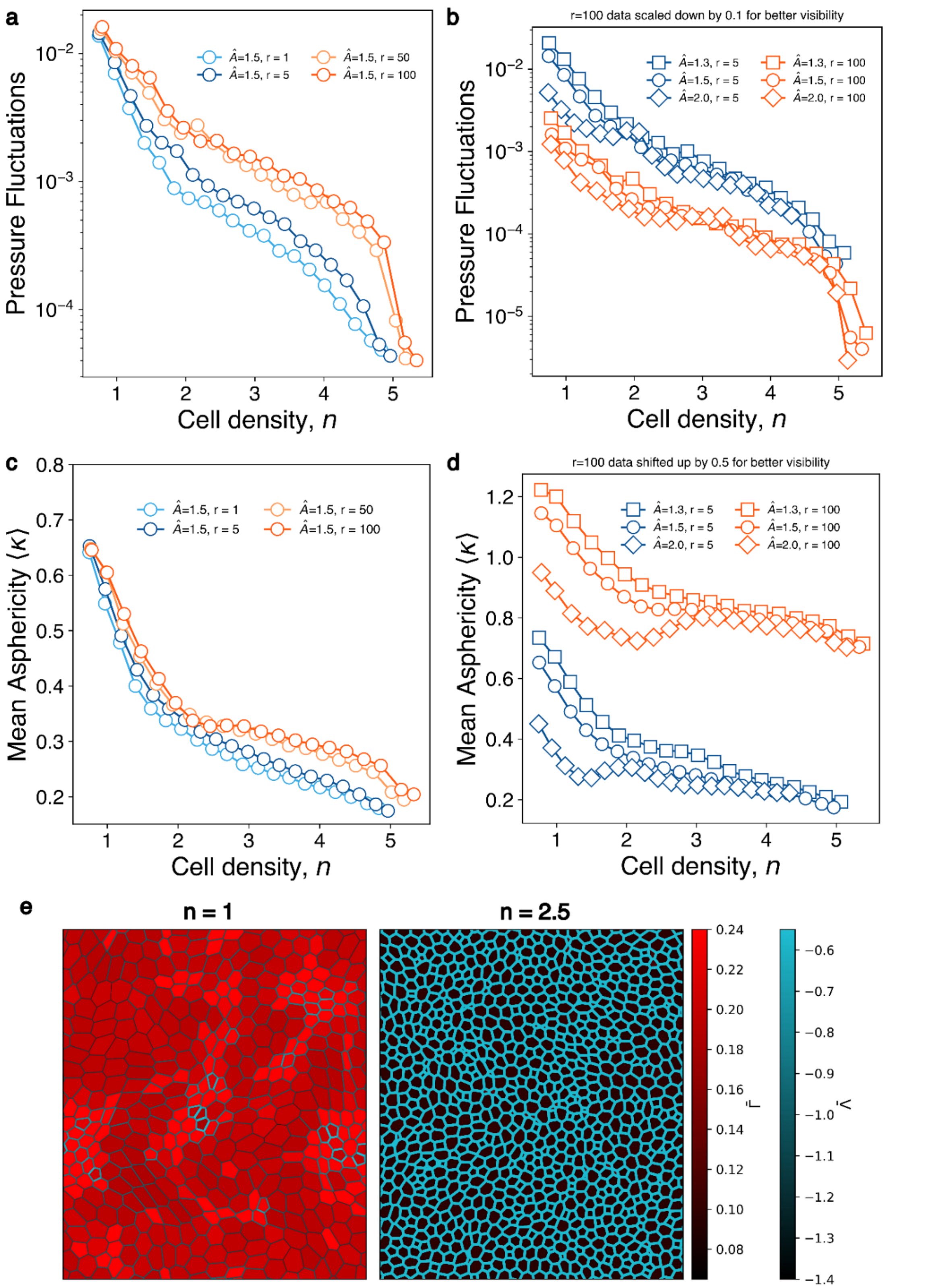


***Supplementary figure 10: Glass transition in the vertex model:*** *(a) Pressure fluctuations vs. density for* $\hat{A} = 1.5$ *shows rate dependence. (b) Pressure fluctuations vs. density for different* $\hat{A}$ *but one rate value. (c) Mean asphericity vs. density for* $\hat{A} = 1.5$ *shows clear rate dependence. (d) Mean asphericity vs. density for different* $\hat{A}$ *but one rate value. e) Contractility and line tension plots at different densities from a vertex model simulation with cell divisions and mechanochemical feedback.*

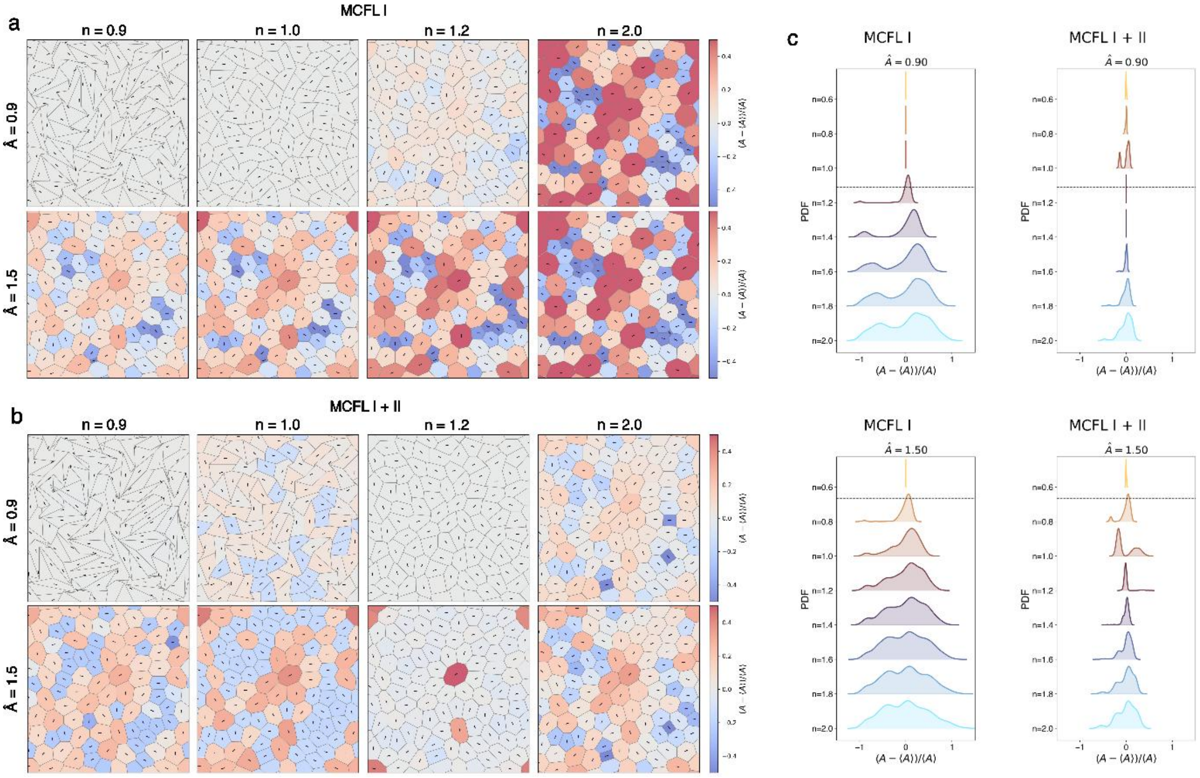


***Supplementary Fig 11: Constant density steady state configurations of the vertex model with mechanochemical feedback: a) Configurations with MCFL-I: At low , the configurations are floppy at , which progressively becomes regular at higher . For , the configurations are regular for most densities. The cells are coloured according to area strain b) Configurations with MCFL-I+II: The trend with increasing is similar to (a). c) Area strain distributions for configurations with MCFL-I and MCFL-I+II.***

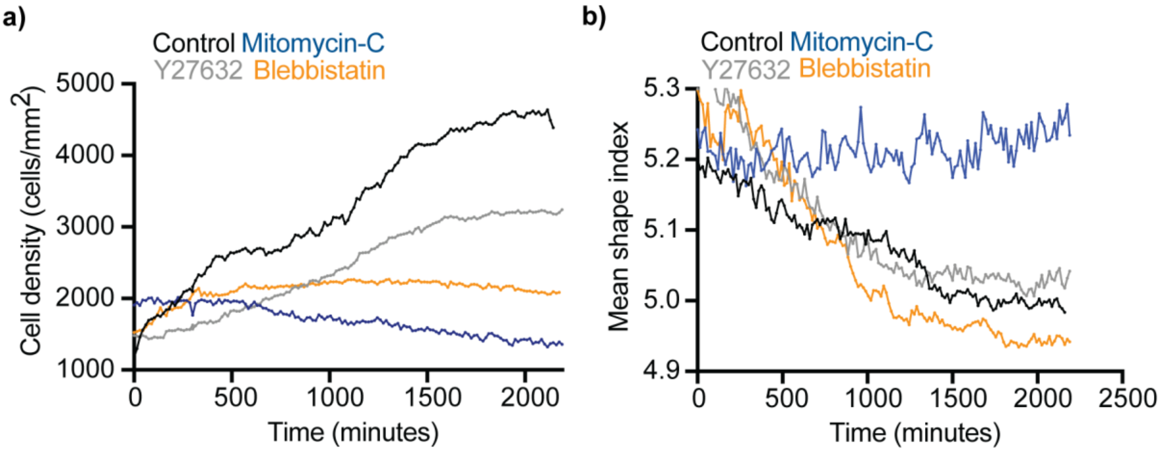


**Supplementary figure 12**: **Effect of drugs on division rate:** a) Plot of cell density over time in control, Mitomycin-C, Blebbistatin and Y27632 treated monolayers as a function of time, b) Change in division rate is also reflected in the mean shape index, as observed in the plot of Mean shape index over time in control, Mitomycin-C, Blebbistatin and Y27632 treated monolayers

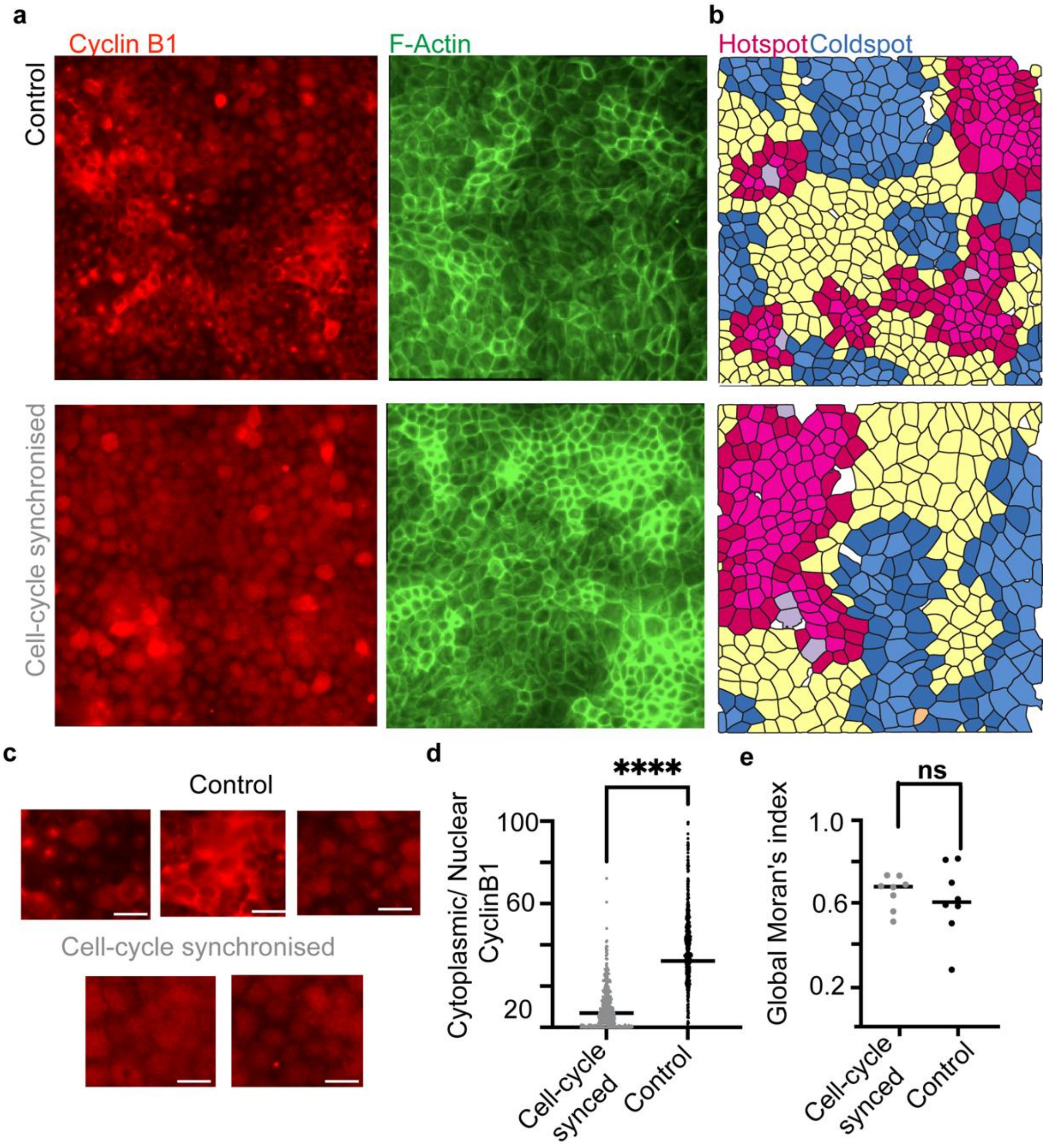


**Supplementary figure 13**: **Spatial Actin heterogeneity persists after cell cycle synchronization with Thymidine double block:** a) Confluent dishes of MDCK cells stained Cyclin B1 (Left-red) and F-Actin (Right green in cell-cycle synchronized by Thymidine double block (Bottom) and control (Top) samples, b) F-Actin LISA clusters maps corresponding to the images in panel A. c) Zoomed in images of Cyclin B1 corresponding to images in panel A demonstrating nuclear, cytoplasmic, perinuclear distribution of Cyclinb1at different locations in a control monolayer (top), compared to a more uniform distribution in nuclear or cytoplasmic compartments in the cell-cycle synchronized samples. d) Ratio of Cyclin B1 intensity at the nucleus and cytoplasm shows lesser variance in the cell-cycle synchronised samples suggesting uniformity in their cell-cycle state and e) No significant difference in the Global Moran's Index of Control and Cell cycle synchronized samples suggesting that inhomogeneous cell cycle state does not contribute to the observed heterogeneity in Actin levels. Lines represent the median. Scale bars=50μm.

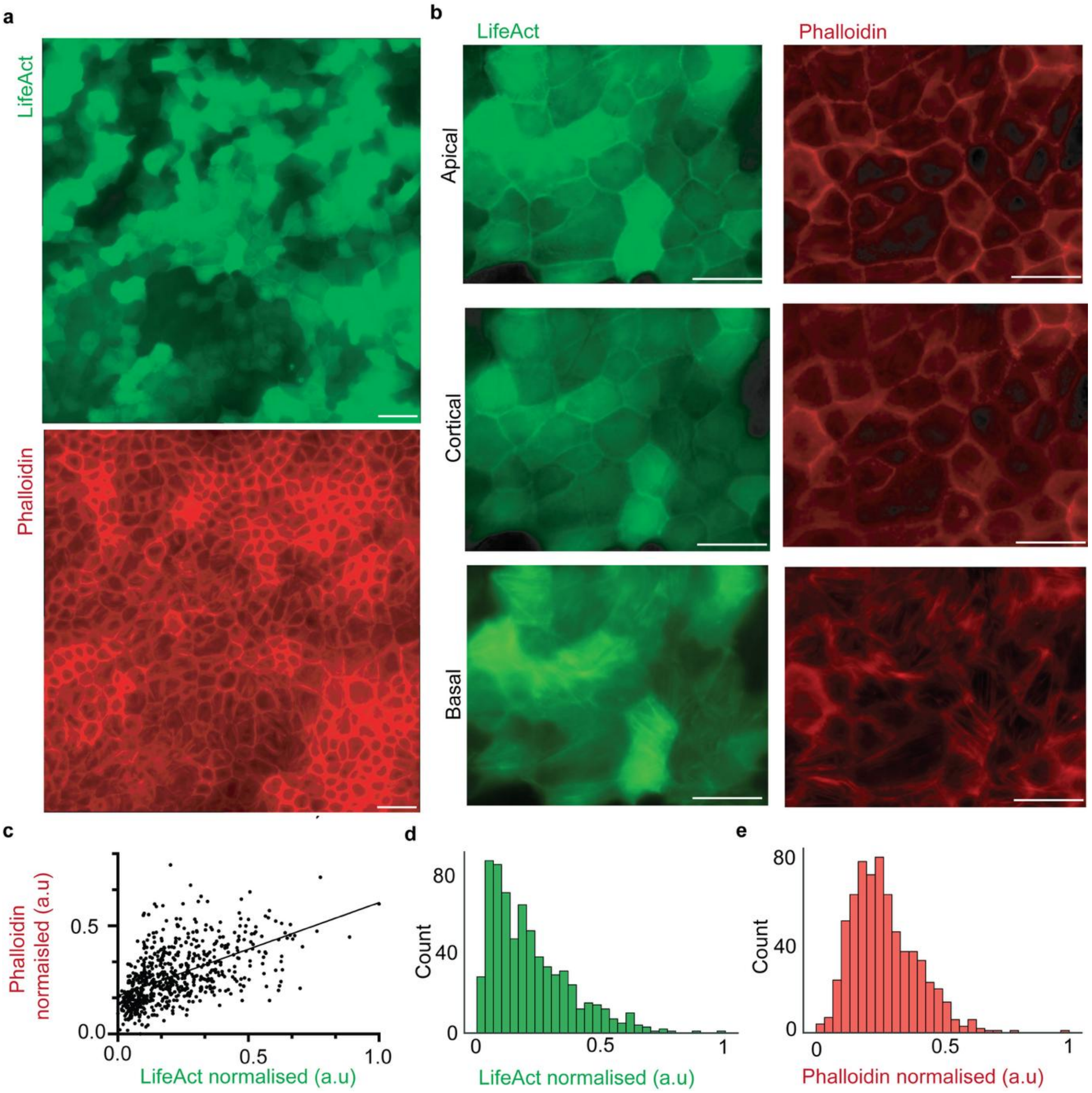


**Supplementary figure 14: Comparison of Actin staining with LifeAct and Phalloidin shows similar distributions:**

Confluent dishes of MDCK cells stained both for LifeAct-GFP (top) and Phalloidin 594 (bottom) imaged in 3D Z-stacks b) Zoomed images of LifeAct (Left-green) and Phalloidin (Right-red) at the top plane (Apical- top), middle plane (Cortical- middle), and bottom plane (Basal plane- bottom), c) Scatter plot of Min-Max Normalised LifeAct and Phalloidin intensities; $R^2$= 0.2925, d) Histogram of LifeAct and e) Phalloidin intensities. Scale bars=50µm.

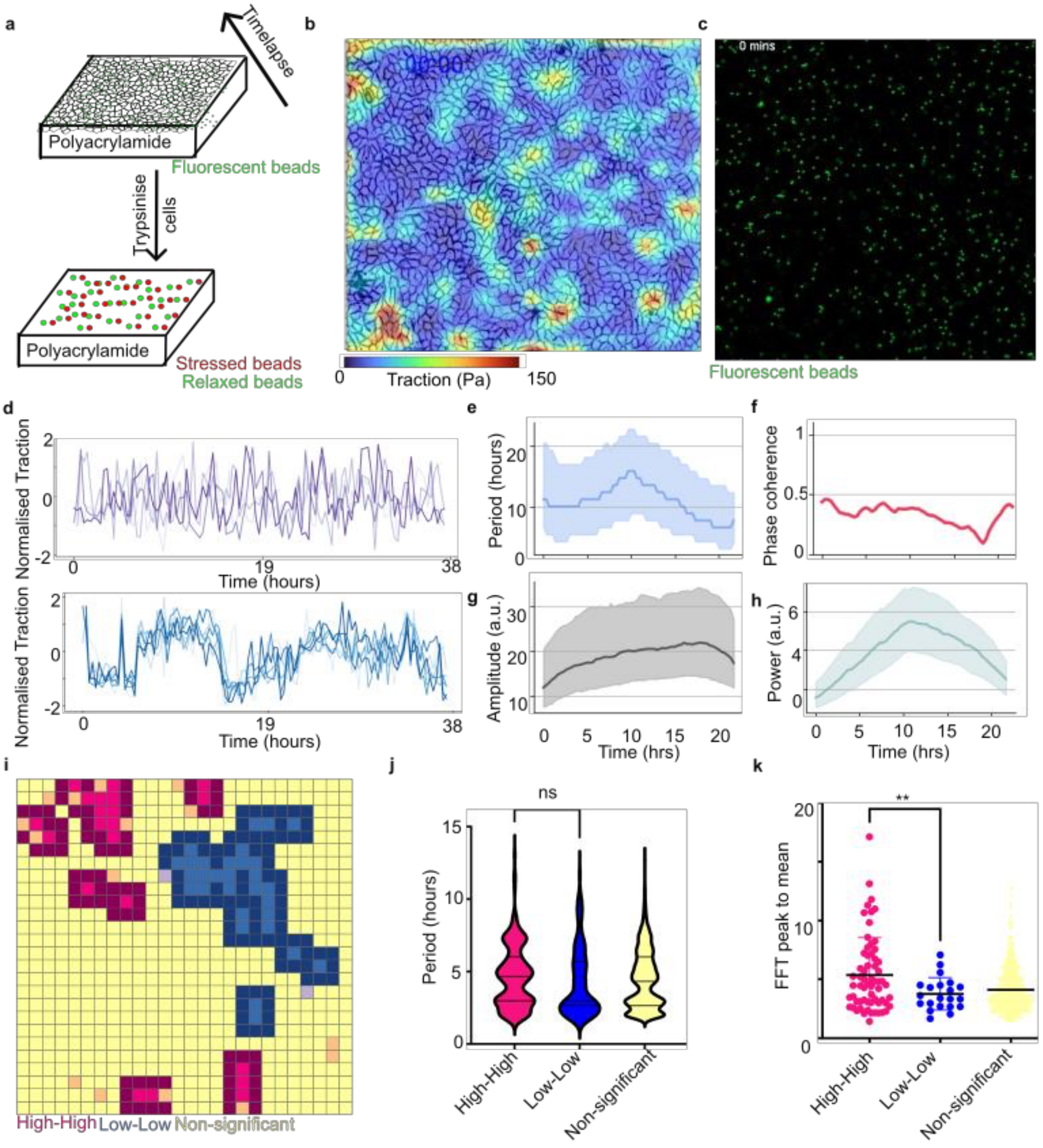


**Supplementary Figure 15: Spatiotemporal dynamics of traction force shows hours-scale oscillations**: a) Schematic of timelapse traction force Microscopy experiment, b) Traction force heatmap overlaid on cell image, c) Image of beads used in traction force experiment, d) Bottom- Representative locally oscillatory traction signals and top- Representative locally non- oscillatory traction signals, e) Period of oscillation obtained from Wavelet transform using PyBoat, f) Phase coherence over time obtained from Wavelet transform using PyBoat, g) Amplitude of traction signal over time obtained from Wavelet transform using PyBoat, h) Wavelet power over time obtained from Wavelet transform using PyBoat, i) LISA plot of traction force at T=20 hours, j) Plot comparing period of hotspots, coldspots and non-significant regions, k) Plot comparing FFT (Fast Fourier Transform) peak/mean ratio, a metric of oscillation strength at hotspots, coldspots and non-significant regions. Lines represent the median.

**Supplementary Video S1**: Timelapse movie of homeostatic LifeAct MDCK cells (left) with cell segmentation and tracking (right) used for MSD and Q(t) analysis, imaged every 2.5 minutes. Cell outlines are pseudocoloured for cell area, corresponding to the colourbar in the video. Cell tracks are coloured based on the average track speed, with the cell speed linearly increasing from 2 to 10 μm per hour from blue to red colour. Dark spaces correspond to cells that were not identified during segmentation. Scale bar=50μm.

**Supplementary Video S2**: Timelapse movie of homeostatic control LifeAct MDCK monolayer (left) and monolayer treated with Mitomycin C (right) to inhibit cell division (right).

**Supplementary video S3:** Timelapse movie of homeostatic LifeAct MDCK cells pseudocoloured for per-cell Actin intensity that were used to analyse the timeperiod of actin clusters. Dark spaces correspond to cells that were not identified during segmentation. Scale bar=50μm.

**Supplementary video S4:** Timelapse LISA segmentation of Actin intensity- LISA analysis performed for homeostatic LifeAct MDCK cells imaged every 30 minutes. The colours corresponding to LISA classification scheme are as follows: Pink- High-High, Yellow- Non-significant and Blue- Low- Low. Dark spaces correspond to cells that were not identified during segmentation. Scale bar=20μm.

**Supplementary video S5:** Timelapse movie of homeostatic single cell LifeAct-MDCK cells without cell-cell contacts imaged every two minutes, that were used for the analysis of Actin oscillation period in single cells.

**Supplementary video S6:** Cell tracking of low (left) and high (right) density monolayers compared side-by-side. The final timepoint shows the tracks for the cells over all timepoints. Cells are pseudocoloured for cell area, corresponding to the colourbar in the video. Cell tracks are coloured based on the average track speed, with the cell speed linearly increasing from 2 to 10 μm per hour from blue to red colour. Scale bar=50μm.

**Supplementary Video S7**: Cropped regions showing Actin oscillations in neighbouring patches. Cell outlines are pseudocoloured for cell area, corresponding to the colour bar below

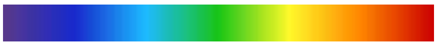

200 Cell Area (μm$^2$) 1000

**Supplementary Video S8:** Experimental videos depicting oscillations of different periods at the actin hotspots and coldspots. Left- Evolution of LISA clusters over time, Middle- Zoomed in view of cells from the coldspots (topleft) and hotspots (bottom left) and corresponding segmented and tracked cells pseudocoloured for Actin intensity (right). Right- Representative detrended Actin signals from coldspots (bottom) and hotspots (top). Scale bar=50μm.

**Supplementary Video S9:** Video showing vertex model simulation with cell divisions: i) without mechanochemical feedback, ii) with only MCFL-I, iii) with MCFL-I+II.

**Supplementary Video S10:** Vertex model simulation with cell divisions and MCFL-I+II. The colormap shows the local cell density or 1/ (cell area) and the arrows indicate the PIV or the average velocity around that grid point.

### Supplementary Information: Theory

Sindhu Muthukrishnan,[1] Phanindra Dewan,[2] Tanishq Tejaswi,[1] Michelle B Sebastian,[1] Tanya Chhabra,[1] Soumyadeep Mondal,[2] Soumitra Kolya,[3] Sumantra Sarkar,[2, *] and Medhavi Vishwakarma[1, †]

[1] *Department of Bioengineering, Indian Institute of Science, Bengaluru, Karnataka, India, PIN 560012*
[2] *Department of Physics, Indian Institute of Science, Bengaluru, Karnataka, India, PIN 560012*
[3] *Tata Institute of Fundamental Research, Hyderabad, India, PIN 500046*
(Dated: March 17, 2026)

#### I. RATIONALE BEHIND THE MODEL AND THE CONSTITUTIVE RELATIONS

##### A. The canonical vertex model

The canonical vertex model is described by the following hamiltonian [1, 2]:

$$H = \sum_{\alpha} \frac{1}{2} K (A_\alpha - A_0)^2 + \sum_{\alpha} \frac{1}{2} \Gamma P_\alpha^2 + \sum_{i,j} \Lambda_{i,j} l_{i,j}, \tag{S1}$$

where $A_\alpha$, $P_\alpha$ are the area and perimeter of the cell $\alpha$. $K, \Gamma$, and $A_0$ are the area elasticity coefficient, perimeter elasticity coefficient, and preferred area of the cells, respectively, which we take to be identical for all the cells. $l_{ij}$ is the length of the interface or edge between vertices $i$ and $j$, and $\Lambda_{ij}$ is the line tension along this interface. If we assume that the line tension is the same across all the interfaces, such that $\Lambda_{ij} = \Lambda$, the vertex model can be written in another well-known canonical form:

$$H = \sum_{\alpha} \frac{1}{2} K (A_\alpha - A_0)^2 + \sum_{\alpha} \frac{1}{2} \Gamma (P_\alpha - P_0)^2, \tag{S2}$$

where $P_0 = -\Lambda/\Gamma$. The parameters of the vertex model can be normalized to define contractility, $\bar{\Gamma}$ and the normalized tension, $\bar{\Lambda}$ as follows:

$$\begin{aligned} \bar{\Gamma} &= \left(\frac{\Gamma}{K}\right) \frac{1}{A_0} \\ \bar{\Lambda} = \frac{\Lambda}{K A_0^{3/2}} &= -\left(\frac{\Gamma}{K}\right) \frac{P_0}{A_0^{3/2}} \end{aligned} \tag{S3}$$

*a. Ground state phase diagram:* The ground state of the vertex model has been analyzed from which a phase diagram can be constructed (Fig. T1). We use this phase diagram as a starting point for our analysis.

##### B. Biological origin of contractility and tension

The actin cytoskeleton of the cell controls the contractility of the cell and the line tension of cell-cell contacts in epithelial tissues. However, they originate from different morphologies of the actin cytoskeleton. Contractility controls the length of the perimeter, and it arises from the formation of contractile actomyosin rings and stress fibers. In contrast, the line tension arises from the accumulation of cortical actomyosin along the cell-cell junctions. Therefore, we can assume that high line tension implies higher accumulation of the cortical actin, whereas high contractility implies higher accumulation of the stress fibers and the contractile rings. Translating these observations into the parameters of the vertex model (Eq. S3), we can construct the following picture (Fig. T2). This picture implies that for fixed $\Gamma/K$, increasing $A_0$ decreases the stress fibers and increases the junctional actins. Similarly, an increase in line tension is associated with decreasing $P_0$ or increasing $A_0$ or both.

As a tissue matures through cell division and cell-cell interactions, we expect it to explore some trajectories in this phase diagram. In our experiments, the tissue solidifies through cell division. Time-lapse imaging of actin, myosin,

---

*
†

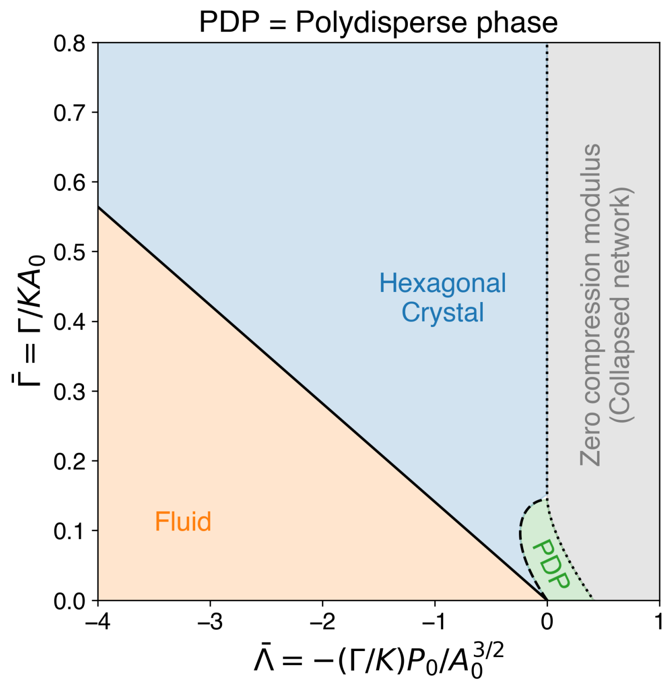


FIG. T1. The ground state phase diagram of the canonical vertex model. The solid and the dashed black lines denote regions where the shear modulus of the hexagonal crystal vanishes. The orange fluid state is composed of soft networks, whereas the green polydisperse phase is composed of polygons of various shapes and sizes. In the canonical vertex model, the PDP can contain 4-8 or 3-12 crystalline lattices.

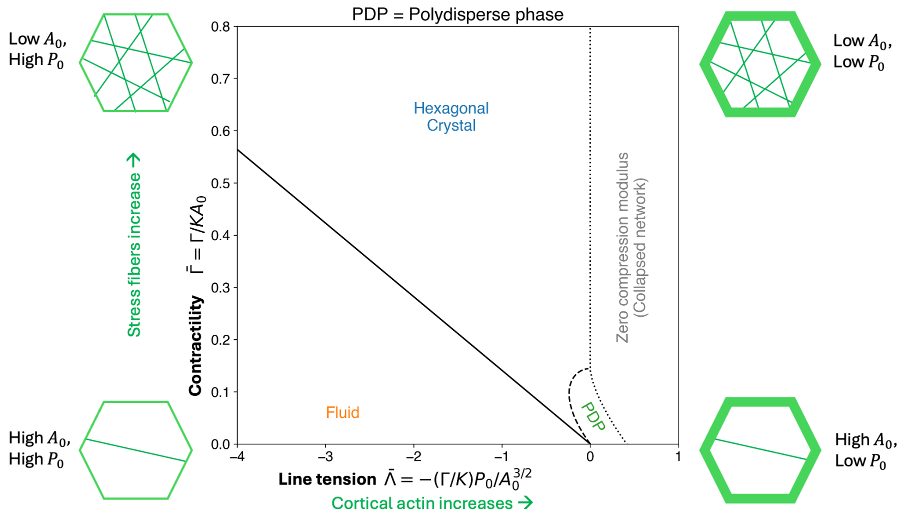


FIG. T2. The biological origin of the contractility and tension.

and E-cadherin during this solidification process reveals that the cells contain many elongated stress-fibers and low accumulation of junctional actin at low densities. The opposite trend is seen at high density, in which the cells have highly mature cell-cell junctions with high accumulation of actin, myosin, and E-cadherin at the junctions. In contrast, the actin is highly depleted in the bulk, and stress fibers are hardly seen. Similar observations were also reported earlier. Hence, during tissue solidification through cell division, the trajectory should start at the upper left part of the phase diagram and move to the lower right part, as shown in the figure below (Fig. T3).

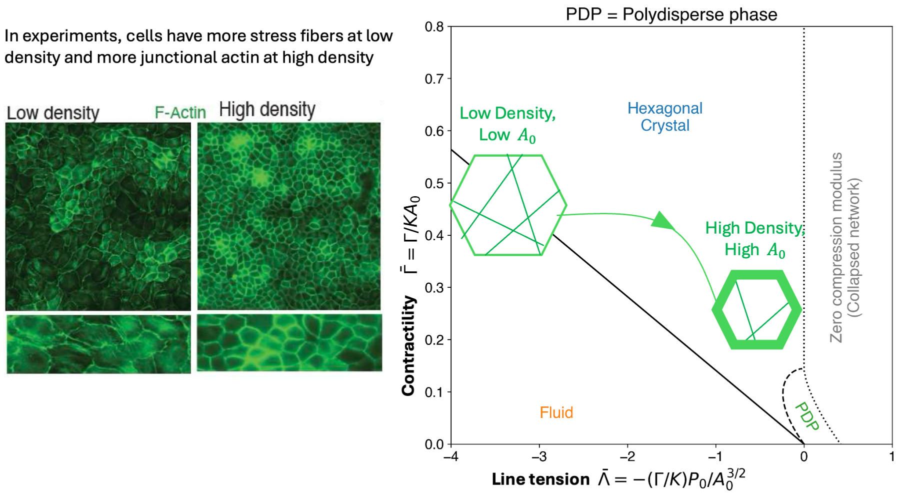


FIG. T3. Trajectory of our experiment on the phase diagram. A theoretical description of tissue solidification through cell division will require a similar change in the contractility and line tension.

#### C. Constitutive laws for tissue solidification through cell division

Based on the analysis above, it is clear that any model of crowding-induced tissue solidification, such as through cell division, would require constitutive relations that ensure high contractility & low tension at low density and low contractility & high tension at high density. One way to achieve this phenomenology is to consider the load-dependent binding of myosin to actin in the cell-cell junction. As cell density increases, cells touch each other and form adhesive interfaces. This affects cell mechanics in two ways. First, the density-dependent binding of E-cadherin catalyzes the production of Rho-GAP [3], which inhibits Rho-dependent actomyosin contractility in the cell cytoplasm, inhibiting the formation and the maintenance of the stress fibers [4]. Second, E-cadherin, recruits Rho at the cell-cell junction, increasing the actomyosin contractility through a density-dependent positive feedback [5]. The abrogation of the stress fibers in the cytoplasm pays for the enhanced junctional/cortical actomyosin. We can capture the fraction of cortical actomyosin, $p_{bound}$, as a function of cell density from this consideration.

##### 1. *Load-dependent binding of myosin*

The unbinding rate of myosin from actin depends on the local strain. The higher the contraction of the actomyosin network, the lower the unbinding rate. In fact, to a good approximation, the unbinding rate, $k_u$, decays exponentially with the strain, $\epsilon$. Specifically, for an isotropic deformation of the cell in the 2D vertex model, the strain is a scalar:

$$\epsilon = \frac{A - \hat{A}}{\hat{A}}, \tag{S4}$$

where $\hat{A}$ is a reference area, whose form will be determined later. The unbinding rate can be written as [6, 7]:

$$k_u = k_u^0 \exp(\chi\epsilon) \tag{S5}$$

$$= k_u^0 \exp\Big(\chi(A/\hat{A} - 1)\Big) \tag{S6}$$

$$\equiv k_u^0 \exp(\chi(x - 1)) \tag{S7}$$

$$x = A/\hat{A} \tag{S8}$$

Hence, for $\chi > 0$, which is what is observed experimentally [6, 7], the unbinding rate decreases with increasing contraction, $x < 1$, leading to increased actomyosin binding, which increases the contraction further. This mechanochemical feedback loop, which we call MCFL-I, changes the bound fraction of actomyosin in a density-dependent manner. To see this, consider the following reaction:

$$M + F \underset{k_b}{\overset{k_u}{\rightleftharpoons}} MF \tag{S9}$$

Here $M$ and $F$ denotes myosin and F-actin, respectively, and $MF$ is their bound state. Assuming that the reactions occur at a much faster rate than appreciable changes in the cell area, the bound fraction of myosin can be calculated assuming chemical equilibrium.

$$p_{bound} = \frac{[MF]}{[M]+[MF]} \tag{S10}$$

In chemical equilibrium,

$$k_b[M][F] = k_u[MF] \tag{S11}$$

$$\therefore p_{bound} = \frac{1}{1+\frac{k_u}{k_b[F]}} \tag{S12}$$

$$= \frac{1}{1+\frac{k_u^0}{k_b}\frac{\exp[\chi(x-1)]}{[F]}} \tag{S13}$$

$$= \frac{1}{1+K_D^0\frac{\exp[\chi(x-1)]}{[F]}} \tag{S14}$$

The equation simplifies significantly when $\chi = 1$, and $x \approx 1$. In this limit,

$$p_{bound} \approx \frac{1}{1+\frac{K_D^0}{N_F/A}x} \tag{S15}$$

$$\tag{S16}$$

where $N_F = [F]A$ is the number of f-actin in the cell. Our experiments show that the total amount of actin remains constant. Hence, $K_D^0/N_F$ is a constant. In fact, we define:

$$\hat{A} = N_F/K_D^0. \tag{S17}$$

Clearly, $\hat{A}$ is inversely proportional to the dissociation constant of the actomyosin complex. Hence, it can be controlled by controlling the binding affinity of the myosin. In general, the lower the dissociation constant, the higher the $\hat{A}$, and hence the higher the binding affinity of myosin to actin. With this definition, we can write the bound fraction as:

$$p_{bound} = \frac{1}{1+\left(\frac{A}{\hat{A}}\right)^2} = \frac{1}{1+x^2} \tag{S18}$$

Eq. S18 works remarkably well for a range of area, $A$ (Fig. T4). Furthermore, this formula is much easier to work with in analytical calculations than the more general equation:

$$p_{bound} = \frac{1}{1+x\exp\{[\chi(x-1)]\}}. \tag{S19}$$

Hence, we use this equation for all our simulations and analytical calculations.

##### 2. Constitutive relations

The line tension, $\bar{\Lambda}$ should be proportional to the bound fraction. Because $\bar{\Lambda} \propto -P_0$, $P_0$ should decrease with $p_{bound}$, the fraction of junctional actin and increase with $1-p_{bound}$, the fraction of stress fibers. $p_{bound}$ increases with decreasing area, that is, increasing density. Hence, the fraction of stress fibers decreases with increasing density. This reduction in stress fiber arises from the conservation of the total actin in the cell and allocation of more F-actin into the cell-cell junctions as the density increases. This observation implies that contractility, $\bar{\Gamma}$, should decrease with increasing $p_{bound}$. Finally, because $\bar{\Gamma} \propto 1/A_0$, $A_0$ should increase with $p_{bound}$. Taken together, we propose the following constitutive relations:

$$\begin{aligned} A_0(A) &= 2a_0\frac{1}{1+\left(\frac{A}{\hat{A}}\right)^2} \\ P_0(A) &= 2\hat{q}_0\sqrt{a_0}\frac{\left(\frac{A}{\hat{A}}\right)^2}{1+\left(\frac{A}{\hat{A}}\right)^2}. \end{aligned} \tag{S20}$$

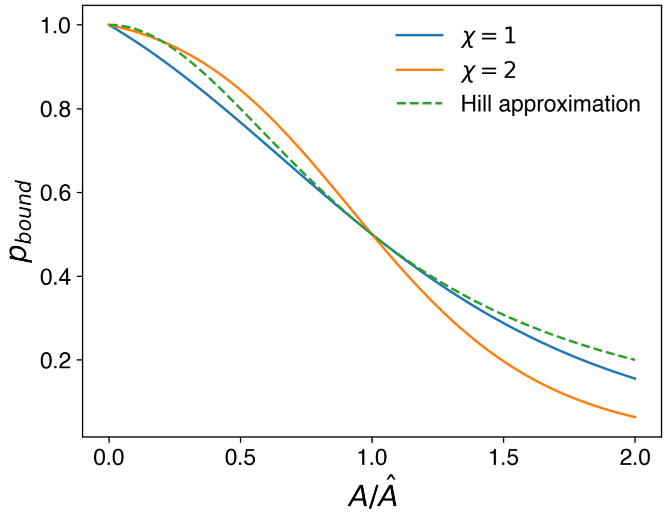


FIG. T4. Comparison of $p_{bound}$ computed from the true expression (Eq.S19) vs the hill approximation (Eq.S18).

Here,

$$a_0 = A_0(\hat{A}) = 1 \tag{S21}$$

$$\hat{q}_0 = P_0(\hat{A})/\sqrt{A_0(\hat{A})} \tag{S22}$$

$$\text{Cell Density, } n = 1/A \tag{S23}$$

These two constitutive relations are sufficient to reproduce the "static" properties of the tissue, such as its zero-frequency elastic moduli. To understand tissue dynamics, such as glass transition through cell division, we need to quantify the observed dependence of cell motility on the intensity of stress fibers. Specifically, it has been observed that the cell motility is directly proportional to the stress fibers present in the cell [8, 9]. Because in our model $(1 - p_{bound})$ is the fraction of stress fibers, we postulate the following constitutive relation relating cell motility to the fraction of stress fibers.

$$v(A) = 2v_0 \frac{\left(\frac{A}{\hat{A}}\right)^2}{1 + \left(\frac{A}{\hat{A}}\right)^2} \tag{S24}$$

This relation ensures that at high densities the motility is zero. Although we have used this specific relationship for the motility, any decreasing function of cell density works.

These constitutive relations, shown in Fig. T5, reproduce the experimentally observed behavior with fluid phase at low densities and solid phase at high densities, as shown in Fig. T6. In the next section, we will use them to construct the ground state phase diagram as a function of cell density. However, it is important to consider alternate constitutive relations, which we do next.

#### D. Alternate constitutive relations

##### 1. *Constant $A_0$ and $P_0$, but density-dependent $\Gamma/K$*

If $A_0$ and $P_0$ are constants, then a simple linear relationship exists between $\bar{\Gamma}$ and $\bar{\Lambda}$:

$$\bar{\Lambda} = -\left(\frac{\Gamma}{K}\right)\frac{P_0}{A_0^{3/2}} = -\frac{P_0}{\sqrt{A_0}}\frac{\Gamma}{KA_0} \tag{S25}$$

$$\therefore \bar{\Lambda} = -\hat{q}_0\bar{\Gamma}. \tag{S26}$$

The line separating the fluid region from the hexagonal crystal in Fig. T1 satisfies the equation:

$$\bar{\Lambda} = -4\sqrt{\pi}\bar{\Gamma}. \tag{S27}$$

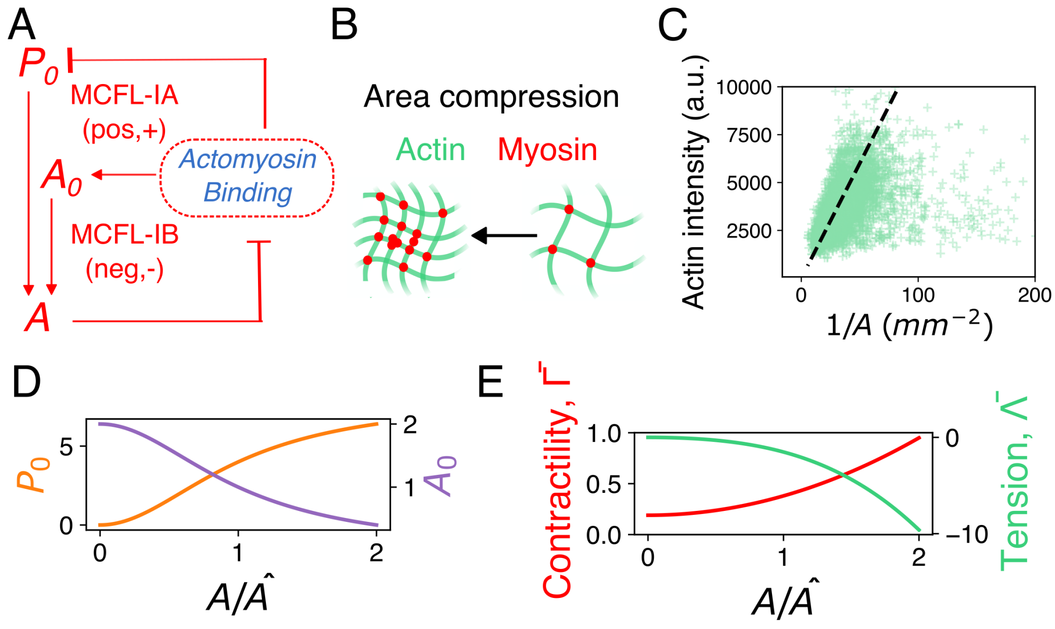


FIG. T5. (A)MCFL-I consists of two interacting positive and negative feedback loops. MCFL-IA leads to bistability, which creates coexisting cells with larger and smaller areas. MCFL-IB stabilizes the bistable areas from increasing or decreasing indefinitely, creating a stable distribution of cells even at high densities. (B) It arises from the load-dependent binding of myosin to actin. (C) Actin intensity is inversely proportional to the area of the cell, implying that the total actin is constant. (D) $A_0$ and $P_0$ as a function of $A/\hat{A}$ shows the density-dependence of the constitutive relations. (E) Contractility and tension as a function of $A/\hat{A}$ computed from these constitutive relations.

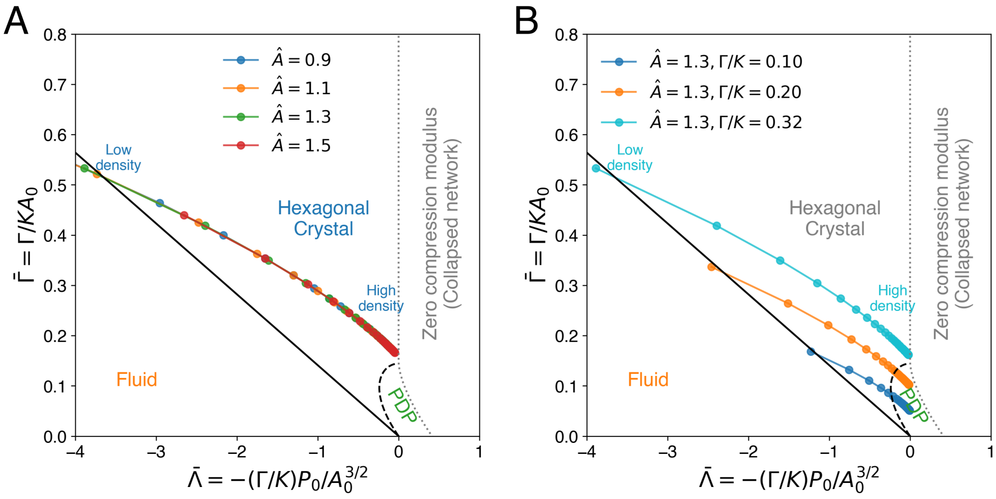


FIG. T6. (A)The constitutive relations (Eq. S20) reproduce density dependence consistent with experimental trajectories. Here, trajectories for a few $\hat{A}$ values are shown. (B) Changing $\Gamma/K$ shifts these trajectories along the $\bar{\Gamma}$ axis. For $\Gamma/K < 1/2\sqrt{3}$, the trajectories can access the PDP phase at high density. The rigidity of this state is yet to be determined.

Hence, as long as the shape parameter $\hat{q}_0 < 4\sqrt{\pi} \approx 7.1$, the trajectory always remains in the crystalline region. Conversely, if $\hat{q}_0 > 4\sqrt{\pi}$, then the trajectory strictly remains in the fluid region of the phase diagram. For sufficiently small $\hat{q}_0$, the trajectories in the crystalline phase can reach the PDP phase at small $\bar{\Gamma}$ values. Clearly, this protocol does not match experimental observation, where a single trajectory remains in the fluid phase at a low density and reaches the crystalline phase at high density. Doing so requires both $\bar{\Gamma}$ and $\hat{q}_0$ to change with density, in which case, they are equivalent to the constitutive relations Eq. S20.

##### 2. $A_0 \propto 1/n$, $P_0$ and $\Gamma/K$ constant

As cell density, $n$, increases, the area of the cells, $A$, decreases. The average area scales as $1/n$. One possible way to achieve this is to make the preferred area, $A_0 = 1/n$, while keeping all other parameters constant. Hence, in this protocol, the contractility, $\bar{\Gamma}$, increases with $n$, and the line tension decreases with $n$, leading to fluidization at high density. Furthermore, the trajectory is exactly opposite to what is observed in experiments, including ours. Hence, this constitutive relation is unphysical and should not be used to model a proliferating tissue.

#### E. Effect of Blebbistatin

With the assumption that the only way blebbistatin ($B$) can interact with myosin is via competitive inhibition, we can write the kinetic equations as:

$$M + F \underset{k_b}{\overset{k_u}{\rightleftharpoons}} MF \tag{S28}$$

$$M + B \underset{k_1}{\overset{k_2}{\rightleftharpoons}} MB \tag{S29}$$

At equilibrium, we get

$$k_b[F][M] = k_u[FM] \tag{S30}$$

$$k_1[B][M] = k_2[BM] \tag{S31}$$

We can calculate the fraction of unbound myosin as

$$\frac{[M]}{[FM]} = \frac{k_u}{k_b[F]} = \frac{k_D^F}{[F]} \tag{S32}$$

and,

$$\frac{[BM]}{[FM]} = \frac{k_u}{k_2}\frac{k_1[B]}{k_b[F]} = \frac{k_u}{k_b[F]}\frac{k_1[B]}{k_2} = \frac{k_D^F}{[F]}\frac{[B]}{k_D^B} \tag{S33}$$

And finally we can calculate the fraction of myosin bound to actin as

$$\frac{[FM]}{\text{Total M}} = \frac{[FM]}{[M+[FM]+[BM]]} \tag{S34}$$

$$= \frac{1}{1+\frac{[M]}{[FM]}+\frac{[BM]}{[FM]}} \tag{S35}$$

$$= \frac{1}{1+\frac{k_D^F}{[F]}+\frac{k_D^F}{[F]}\frac{[B]}{k_D^B}} \tag{S36}$$

$$= \frac{1}{1+\frac{k_D^F}{[F]}\left(1+\frac{[B]}{k_D^B}\right)} \tag{S37}$$

$$= \frac{1}{1+\left(\frac{A}{\hat{A}}\right)^2\left(1+\frac{[B]}{k_D^B}\right)} \tag{S38}$$

$$= \frac{1}{1+(A/\hat{A}')^2} \tag{S39}$$

where, $\hat{A}' = \frac{\hat{A}}{\sqrt{1+[B]/k_D^B}}$, is the modified $\hat{A}$. From this calculation we can see that the presence of blebbistatin can reduce $\hat{A}$.

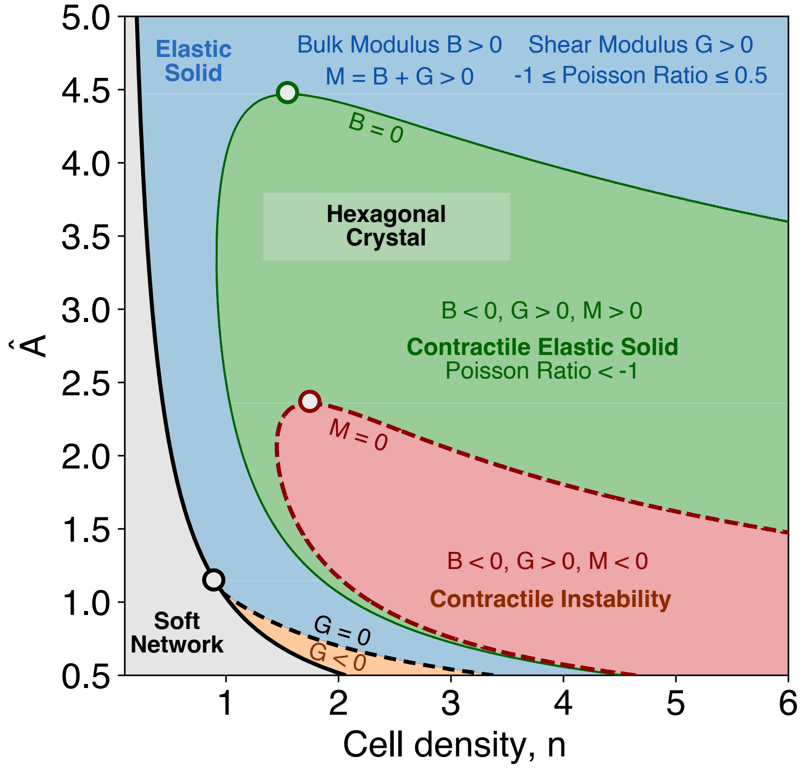


FIG. T7. The ground state phase diagram of the vertex model with MCFL-I in the absence of any motility. There are five different regions, marked by whether $q_0 = P_0/\sqrt{A_0} = 3.722$ (solid black line), which is also the jamming density $n_J$, and the elastic moduli of the crystalline ground state. In the gray region, $q_0 = P_0/\sqrt{A_0} > 3.722$, and the ground state is a floppy soft network. Above a density (the non-gray regions), $n_{lo}(\hat{A})$, marked by the solid black line, the ground state is a hexagonal crystal. In the light blue region, $B, G, M > 0$, and the ground state is a conventional crystalline elastic solid. In the orange region, the bulk modulus, $B > 0$, but the pure shear modulus, $G < 0$, while the P-wave modulus, $M = B + G > 0$. We expect the ground state to be fluid and contain highly anisotropic polygons. In the green region, $B < 0$, but $G, M > 0$. In the red region, $G > 0$, but $B, M < 0$. In this region, the P-wave speed is imaginary, and hence, the tissue must undergo a contractile instability of the homogeneous crystalline state and generate regions of high and low densities, which maybe identified with actin hotspots and coldspots. The white markers with black, red, and green borders mark the maximum $\hat{A}$ values at which $G = 0$, $M = 0$, and $B = 0$, respectively.

#### II. GROUND STATE PHASE DIAGRAM OF THE VERTEX MODEL WITH MCFL-I

##### A. Vertex Model

We use the following vertex model with mechanochemical feedback.

$$E = \frac{1}{2}\left[K(A - A_0)^2 + \Gamma(P - P_0)^2\right] \tag{S40}$$

$$A_0(A) = 2a_0 p_b(A) \tag{S41}$$

$$P_0(A) = 2\hat{q}_0\sqrt{a_0}[1 - p_b(A)] \tag{S42}$$

$$p_b(A) = \frac{1}{1 + \left(\frac{A}{\hat{A}}\right)^2} \tag{S43}$$

##### B. Elastic moduli of the hexagonal ground state

Following the procedure described in [10], we can calculate the pure shear and the bulk moduli of the hexagonal crystal. Applying this approach to our model, we get:

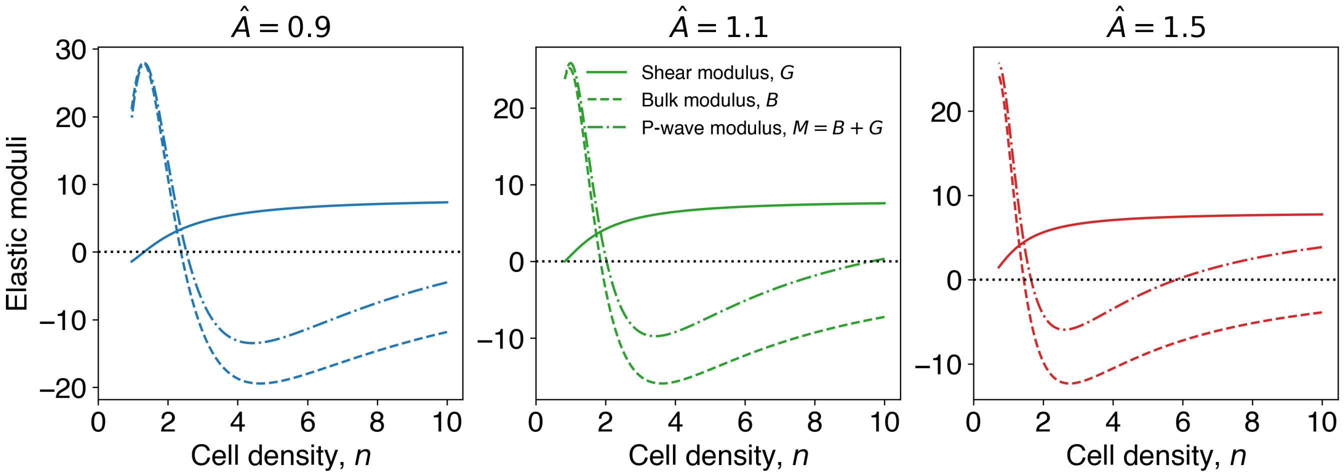


FIG. T8. The elastic moduli of the hexagonal ground state for different $\hat{A}$. At the lowest permissible density, $n_{min}$, the shear moduli can have negative, zero, or positive values depending on $\hat{A}$. These three cases correspond to an elastic fluid, fluid, and a solid phase.

*a. Shear Modulus*

$$G = \sqrt{3}\Gamma\left(12 - \frac{2}{\sqrt{\frac{2A}{3\sqrt{3}}}}P_0(A)\right) \tag{S44}$$

This expression is slightly different from what was obtained in [2, 10], because of the slightly different form of the energy. There, the following expression is obtained:

$$G_F = \sqrt{3}\Gamma\left(12 - \frac{P_0}{\sqrt{\frac{2A}{3\sqrt{3}}}}\right) \tag{S45}$$

The key difference between $G$ and $G_F$ is that the former can be negative, but the latter is always positive. In fact, in the pioneering work of Bi et.al. [11], $G$ was used to mark the boundary between the solid and the liquid phase.

*b. Bulk Modulus* The expression for the bulk modulus, $B$, is cumbersome due to the $A$-dependence of $A_0$ and $P_0$. The expression also does not provide any intuition. Hence, here we provide a sympy code to calculate the expression, and show the results. The key surprise is that the bulk modulus can become negative, while the shear modulus remains positive. In a range of density, the P-wave modulus, $M = B + G$, becomes negative. In this range, no longitudinal (P) waves can propagate, but transverse (S) waves can, which may explain why the amplitude of the actin oscillations dampen with density.

SymPy code for the bulk modulus

```python
### Hexagonal lattice Bulk Moduli
import sympy as sp
import numpy as np

sp.init_printing()
x, A, Ahat, P, Phat, a0, qhat0, K, Gamma,m, a, Lambda, A0, P0, n = sp.symbols('x A Ahat P
↪  Phat a0 qhat0 K Gamma m a Lambda A0 P0 n',positive=True)


### Side length of a hexagon
a = sp.sqrt(2*A/(3*sp.sqrt(3)))
### Box dimensions
# x  = 1+epsilon is the scaling factor. For bulk modulus calculation, both Lx and Ly are
↪  scaled by x.
Lx = x*sp.sqrt(3)*a
Ly = x*3*a
l1 = Ly/3
l2 = sp.sqrt((Lx/2)**2 + (Ly/6)**2)

### Area of a hexagon
Ab = Lx*Ly/2
### Perimeter of a hexagon
Pb = 2*l1 + 4*l2
Sb = (Ab/Ahat)**2
pb = 1/(1+Sb)
A0 = 2*a0*pb
P0 = 2*qhat0*sp.sqrt(a0)*(1-pb)

### There are two hexagons in the box
Ea = K*(Ab - A0)**2

Ep = Gamma*(Pb - P0)**2 ## Canonical vertex
E = (Ea + Ep)

EpF =  Gamma*Pb**2 + Lambda*Pb ### Farhadifar
EF = Ea + EpF


### Bulk modulus from the Vertex Model + MCFL-I
B2 = sp.diff(E,x,2).subs({x:1})/(2*A)

#### Farhadifar bulk modulus
BF = sp.diff(EF,x,2).subs({x:1})/(2*A)

fB2 = sp.lambdify([A, Ahat, P, Phat, a0, q0, K, Gamma],B2,'numpy')


latex_B2 = sp.latex(B2)
sp.simplify(B2,deep=True)
```

##### C. Effect of topological transitions and active forces

The ground state analysis presented here excludes topological transitions, such as cell intercalation through the T1-transitions and the node-switching algorithm used to prevent cell overlaps. Furthermore, we also do not consider the effect of active forces, such as from cell motility and MCFL-II, in these calculations. These transitions and forces can perturb the integrity of the crystalline phase, and we expect them to have nontrivial effects on the stability of the phases. A detailed exploration and characterization of these perturbations is beyond the scope of the current manuscript.

Simulations with just MCFL-I and motility indicate that the motility smoothens out the discontinuous transition. It also indicates that the floppy region below $n_J$ is susceptible to shear deformations.

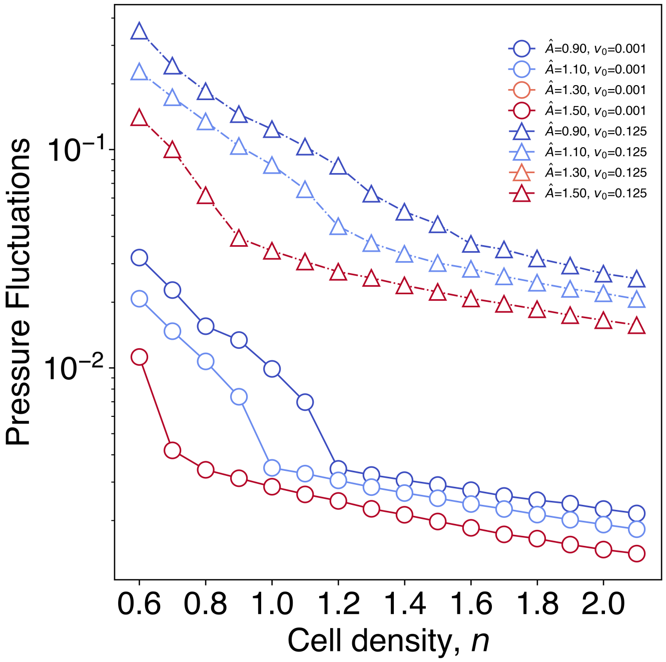


FIG. T9. Pressure fluctuation vs. $n$ for different $\hat{A}$ and motility.

#### III. NUMERICAL SIMULATION OF THE MODEL

To reproduce the diverse behaviors observed experimentally in MDCK cell monolayers, we implement an active vertex model with mechanochemical feedback. An essential feature of the model is that cells can divide within the vertex framework. The mechanochemical feedback consists of two mechanochemical feedback loops (MCFLs). Each component of the model is described below.

##### A. Active Vertex Model

The vertex model represents tissues as a tessellation of polygons, with each polygon representing a cell. The energy of the vertex model for cells labelled by $\alpha$ is written as:

$$E_{VM}(t) = \sum_{\alpha} \left[ \frac{K_\alpha}{2}(A_\alpha - A^0_\alpha(t))^2 + \frac{\Gamma_\alpha}{2}(P_\alpha - P^0_\alpha(t))^2 \right] \tag{S46}$$

Here, the area energy comes from the volume incompressibility of cells whereas the perimeter energy term comes from the actomyosin contractility of cells. The constant parameters $K_\alpha$ is the area modulus which gives the area elasticity and $\Gamma_\alpha$ is the perimeter modulus which is the elasticity due to the actomyosin cortex. The cells can be either solid-like or fluid-like according to the shape index defined as $q = P/\sqrt{A}$.

The vertices in the vertex model undergo overdamped dynamics. So, the equation of motion of the $i$th cell is given by:

$$\zeta \frac{\partial \vec{r}^{(i)}}{\partial t} = \vec{F}^{(i)} + \vec{F}^{(i)}_{act} + \vec{F}^{(i)}_{motility} \tag{S47}$$

Here, $\vec{r}^{(i)}$ is position of the $i$th vertex, and $\vec{F}^{(i)}$ is the equilibrium force on each vertex given by the gradient of the vertex model energy: $\vec{F}^{(i)} = -\nabla_{\vec{r}^{(i)}} E_{VM}$. Additionally, there can be other active forces $\vec{F}^{(i)}_{act}$ and motility force $\vec{F}^{(i)}_{motility}$. We have $\vec{F}_{act} = 0$ in our simulations. $\vec{F}^{(i)}_{motility}$ is a force which propels individual cells by $\vec{F}^{(i)}_{motility} = v_0 \vec{p}_\alpha$, where $\vec{p}_\alpha$ is the polarity of a given cell given by $\vec{p}_\alpha = (\cos\theta_\alpha, \sin\theta_\alpha)$, where this angle $\theta_\alpha$ performs rotational diffusion:

$$\partial_t \theta_\alpha = \sqrt{2D_r}\eta_\alpha(t) \tag{S48}$$

where $\eta_\alpha(t)$ is a Gaussian white noise with zero mean and correlation $\langle \eta_\alpha(t)\eta_{\alpha'}(t')\rangle = \delta(t-t')\delta_{\alpha\alpha'}$.

#### B. Mechanochemical Feedback

##### 1. MCFL-I

The first mechanochemical feedback loop arises from the load-dependent binding of myosin to actin. When a cell is compressed, i.e., when the cell area $A$ decreases, then the unbinding rate of myosin decreases, which changes the preferred area $A_0$ and the preferred perimeter $P_0$ of the cell in the vertex model. In our model, the preferred area and perimeter undergo changes in an area-dependent manner. As mentioned in the main text, actin reorganizes and concentrates at cell-cell junctions as the tissue matures. This means that junctional tension should increase, and the contractility should decrease as a tissue matures. Such an effect must be accounted for by one positive and one negative MCFL. Since junctional tension is given by $-P_0/A_0^{3/2}$ and the contractility is given by $A_0^{-1}$ [1], the feedback should be such that $P_0$ decreases with areal compression and $A_0$ should increase. Such feedback can be accounted for in the following way:

$$\begin{aligned} \tau_A \dot{A}_0 &= -[A_0 - \hat{A}_0(A)] \\ \tau_P \dot{P}_0 &= -[P_0 - \hat{P}_0(A)] \end{aligned} \tag{S49}$$

where we see that the vertex model parameters $A_0$ and $P_0$ are time-dependent and they relax to area-dependent values $\hat{A}_0(A)$ and $\hat{P}_0(A)$. The $\hat{A}_0$ and $\hat{P}_0$ depend on the binding probability of myosin to actin. If the myosin-binding probability is given by $p_{bound} = 1/(1 + (A/\hat{A})^2)$. Here, $\hat{A}$ is related to the dissociation constant $K_d$ of myosin. Then, we know the constitutive relations from before:

$$\begin{aligned} \hat{A}_0 &= 2a_0 p_{bound} \\ \hat{P}_0 &= 2\hat{q}_0\sqrt{a_0}(1 - p_{bound}) \end{aligned} \tag{S50}$$

##### 2. MCFL-II

To model the actin oscillations observed in experimental MDCK monolayers, we model ERK as a Hopf oscillator, which, in this case, is a Brusselator. Other Hopf oscillators also exhibit similar collective behaviours, as we have checked using the FitzHugh-Nagumo model [12]. The chemistry of the Hopf oscillator is coupled to the mechanics via areal compressions, just like in MCFL-I. The 1D version of the model has been studied before to model for ERK oscillations [12, 13]. The above Eq. S49 will now be modified as:

$$\begin{aligned} \dot{M} &= a - (b+1)M + cM^2E \\ \dot{E} &= bM - cM^2E - DE \\ \tau_D \dot{D} &= -(D - D_0) - \beta D(A - 1) \\ \tau_A \dot{A}_0 &= -[A_0 - \hat{A}_0(A)] - \alpha(E - E_0) \\ \tau_P \dot{P}_0 &= -[P_0 - \hat{P}_0(A)] - \frac{\alpha}{2\sqrt{A_0}}(E - E_0) \end{aligned} \tag{S51}$$

where $E$ is ERK, $M$ is MEK, and $D$ is a degrader of ERK, which couples the mechanics to the chemistry via the term $\beta D(A-1)$. The chemistry is coupled back to the mechanics via the terms $\alpha(E-E_0)$ and $\frac{\alpha}{2\sqrt{A_0}}(E-E_0)$.

##### 3. *Cell Divisions*

To introduce cell division in our model, we assume each cell has a cyclin level, $c$. The cyclin increases with time according to the following growth law:

$$\frac{\partial c}{\partial t} = 1 \tag{S52}$$

A cell is picked with a probability rate $r$, and only if $c > c_{th}$ and $A > A_{th}$, where $c_{th}$ is a threshold cyclin value and $A_{th}$ is a threshold area, the picked cell is divided. The selected cell is divided by a line perpendicular to its long axis.

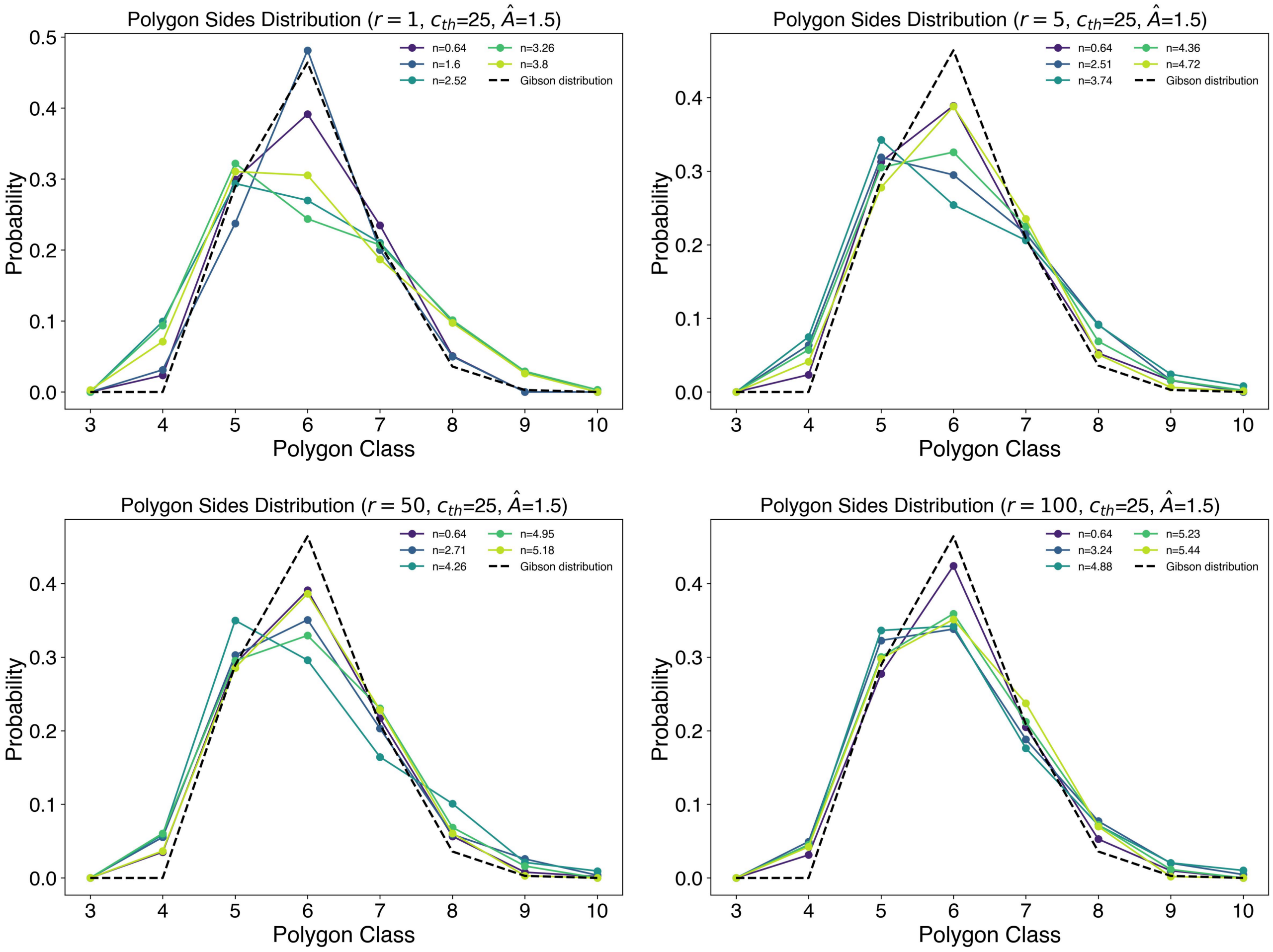


FIG. T10. The polygon distribution at different densities for different $r$ values, compared to the distribution given by Gibson et al.

#### IV. ANALYSIS OF THE EXPERIMENTAL AND SIMULATION DATA

##### A. Ageing

To show ageing in simulation, we have calculated the overlap function $Q(\Delta t)$ and self-intermediate scattering function $F_s(k, \Delta t)$, for cell trajectories with different time origins $t_{origin}$. The $Q(\Delta t)$ plot is shown in Fig. T11 and the $F_s(k, \Delta t)$ plot is shown in Fig. T13.

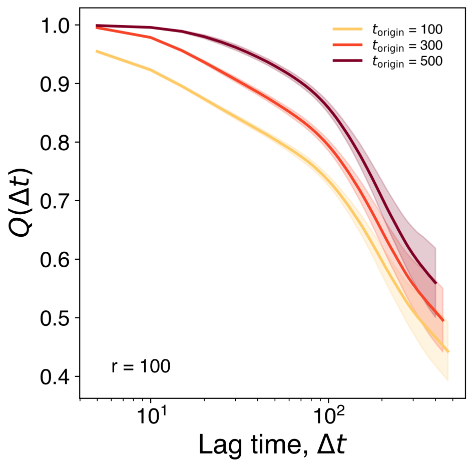


FIG. T11. $Q(\Delta t)$ for different $t_{\text{origin}}$, for $r = 100 \times 10^{-5}$ and $\hat{A} = 1.50$.

To calculate the self-intermediate scattering function, we need to pick a wavevector, $k$. We use the $k$ at which the static structure function $S(k)$ peaks [Fig. T12].

#### B. Force-moment tensor of a cell

Forces in the vertex model act on the vertices of a cell, $c$. If the position vector of vertex $i$ is $\vec{r}_i$ and the force acting on it is $\vec{f}_i$, then the force moment tensor, $\Sigma$, is defined as:

$$\Sigma = \sum_{i \in V(c)} \vec{r}_i \otimes \vec{f}_i \tag{S53}$$

$$\therefore \Sigma_{mn} = \sum_{i \in V(c)} r_{i,m} f_{i,n}. \tag{S54}$$

Here, $V(c)$ is the set of vertices belonging to the cell $c$, and $m, n$ are the components of the vectors. The force-moment tensor serves as the analog of the stress tensor for a single cell. The trace (sum of the eigenvalues) of $\Sigma$ measures the compressive stress at the cell, whereas the difference of the eigenvalues measures the shear stress. In thermal

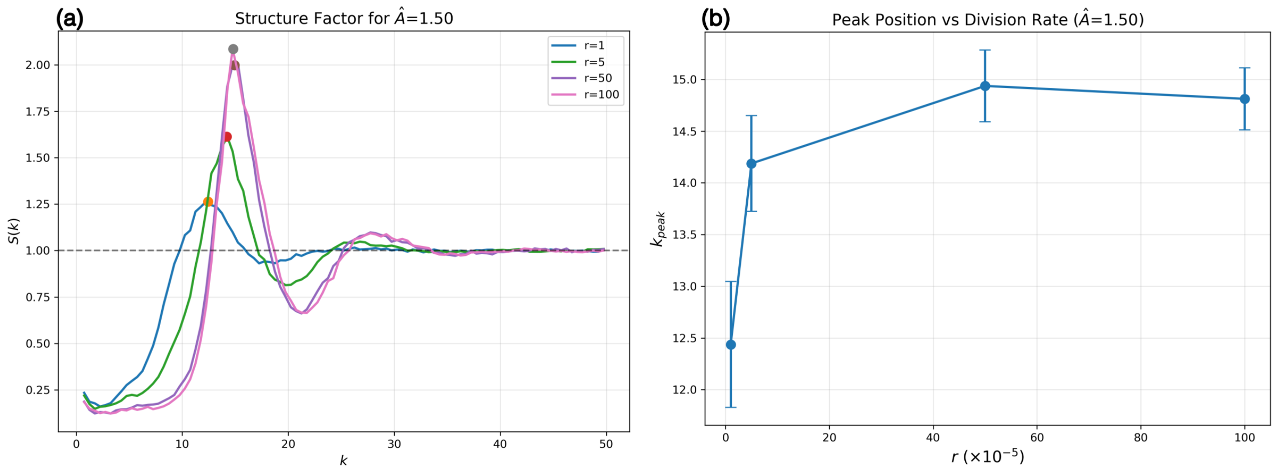


FIG. T12. Time-averaged structured factor for $\hat{A} = 1.50$.(b) $k$ values corresponding to the peaks of $S(k)$.

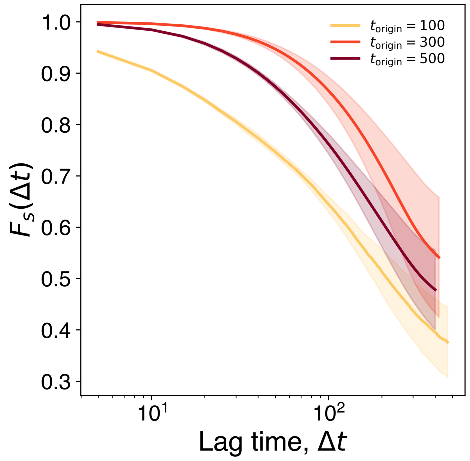


FIG. T13. $F_s(k, \Delta t)$ for different $t_{origin}$, for $k =$, $r = 100 \times 10^{-5}$ and $\hat{A} = 1.50$.

equilibrium, the standard deviation of $Tr.\Sigma$ is inversely proportional to the bulk viscosity, and the standard deviation of the shear stress is inversely proportional to the shear viscosity. In out-of-equilibrium systems, such as the vertex model in this paper, fluctuations and viscosities are inversely related, but this relationship does not arise from thermal equilibrium.

##### C. Asphericity of cell shapes

We quantify cell shape anisotropy using asphericity. To compute asphericity, we define the shape-moment tensor:

$$M = \sum_{i \in V(c)} \vec{r}_i \otimes \vec{r}_i \tag{S55}$$

$$\therefore M_{mn} = \sum_{i \in V(c)} r_{i,m} r_{i,n}. \tag{S56}$$

Next, we compute the eigenvalues of this matrix. In 2D, these are $\lambda_1$ and $\lambda_2$. The asphericity, $\kappa$, is defined as:

$$\kappa = \left| \frac{\lambda_1 - \lambda_2}{\lambda_1 + \lambda_2} \right|. \tag{S57}$$

For a nearly isotropic shape, such as a circle or a regular hexagon, $\kappa \to 0$, whereas for anisotropic shapes $\kappa > 0$ and approaches 1 for shapes with high aspect ratios. Geometrically, it is equivalent to fitting an ellipse to the underlying shape, whose semimajor and semiminor axis lengths are $\lambda_1$ and $\lambda_2$. We use this equivalence to calculate the asphericity of the MDCK cells in the experiments. Each segmented cell is fitted to an ellipse in TrackMate, from which we compute the asphericities.

The asphericity here is related to the aspect ratio (AR) as follows:

$$\kappa = \left| \frac{\lambda_1 - \lambda_2}{\lambda_1 + \lambda_2} \right| = \left| \frac{\lambda_1/\lambda_2 - 1}{\lambda_1/\lambda_2 + 1} \right| = \left| \frac{AR - 1}{AR + 1} \right|. \tag{S58}$$

where the aspect ratio is given by $AR = \lambda_1/\lambda_2$. The asphericity $\kappa$ is related to the aspect ratio, which has been shown to have a universal distribution for confluent epithelial monolayers [14].

##### D. Dynamic Heterogeneity Size Calculation

First, we grid the system into a $25 \times 25$ grid and average the cell velocities in each box. This gives a velocity map of the system. After this, we find the grid points with the top 20% velocity. Following [15], we classify the dynamic heterogeneity region as the largest contiguous region (here, a collection of boxes) as the dynamic heterogeneity region and the size of that region as the dynamic heterogeneity size.

##### E. Wavelet Analysis of the signal

We analyzed the spatiotemporal dynamics of both experimental and theoretical datasets using a unified framework. In the experimental system, actin-labeled images were acquired continuously over a 30-hour period with a temporal resolution of 30 minutes, yielding a sequence of spatially resolved intensity maps. In parallel, from simulations of our mechanochemical feedback model, we obtained the temporal evolution of the projected inverse area over the same time window. To make the two datasets directly comparable, the field of view in both cases was divided into a uniform $20 \times 20$ grid. Within each grid element, local time series data were extracted by averaging pixel intensities from actin images in the experiment, and inverse area images from the simulations.

A central consideration in this comparison is the correspondence between actin intensity in experiment and inverse area in simulation. Experimentally, actin accumulation reflects structural compaction: higher actin levels are associated with smaller areas. Accordingly, we treated inverse area from simulations of our model as a proxy for actin concentration in the experimental data. This mapping enabled a consistent interpretation across modalities, thereby allowing theoretical predictions to be grounded in experimentally measurable quantities.

To isolate oscillatory components, the raw time series from both experiment and theory were first detrended using the PyBOAT software package. PyBOAT (A Biological Oscillations Analysis Toolkit) uses smoothing splines to remove low-frequency baseline trends, ensuring that slow signal variations do not mask the underlying oscillatory dynamics. After detrending, PyBOAT applies a continuous wavelet transform using the Morlet wavelet, generating a time–frequency representation of the signal. This approach captures transient and non-stationary oscillations that standard Fourier analysis would miss. From the resulting wavelet spectra, we extracted the dominant period and quantified the oscillation power for each grid element and then averaged the power values to obtain a representative measure of oscillatory strength, which was compared across simulated and experimental datasets. In the main text, to compare with the experiment, we choose $r = 5 \times 10^{-5}$, $c_{th} = 25$, and $\hat{A} = 1.50$ from our simulation in wavelet analysis.

#### V. CHOICE OF PARAMETERS

##### A. Choice of $\alpha$ and $\beta$

MCFL-II generates oscillations in the model, which has the mechanochemical couplings $\alpha$ and $\beta$. From simulations, we have found that the time period distributions of oscillations do not change with $\alpha$ and $\beta$ but change with $\tau_D$, $\tau_A$ and $\tau_P$. We have chosen values $\alpha = 2.25$ and $\beta = 2.88$.

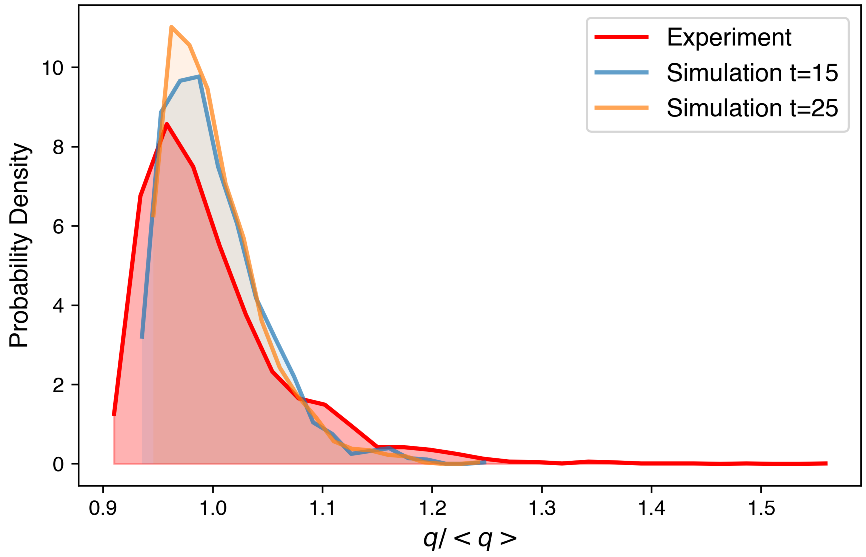


FIG. T14. Comparison of shape index distributions from experiment and simulation for $\alpha = 2.25$ and $\beta = 2.88$.

##### B. Choice of $\tau_D$, $\tau_A$, $\tau_P$

The time period of oscillations generated by MCFL-II depends on $\tau_D$, $\tau_A$ and $\tau_P$. We have chosen the values by comparing the distributions with the experimentally observed distributions.

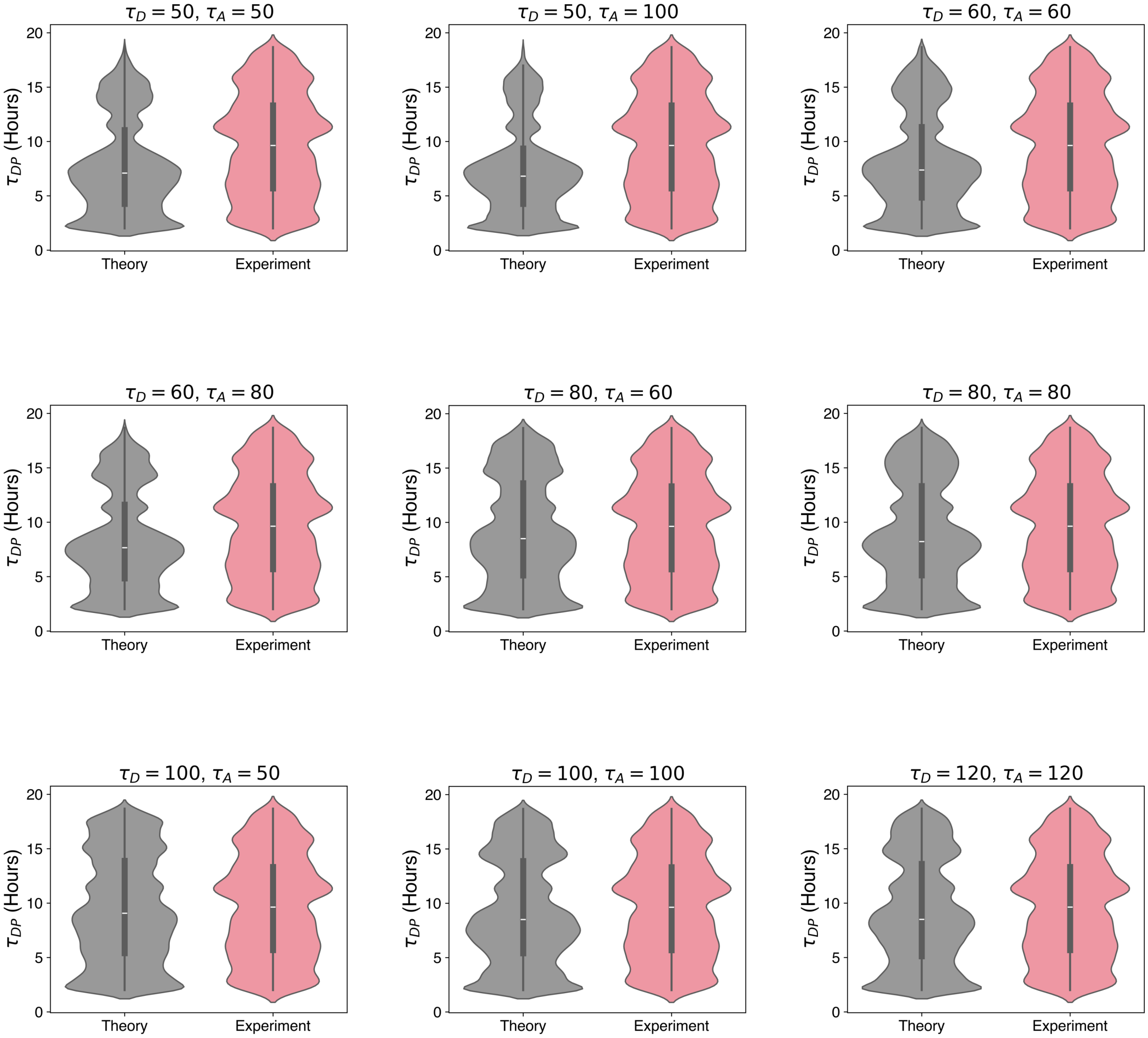


FIG. T15. Violin plots showing the dominant time period distributions compared with those obtained from experimental data. Here, $\tau_A = \tau_P$, $\hat{A} = 1.50$, $r = 50 \times 10^{-5}$.

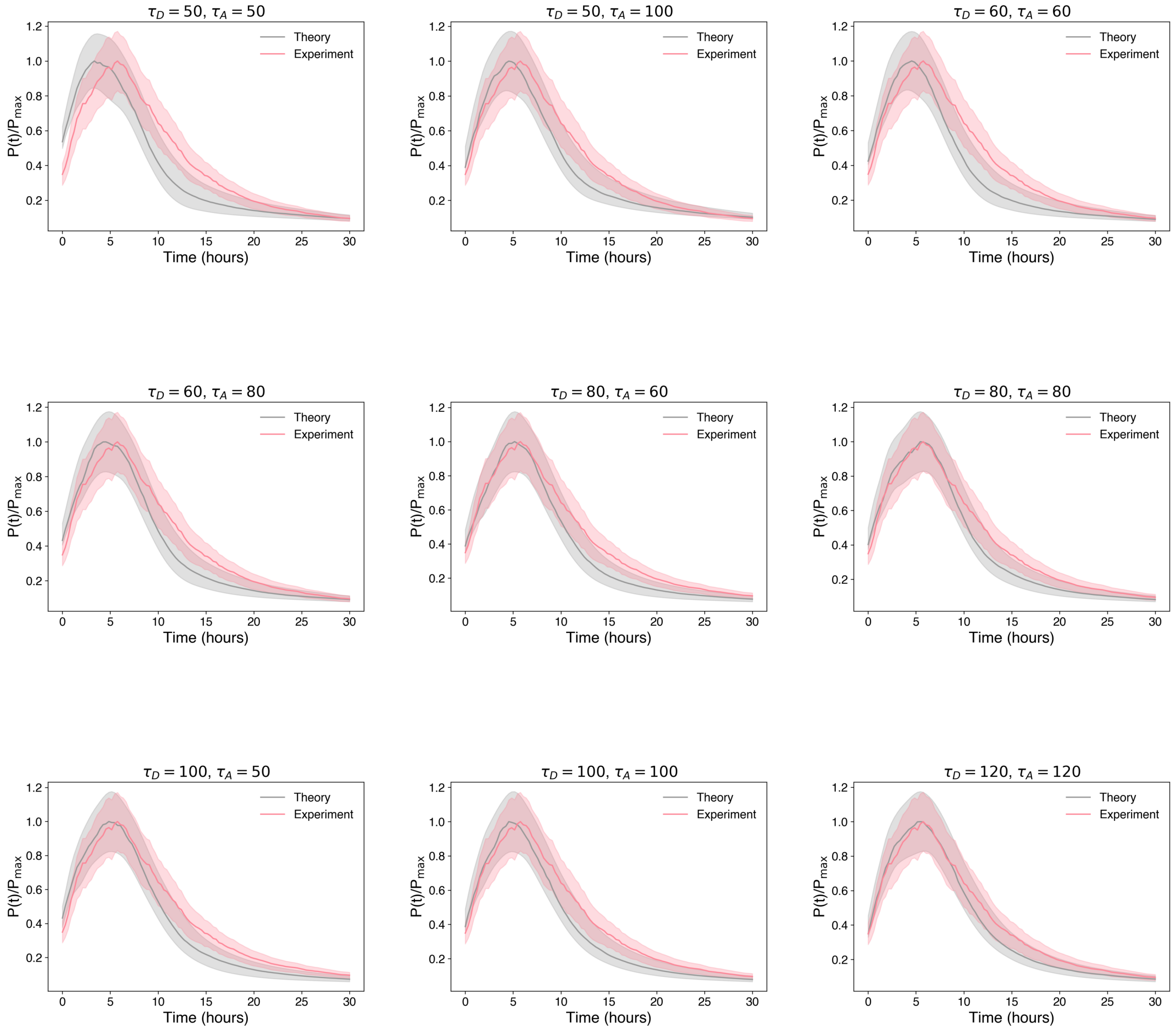


FIG. T16. Power time series plots comparison between experiment and simulation. Here, $\tau_A = \tau_P$, $\hat{A} = 1.50$, $r = 50 \times 10^{-5}$.

By observing that the distributions match best for $\tau_D = \tau_A = \tau_P = 120$, we use those values for the simulations.

#### C. Choice of $r$, $c_{th}$ and $A_{th}$

We have chosen the values of $r$ and $c_{th}$ by comparing the simulation data with the experimentally observed growth rate.

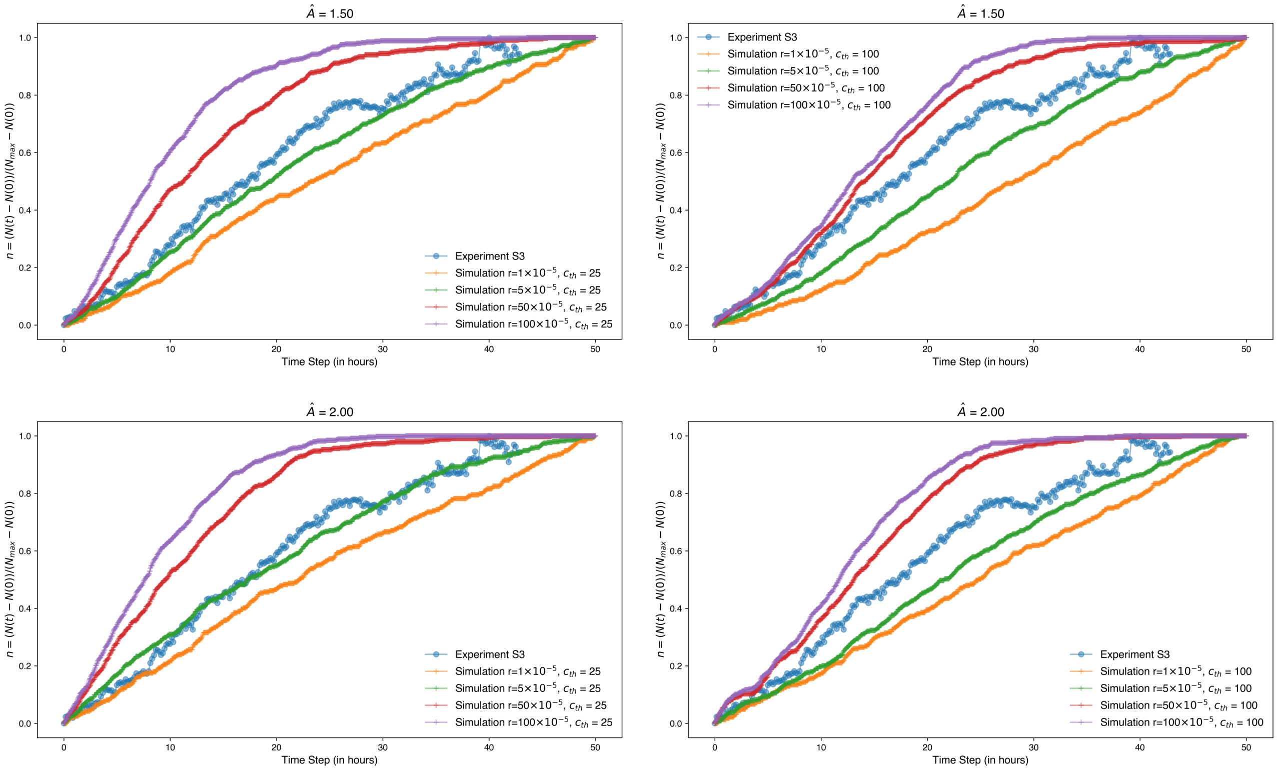


FIG. T17. Cell number growth for different division rates, $r$, for $c_{th} = 25, 100$, and $\hat{A} = 1.5, 2$.

The $N(t)$ curve in simulations plateaus as the cell areas continuously decrease over time [Fig. T17], as seen in the decrease of $dN/dt$ with time [Fig. T18]. We have obtained $\frac{dN}{dt}(t)$ by fitting a function $N(t) = A(1 - e^{-kt}) + N_0$ to the $N(t)$ data and then finding $\frac{dN}{dt} = Ake^{-kt}$. The time at which the $N(t)$ starts to plateau depends on the value of $A_{th}$. The value for simulations is set to $A_{th} = 0.3$.

Moreover, we see that the time period distribution fits better for the $r \approx 5 \times 10^{-5}$, which match the experimental growth rate [Fig. T19].

The growth rate $dN/dt$ in simulations varies with the parameters $r$ and $c_{th}$ as given in Fig. T20.

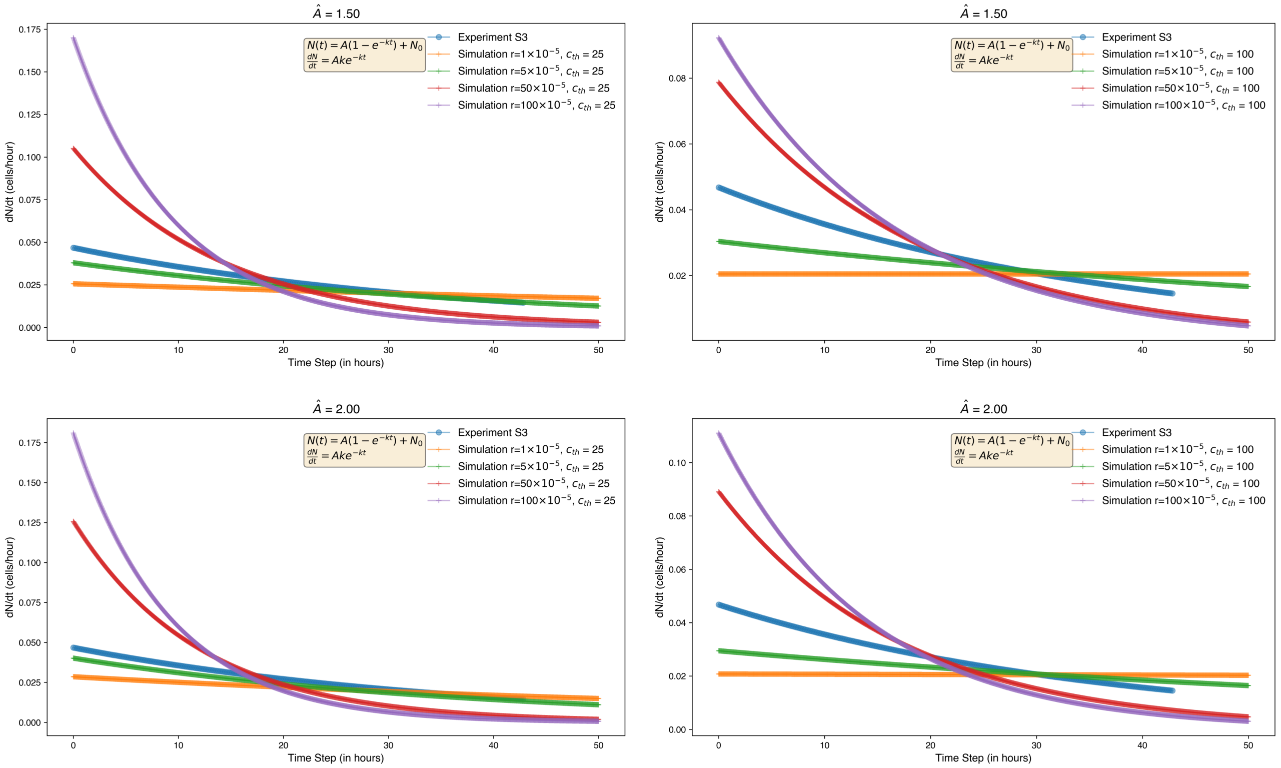


FIG. T18. Cell number growth rate for different division rates, $r$, for $c_{th} = 25, 100$, and $\hat{A} = 1.5, 2$.

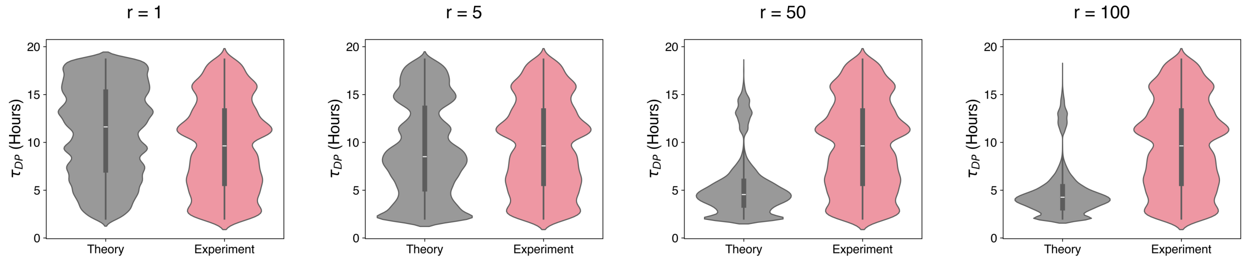


FIG. T19. Comparison between the dominant time period distributions from simulation and experiment for different $r$ (in $\times 10^{-5}$), $\hat{A} = 1.5$, and $c_{th} = 25$

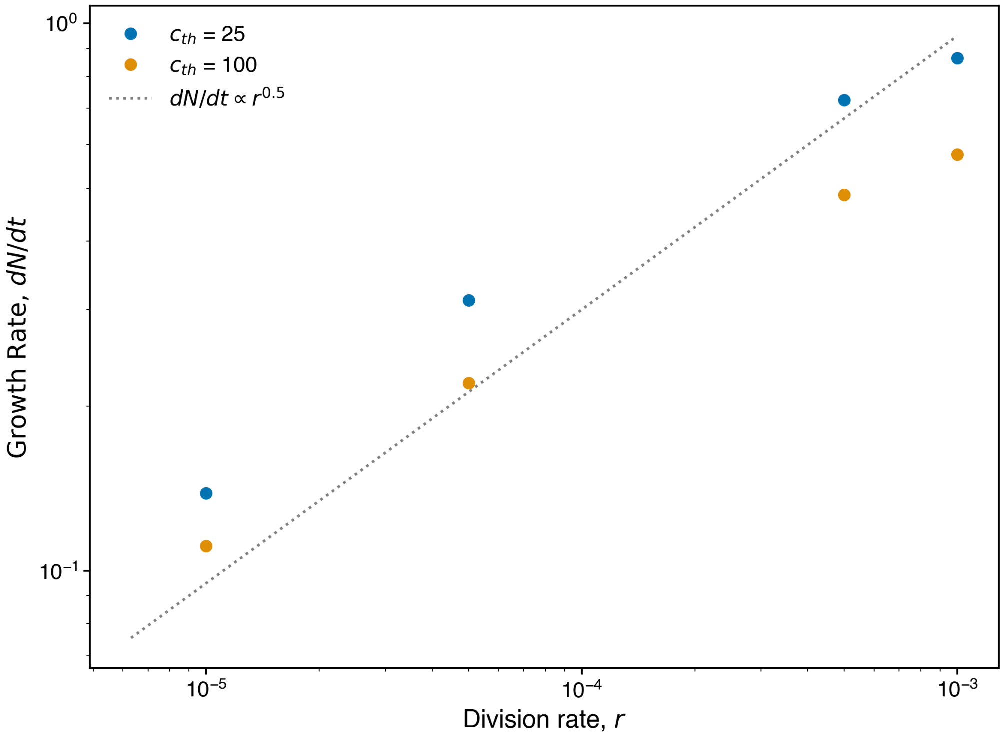


FIG. T20. Growth rates for different $r$ and $c_{th}$.

The growth rate here is calculated by fitting a straight line to the initial increase in the $N(t)$ data from simulations.

#### D. Parameters Used

| Parameter | Value |
|---|---|
| $K$ | 1.2 [16] |
| $\Gamma$ | 0.38 [16] |
| $\zeta$ | 1 |
| $\hat{q}_0$ | 3.9 |
| $v_0$ | 0.125 |
| $D_r$ | 0.5 |
| $\tau_D = \tau_A = \tau_P$ | 120 |
| $\Delta t$ | 0.01 |

- **Simulation and Experimental timescales:** 1 simulation time step = 2 minutes in experiment.
- The values of other parameters are mentioned in the caption of the main text figures.
- **Topological transitions:** The simulation has the following topological transitions:
  - T1 transitions
  - T2 transitions: cell divisions
  - A node switch operation has also been implemented to prevent overlaps [17].

[S1] R. Farhadifar, J.-C. Röper, B. Aigouy, S. Eaton, and F. Jülicher, The influence of cell mechanics, cell-cell interactions, and proliferation on epithelial packing, Current Biology **17**, 2095–2104 (2007).

[S2] D. B. Staple, R. Farhadifar, J. C. Röper, B. Aigouy, S. Eaton, and F. Jülicher, Mechanics and remodelling of cell packings in epithelia, The European Physical Journal E **33**, 117–127 (2010).

[S3] N. K. Noren, W. T. Arthur, and K. Burridge, Cadherin engagement inhibits rhoa via p190rhogap, Journal of Biological Chemistry **278**, 13615–13618 (2003).

[S4] I. Molina-Ortiz, R. A. Bartolomé, P. Hernández-Varas, G. P. Colo, and J. Teixidó, Overexpression of e-cadherin on melanoma cells inhibits chemokine-promoted invasion involving p190rhogap/p120ctn-dependent inactivation of rhoa, Journal of Biological Chemistry **284**, 15147–15157 (2009).

[S5] R. Priya, A. S. Yap, and G. A. Gomez, E-cadherin supports steady-state rho signaling at the epithelial zonula adherens, Differentiation **86**, 133–140 (2013).

[S6] M. Abhishek, A. Dhanuka, D. S. Banerjee, and M. Rao, Excitability and travelling waves in renewable active matter, arXiv preprint arXiv:2503.19687 (2025).

[S7] R. Sknepnek, I. Djafer-Cherif, M. Chuai, C. Weijer, and S. Henkes, Generating active t1 transitions through mechanochemical feedback, Elife **12**, e79862 (2023).

[S8] A. Saraswathibhatla and J. Notbohm, Tractions and stress fibers control cell shape and rearrangements in collective cell migration, Phys. Rev. X **10**, 011016 (2020).

[S9] J.-A. Park, J. H. Kim, D. Bi, J. A. Mitchel, N. T. Qazvini, K. Tantisira, C. Y. Park, M. McGill, S.-H. Kim, B. Gweon, J. Notbohm, R. Steward Jr, S. Burger, S. H. Randell, A. T. Kho, D. T. Tambe, C. Hardin, S. A. Shore, E. Israel, D. A. Weitz, D. J. Tschumperlin, E. P. Henske, S. T. Weiss, M. L. Manning, J. P. Butler, J. M. Drazen, and J. J. Fredberg, Unjamming and cell shape in the asthmatic airway epithelium, Nature Materials **14**, 1040–1048 (2015).

[S10] R. Farhadifar, *Dynamics of cell packing and polar order in developing epithelia*, Ph.D. thesis (2009).

[S11] D. Bi, J. Lopez, J. M. Schwarz, and M. L. Manning, A density-independent rigidity transition in biological tissues, Nature Physics **11**, 1074 (2015).

[S12] P. Dewan, S. Mondal, and S. Sarkar, Oscillation death by mechanochemical feedback (2025), arXiv:2504.19655 [cond-mat.soft].

[S13] D. Boocock, N. Hino, N. Ruzickova, T. Hirashima, and E. Hannezo, Theory of mechanochemical patterning and optimal migration in cell monolayers, Nature Physics **17**, 267–274 (2020).

[S14] S. Sadhukhan and S. K. Nandi, On the origin of universal cell shape variability in confluent epithelial monolayers, eLife **11**, e76406 (2022).

[S15] T. E. Angelini, E. Hannezo, X. Trepat, M. Marquez, J. J. Fredberg, and D. A. Weitz, Glass-like dynamics of collective cell migration, Proceedings of the National Academy of Sciences **108**, 4714 (2011), https://www.pnas.org/doi/pdf/10.1073/pnas.1010059108.

[S16] D. Boocock, T. Hirashima, and E. Hannezo, Interplay between mechanochemical patterning and glassy dynamics in cellular monolayers, PRX Life **1**, 013001 (2023).

[S17] A. G. Fletcher, J. M. Osborne, P. K. Maini, and D. J. Gavaghan, Implementing vertex dynamics models of cell populations in biology within a consistent computational framework, Progress in Biophysics and Molecular Biology **113**, 299 (2013).
